# Optimizing DNA origami assembly through selection of scaffold sequences that minimise off-target interactions

**DOI:** 10.1101/2025.01.29.635450

**Authors:** Ben Shirt-Ediss, Emanuela Torelli, Silvia Adriana Navarro, Hadeel Khamis, Ariel Kaplan, William Trewby, Juan Elezgaray, Nima Moradzadeh-Esmaeili, Michael Haydell, Daniel Keppner, Michael Famulok, Natalio Krasnogor

## Abstract

DNA origami is a mainstay of DNA nanotechnology and several efforts have been devoted to understanding how various factors of the self-assembly reaction affect the final yield of the target origami structure. This study analyses how base sequence affects origami yield through the generation of off-target side reactions during self-assembly. Off-target bindings are an under-explored phenomenon and can potentially introduce unwanted assembly barriers and kinetic traps in the origami folding pathway. We developed a multi-objective computational approach that takes a given origami design and scores different scaffold sequences (and their complementary staples) for the prevalence of four different types of off-target binding events. Using our method on DNA origami, we can select ‘bad’ regions of biological sequences (like lambda DNA phage) that, when used as origami scaffold sequences, have an excessive number of off-target side reactions for each shape. We show, using high-resolution atomic force microscopy (AFM), that these scaffold sequences largely fail to fold into the target triangle or rectangle structure *in vitro*, despite the scaffold sequence having a fully complementary staple set present. Conversely, using our method we can also select ‘good’ regions of biological sequences. These sequences are deficient in off-target reactions and when used as origami scaffolds, fold more successfully into their target structures as characterised by AFM.

These results have been validated in “blind” folding experiments at two different laboratories in which the experimenters did not know which scaffolds were good or bad folders. To further investigate assembly behaviour, optical tweezers experiments revealed distinct mechanical response profiles, correlating with scaffold-specific off-target interactions. While variants with higher GC content show a high mean unfolding force, variants with lower off-target binding demonstrated more uniform force-extension curves. Our analysis confirmed that high off-target binding leads to increased structural heterogeneity, as seen in the clustering behaviour of unfolding traces of OT experiments. Overall, our work demonstrates how the off-target reactions implicit in base sequences can derail the origami self-assembly process if sufficiently prevalent, and we provide a software tool to select scaffold sequences that minimise off-target reactions for any DNA origami design.

## INTRODUCTION

The scaffolded DNA (and RNA) origami technique permits fabrication of nano-scale objects at high spatial precision via the programmable self-assembly of nucleic acids^1^. The bio-compatibility and bio-interpretability of origami nano-structures naturally position them for future biomedical applications^2–4^.

Rather than top-down fabrication, DNA origami nanostructures are obtained through a complex ‘bottom-up’ self-assembly reaction involving hundreds of different types of single-stranded nucleic acids interacting in parallel via hybridisation, melting and strand-displacement reactions in a de-creasing temperature gradient applied over an extended period. The result of the cooled self-assembly reaction is, ideally, the target nanostructure at high copy number and yield.

An increasing number of experimental and theoretical/computational studies are being devoted to understanding the detailed biophysics of origami self-assembly, both to pin-point which factors are the most important to achieve high-yield origami folding and to understand the surprising robustness of the folding process.

These works have investigated how *in vitro* origami assembly is affected by factors such as the origami design itself^5–8^ (i.e. the staple and scaffold routing patterns), the temperature protocol^9,10^, the stoichiometric ratio of staples to scaffold^7,11,12^, the type and amount of salts in the assembly buffer^10,13^, the solvent^14^ and the primary base sequences of the scaffold and staple strands.

The focus of the current study is on how the latter factor of base sequence can be manipulated to increase origami self-assembly yield. Although base sequences encode the folding information itself and have a combinatorially large space of possibilities, this assembly factor is often just reduced to one number: GC content. Base sequences have traditionally been regarded as providing just an enthalpic contribution to origami folding^15^, where higher nearest-neighbour base energies have been experimentally demonstrated to generally raise the melting temperature of the structure^16,17^ and higher GC content scaffolds have been shown to assemble at lower salt concentrations^16^.

However, more nuanced effects of base sequence on self-assembly are emerging. For example, in a recent study, scaffold sequence has been demonstrated to be able to control the folding pathway of specialised origami designs that have two distinct possible end states^18^ by specifying the locations of folding nucleation points at the early high-temperature phase of folding.

Another nuanced but potentially important effect of the base sequence is the long-range effects of off-target side reactions (i.e. partial mis-bindings) that particular sequences make possible throughout the whole system of hundreds of assembling strands. This was first touched on by Rothemund^19^ who avoided base positions 5515-5587 of the M13 scaffold in his original origami designs because it contained a strong hairpin loop. Since then, some origami CAD tools have included design algorithms to make synthetic scaffold sequences deficient in repeated regions and with controlled GC content^20^. In a similar vein, 4-letter de Bruijn sequences^21^ have also been suggested as suitable synthetic scaffold sequences due to their ability to repress all repeats shorter than a certain number of bases, thus, in principle, providing ‘unique addressability’ for binding staples^16,22,23^.

By and large, off-target reactions in DNA origami assembly have so far been ignored by the DNA nanotechnology community because of the perceived robustness of the DNA origami folding process^15,24^. Origami assembly is largely conceived as a ‘sequence-agnostic nucleation and growth system’^12^. As observed by atomic force microscopy (AFM)^10,25^ it likely involves the formation of nucleation centres at high temperature where on-target staples bind and make short loops in the scaffold strand. Bound staples then begin to ‘organise the scaffold’^19^ and co-operatively help the binding of immediately adjacent on-target staples, so efficient folding fronts rapidly extend from the initial nucleation sites. Folding from high temperature also helps to ensure that on-target staple bindings form before weaker off-target bindings have any opportunity, and, if off-target bindings are present, they are assumed to be corrected by strand displacement from invading on-target staples^19^.

However, as robust as self-assembly seems, it is conceivable that the nucleation-and-growth process could still get kinetically trapped (for potentially long times) away from the target equilibrium origami structure if base sequences made particular off-target bindings possible. For example, a strong scaffold secondary structure made more favourable because of the high effective concentration of the scaffold strand with respect to itself, could block on-target staple binding sites and instead could co-operatively assist the mis-binding of staples. In this way, via unplanned off-target bindings, the sequence could make entropic (via unintended loop formation) as well as enthalpic contributions to folding. Furthermore, the ability of on-target staples to strand-displace mis-bound configurations could be hampered by off-target staples reversibly binding and occluding toeholds. Staples with a leg misbound could conceivably permit staple blocking^12,26^ by other copies of the same staple. Finally, off-target bindings between the staples themselves could create secondary structure slowing the kinetic rate of on-target staple attachment to the scaffold.

Off-target reactions could be considered as more relevant when scaffold sequences are derived from repeat-prone biological sources^27^, when staple legs are fairly short (7 or 8nt) and bind initially at intermediate temperatures, and when constant-temperature folding with no initial thermal denaturation step is employed^10,28,29^.

Definitive answers about when the ‘noise’ of off-target reactions is sufficient to derail the nucleation-and-growth assembly process for specific origami designs with specific scaffold sequences could seemingly be obtained from theoretical/computational models of origami folding.

Simulation of DNA origami folding by coarse grain molecular dynamics^26^ does in principle rigorously include all off-target reaction possibilities because of the base-level resolution. However, such simulations are practically restricted to second timescales after months of computation and typically require artificial assembly conditions (a small origami, high staple concentrations, relatively high constant temperature) to be feasible. Further coarse-graining to 8nt per bead and using a switchable force field^6^ has more recently allowed to simulate self-assembly for kilobase-size origamis (again at second timescales, constant temperature). However, the latter model only considers hybridising bead pairs to be those in the target origami design; accordingly, kinetic traps are limited to situations where on-target binding sites become sterically inaccessible during folding. Moreover, the 8nt bead resolution cannot incorporate strand-displacement reactions – a crucial mechanism for resolving off-target binding sites.

Approximate ‘domain-level’ kinetic models of origami assembly based on mass action rates between hybridisation states^17,30^ can access experimentally relevant timescales (hours) by sacrificing exact geometric information for a purely topological hybridisation state. They are also able to include a full-temperature annealing ramp in the simulation. However, again, these models only consider on-target staple-scaffold bindings in their transition possibilities, leaving staple blocking (at very high staple concentrations) as the only mechanism able to create kinetic traps. While off-target reactions and associated strand displacement mechanisms could be (partially) included in these mass action models in future by relaxing the domain-level constraint, it is a formidable undertaking to implement: accurate reaction enumeration and parameterisation of possible hybridisation and (non-standard) strand-displacement reactions is non-trivial, and difficult computational problems such as stiff dynamics and efficient shortest path calculation on complex graphs must be overcome. Furthermore, all theoretical models mentioned above only track a single origami instance and do not take into account e.g. possible undesired inter-origami binding effects (like hybridisations between scaffold strands).

Given the difficulty in completely disregarding the importance of off-target reactions in DNA origami self-assembly, and the challenges of rigorously incorporating off-target side reactions into mechanistic simulation models, in this work, we develop a different ‘static’ optimization approach to identify origami scaffold sequences that fold reliably for a particular origami design (which we call “reliable folders”, or r-folders for short).

Our computational approach is based on exactly enumerating and minimising four types of off-target side reactions for a particular origami design and uses multi-objective optimisation techniques (for example, also recently applied to RNAs^31^, proteins^32^ and gene circuits^33^) to identify a ‘Pareto set’ of scaffold sequences that best minimise off-target reactions from a large pool of alternatives. With a staple set that leads to a reduced number of off-target reactions, these scaffold sequences (and their complementary staple set) are statistically less likely to get kinetically trapped during the origami folding process.

In the remainder of the paper, we describe our computational method and demonstrate *in vitro* using a DNA triangle and rectangle that selecting scaffold sequences with a high number of associated off-target reactions leads to failed origami folding (even when all staples are Watson-Crick complements to the scaffold). Conversely, we demonstrate scaffold sequences with fewer off-target reactions assembled with higher yield for both origamis. Besides AFM verification of our computational predictions, we also conduct detailed single-molecule optical tweezer (OT) experiments on the various designs with results that align with and further confirm our calculations.

## RESULTS

### Principles of multi-objective scaffold sequence selection

Our multi-objective approach to selecting reliable folder origami scaffold sequences is summarised in Figure 1. A target origami design is specified *a priori* and a pool of ‘sequence variants’ (potential scaffold sequences, each with their Watson-Crick complementary staples) is scored on four thermodynamic objective functions, which we also call “metrics”. Each of these metrics *M*_1_ to *M*_4_ characterise the prevalence of a particular class of off-target binding site during folding. Objective (or metric) *M*_1_ represents the total energy of staple-scaffold off-target binding sites; *M*_2_ the total energy of scaffold-scaffold binding sites (all off-target); *M*_3_ the energy of the worst staple-staple co-fold and *M*_4_ the energy of the worst staple hairpin (see Supplementary Note 2 for detailed definitions). When an objective function is minimised to zero it signifies that off-target bindings of that particular type do not exist.

**FIG. 1.**
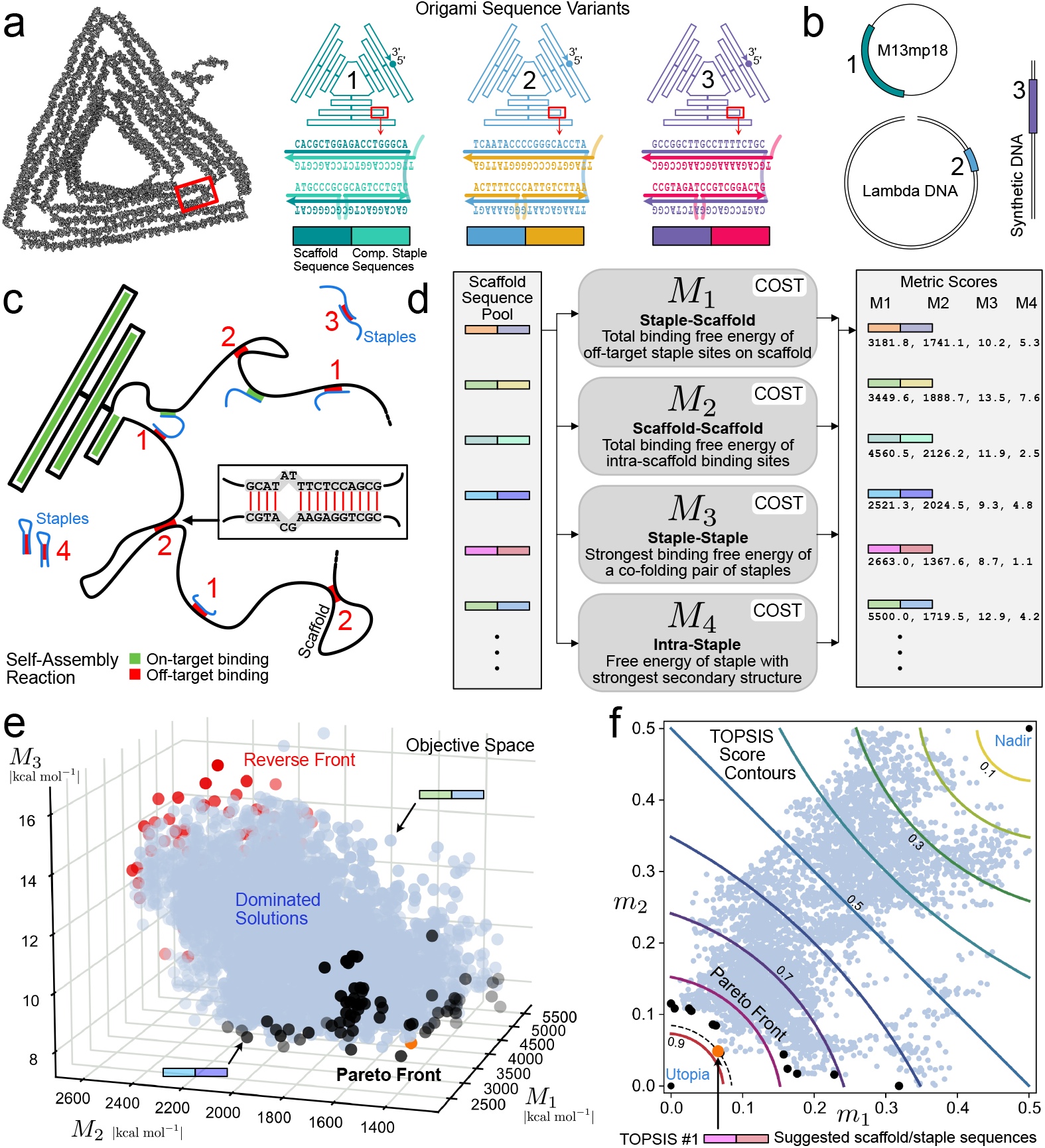
Principles of multi-objective scaffold sequence selection. **(a)** The same DNA origami target design can be realised by many different scaffold sequences. Three possible sequence variants are displayed for a triangle origami. Each variant, denoted as a pair of coloured blocks, consists of a unique scaffold sequence along with the Watson-Crick complementary staple sequences that pin the scaffold strand into a specific rotation. **(b)** Scaffold sequences are commonly sourced from regions of biological vectors, custom-cloned sequences or synthetic sequences. **(c)** The origami self-assembly reaction involves numerous off-target bindings (red) in addition to the intended on-target staple-scaffold bindings (green). Off-target bindings are classified into four types 1-4, which in turn are measured by metrics *M*_1_*… M*_4_. **(d)** Scaffold sequence selection takes place by scoring a large pool of scaffold/staple sequence variants on four cost criteria that quantify the extent of each off-target binding site type in panel (c). Full definition of each metric in Supplementary Note 2. **(e)** After scoring, all variants (here *n* = 5000) are mapped to a low-dimensional objective space (*M*_4_ scores are omitted for 3D visualisation). The Pareto front of optimal trade-off variants is computed in this space (black dots). **(f)** A multi-criteria decision-making method (MCDM; here TOPSIS) ranks Pareto candidates and aids a human decision maker in choosing a single scaffold/staple sequence variant to order for *in vitro* assembly. To visualise TOPSIS ranking, TOPSIS score contours are shown on a 2-dimensional objective space: this method works similarly on a 3 or 4-dimensional objective space. See Supplementary Note 3.

Rather than making a direct prediction of origami assembly yield for a sequence variant, the premise of our method is instead to calculate four scores for each variant that are general heuristics for high-yield folding (but not absolute measures of it). Our assertion is that sequence variants with low-scoring objectives are statistically more likely to fold with high-yield than sequence variants with high-scoring objectives because fewer off-target sites correlate with reduced total kinetic traps on the origami folding pathway. That is, low scores amongst (most) metrics help identify r-folders.

The relative comparison of sequence variants takes place in objective space (Figure 1e) where a set (called the ‘Pareto front’) of optimal trade-off sequence variants becomes defined (black dots). Sequence variants on the Pareto front are special in that they represent the set of optimal trade-offs for the optimisation problem: for Pareto variants, it is not possible to decrease the prevalence of one type of off-target reaction without (potentially) increasing the prevalence of other types of off-target reaction. A Pareto front exists because it is typically impossible for a sequence variant to minimise all metrics *M*_1_ to *M*_4_ to zero simultaneously. Similarly, a reverse Pareto front of ‘worse’ variants can be defined (red dots, Figure 1e) by treating the scoring metrics as benefits to be maximised rather than costs to be minimised.

For a given origami design and pool of scaffold/staple sequence variants realising that design, our method returns the Pareto set of sequence variants with the best trade-offs at minimising off-target reactions during self-assembly. Therefore, not one sequence variant, but a cloud of potentially good sequence variants are returned. Using all four scoring metrics, the Pareto set is typically 2-3% of the initial pool of sequence variants. When the initial sequence variant pool has a size of 5000, the Pareto set is over 100 variants. This number of Pareto sequence variants is still too many for a human decision-maker to manually review and choose between. Therefore, for the last step, we use a Multi-Criteria Decision Making (MCDM) method^34^ to help decide which single variant on the Pareto front is the most appropriate scaffold/staple sequence set to order for implementing the origami design (see Supplementary Note 3 for full details of SAW, KNEE and TOPSIS methods). For each of the MCDM methods, in the absence of better information, we weighted all of the thermodynamic scoring metrics *M*_1_ to *M*_4_ equally.

Finally, we developed a fast approximated energy model parameterised on NUPACK^35^ for calculating metrics *M*_1_ and *M*_2_ (summarised in Figure 2). The energy model uses sliding window techniques to efficiently detect off-target binding sites between staples and scaffold and to detect regions where the scaffold can bind to itself. Binding sites are detected at baselevel resolution and the energy model allows for mismatching symmetric interior loops of different sizes within binding sites (see Supplementary Note 1). Of note, the energy model takes into account all potential scaffold-scaffold bindings when calculating *M*_2_, not just those bindings present in the minimum free energy (MFE) configuration of the scaffold strand; also it is much faster to calculate than the MFE configuration. For metrics *M*_3_ and *M*_4_ requiring fewer calculations, we call NU-PACK to calculate binding energies based on full sequence information.

**FIG. 2.**
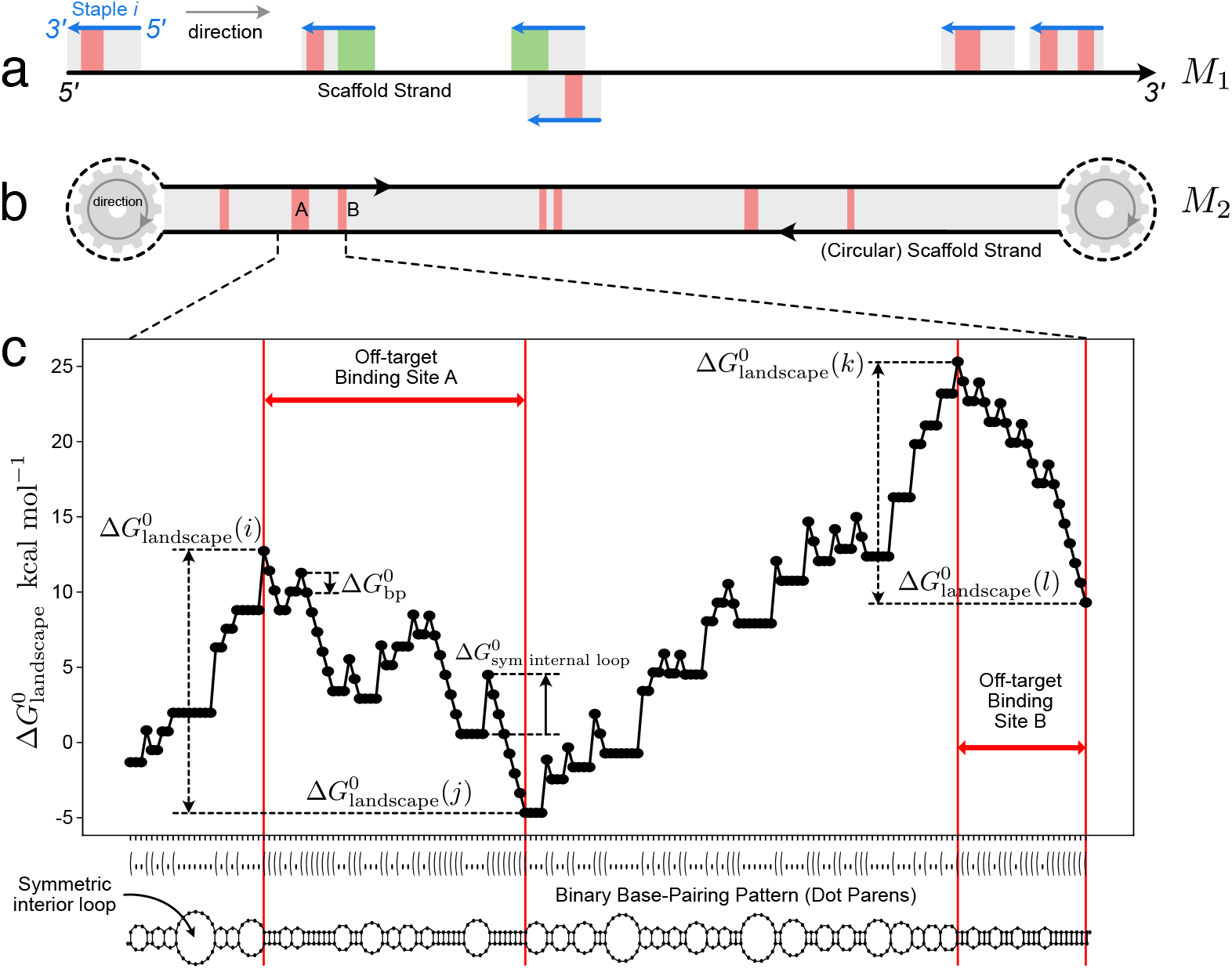
Hybridisation energy model to detect off-target binding sites. Metrics *M*_1_ and *M*_2_ use a fast sequence-averaged energy model of hybridisation to detect staple-scaffold and scaffold-scaffold binding sites respectively. The energy model includes the possibility of symmetric interior loops in binding sites between two nucleic acid strands. **(a)** In Metric *M*_1_, each staple is moved anti-parallel to the scaffold strand in single base increments and the binary base-pairing pattern (‘dot parens’) is calculated at each position from the sequence alignment of the two strands. This binary base-pairing pattern is converted to an approximate hybridisation energy landscape (see panel (c)) which detects both on-target (green) and potential off-target (red) binding sites. **(b)** In Metric *M*_2_, the scaffold strand is aligned against itself to detect scaffold-scaffold binding sites (all of which are off-target). For circular scaffolds, the procedure to find off-target hybridising regions is approximately analogous to stretching the scaffold strand between two rollers; the hybridisation energy landscape of the central ‘helix’ region is calculated for each single base rotation of the rollers. (Note, this does not mean the scaffold strand is being rotated through the origami structure). A section of this energy landscape is shown in panel (c). **(c)** A hybridisation energy landscape is calculated by decrementing the free energy by 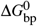 for each base pair encountered and incurring a free energy penalty of 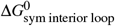 (dependent on loop length) for each symmetric interior loop encountered. A binding site (either on-or off-target) is detected when the difference in free energy between two points on the landscape *i* and *j* is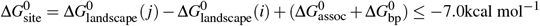.0kcal mol^−1^ at 37°C. The free energy values are parameterised using NUPACK. See Supplementary Note 1 for full details.

### There is plenty of scope for optimising scaffold sequences, especially those arising from biological vectors

To see in which scenarios scaffold sequence selection is most effective at minimising off-target reactions, we performed a large-scale computational investigation that used our method to select scaffold sequences for 14 different 2D and 3D origami designs (Supplementary Notes 4 and 5).

Each origami design had sequence variants (scaffold and complementary staple sequences) selected from three qualitatively different sequence pools; (i) a biological vectors pool containing 5000 scaffold sequences taken from different random contiguous regions of biological vectors pUC19 (2686bp), M13mp18 (7249nt), p7560, p8064 and Lambda DNA (48502bp); (ii) A random pool containing 5000 synthetic random 4-letter scaffold sequences, where each base had an occurrence probability of 0.25; (iii) A de Bruijn pool containing 5000 synthetic de Bruijn sequences (a de Bruijn sequence of order *k* has the special mathematical property that a window of length *k* bases (or larger) will never frame the same sequence fragment twice when it is moved along the sequence base-by-base^22^). We used *k* = 5 for scaffolds 257 to 1024nt; *k* = 6 for scaffolds 1025 to 4096nt, and *k* = 7 for scaffolds 4097nt to 16384nt. Note that the three sequence pools described above were different for each origami. Each sequence pool only provided a very sparse sampling of the combinatorially vast sequence space; however, this sparse sampling was sufficient to create significant variation in the metric *M*_1_ to *M*_4_ scores.

Overall, our computational investigation revealed that the biological vectors pool produced the largest variance in all scoring metrics *M*_1_ to *M*_4_ for each of the 14 test origamis. This is consistent with biological sequences typically having a non-uniform spatial distribution of GC content, which creates higher densities of repeated sub-sequences in specific regions.

Selection of scaffolds for all 14 test origamis from the biological vectors pool above yielded objective spaces where the reverse front and the Pareto front of sequence variants were separated by a large margin. Even for larger 8knt scaffold origamis in the test set, there was still an ≈40% relative reduction in off-target sites from the worst sequence variant on the reverse front (with many off-target reactions) to the best sequence variant on the Pareto front (with fewest off-target reactions). Therefore, we reasoned that scaffold sequences derived from biological vectors were not only the most commonly used to fabricate origamis, but also the most effective target for our multi-objective selection method.

For the synthetic sequence pools, we found that de Bruijn sequences had the best absolute performance across the test set of 14 origamis (Supplementary Note 4). These sequences optimally minimised the total number of off-target reactions between staples and scaffold (but note that *M*_1_ scores did not reduce to zero, since the energy model also acknowledges symmetric mismatches in binding sites that the de Bruijn sequence property does not protect against).

De Bruijn sequences also significantly decreased scaffold-scaffold bindings (*M*_2_). Interestingly, even though the formal de Bruijn sequence property^21^ was not originally developed in relation to nucleic acids – and hence did not consider base-pairing between a de Bruijn sequence and itself – this type of self-interaction also turns out to be quite minimal for de Bruijn sequences.

We found (synthetic) random sequences to minimise scaffold-scaffold interactions equally as well as de Bruijn sequences and to typically perform better than biological sequences (but worse than de Bruijn sequences) at minimising off-target staple-scaffold interactions (*M*_1_). This performance can be attributed to the fact that a non-designed binding region on a random sequence is exponentially less likely to be matched by a complement as it grows in length.

In general, we found little benefit in performing DNA origami scaffold selection from pools of de Bruijn or random sequences because of their generally good baseline performance (but see Supplementary Note 5 for more detail). In particular, de Bruijn sequences *by design*^22^ already incorporated some of the considerations we used in metric *M*_1_, and as mentioned above, also performed well on *M*_2_.

Additionally, we tried selecting scaffold rotations of the published scaffold sequence for those origamis in the test set that had a circular scaffold (where ‘scaffold rotation’ signifies the physical translation of the fixed-sequence scaffold strand around the scaffold routing path in the origami nanostructure by choice of a different set of complementary staples). We observed that rotating the fixed-sequence scaffold strand of a DNA origami through the nanostructure did little to minimise the already present off-target reactions, particularly for larger origamis (Supplementary Note 5). This effect can be anticipated, as rotations of the scaffold only change the staple sequences and different staples inherit similar sequence patterns as the scaffold is rotated. Experimentally, scaffold strand rotations have also been found not to affect assembly yield for large origamis^16,24^.

More generally, measuring the Pearson correlation between all unique pairs of metric scores *M*_1_ to *M*_4_ for all 14 origamis across all sequence pools, we detected no intrinsic linear correlation (Supplementary Note 6) between these metrics. Visual observation of scatter plots also revealed no linear correlations. This signified that the scoring metrics measured independent off-target binding phenomena, as intended.

Finally, the particular MCDM method (SAW, KNEE or TOPSIS) used to select a single origami sequence variant from the Pareto front only made a difference in isolated cases. Largely due to the data normalisation scheme we used (Supplementary Note 3), the MCDM methods approximately agreed about the best candidate sequence variants on the Pareto front and the worst candidate variants on the reverse front in objective space.

### Selection of DNA origami triangle and rectangle variants for *in vitro* assembly

To empirically test if selecting scaffold sequences from regions of biological sequences with relatively many or relatively few off-target reactions had a measurable effect on DNA origami assembly yield, we selected three sequence variants of a 2410nt DNA origami triangle^36^ and three sequence variants of a 2484nt DNA origami rectangle^22^ for *in vitro* assembly. See Figure 3 for details.

**FIG. 3.**
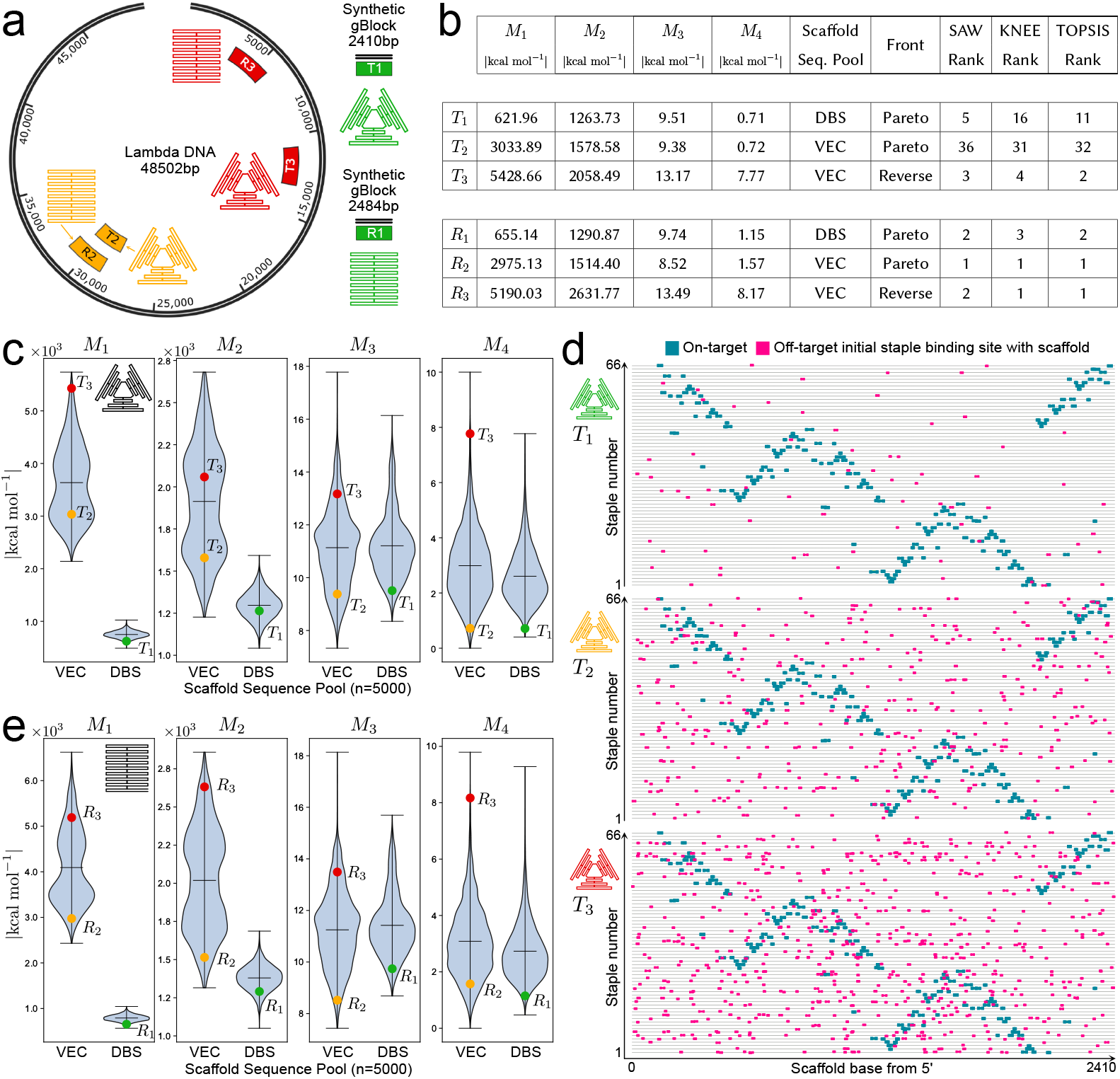
Triangle and rectangle DNA origami sequence variants for *in vitro* assembly. Three variants of a DNA origami triangle *T*_1_, *T*_2_, *T*_3_ (2410nt scaffold) and three variants of a DNA origami rectangle *R*_1_, *R*_2_, *R*_3_ (2484nt scaffold) were selected for *in vitro* assembly. Due to their different scaffold sequences, the origami variants had different extents of off-target side reactions during the self-assembly reaction. **a)** The triangle and rectangle origamis had their scaffold sequences chosen from the worst (*T*_3_, *R*_3_) and best (*T*_2_, *R*_2_) regions of the Lambda DNA sequence, according to scoring metrics *M*_1_ to *M*_4_. Also, each shape had an optimal synthetic de Bruijn scaffold sequence selected for comparison (*T*_1_, *R*_1_). **(b)** Table of metric scores for all origami variants. Lower metric scores indicate fewer off-target binding sites during self-assembly. MCDM rankings were used as a guide to select variants from the Pareto and reverse fronts. **(c)** Full distribution of each scoring metric for the triangle origami, over a pool of 5000 scaffold sequences taken from regions of pUC19, M13mp18, p7560, p8064 and Lambda DNA sequences (VEC) and synthetic de Bruijn sequences (DBS). The location in the distributions of the actual origamis selected for *in vitro* self-assembly is marked. **(d)** Graphical depiction of on- and off-target binding sites existing between staples and scaffold (*M*_1_) for variants of the triangle origami. All variants have the same on-target sites, but *T*_3_ has more off-target staple binding sites than *T*_2_ which in turn has more than *T*_1_. See Supplementary Note 10 for a full analysis of off-target sites. **(e)** Full distribution of each scoring metric for the rectangle origami.

Triangle variants *T*_1_, *T*_2_ and *T*_3_ had a successively increasing number of potential off-target bindings. Variant *T*_1_ acted as a “control”, i.e. best predicted variant, with a synthetic scaffold sequence selected from a pool of 5000 de Bruijn sequences. Variant *T*_2_ was selected from the Pareto front from a pool of 5000 biological vector sequences (derived from contiguous fragments of the pUC19, M13mp18, p7560, p8064 and Lambda DNA sequences), while variant *T*_3_ was selected from the reverse front of the latter pool. Rectangle variants *R*_1_, *R*_2_ and *R*_3_ were obtained similarly.

It should be noted that all sequence variants *T*_1_, *T*_2_, *T*_3_ formed exactly the same triangle design when perfectly selfassembled. Namely, all implemented the same staple pattern, and the same scaffold routing, and all had a staple set that was fully complementary to the scaffold sequence (Supplementary Note 8). Likewise, all sequence variants *R*_1_, *R*_2_, and *R*_3_ implemented the same rectangle design (Supplementary Note 9). The only factor differentiating variants was the number of off-target bindings possible in each case. 2D shapes were chosen for assembly because they deposit flat on mica and are amenable to direct AFM imaging for assessment of assembly quality.

While the 5000 biological vector sequences prepared for each origami came from fragments of five different biological sequences, the scaffold sequences actually selected from the Pareto (*T*_2_,*R*_2_) and reverse fronts (*T*_3_,*R*_3_) all came from the Lambda DNA phage sequence (Figure 3a).

### Synthesis of single-stranded DNA scaffold sequences

Strand-specific T7 exonuclease digestion was employed to synthesise tailor-made single-stranded DNA sequences (ss-DNA) to be used as scaffolds in the folding reactions of all six origami variants (three triangles and three rectangles).

Phosphorothioate protective modification at the 5’ end of the desired ssDNA strand was introduced by polymerase chain reaction (PCR) using a modified forward primer (Supplementary Note 11) as previously reported^37^. In detail, the double-stranded DNAs (dsDNA) of defined sequence and length were amplified from Lambda DNA or gBlocks Gene Fragments templates using a modified forward primer which contains sequential phosphorothioate bonds between the first five nucleotides. The modified phosphate backbone inhibited exonuclease digestion, while the non-required antisense strand was digested by T7 exonuclease. Six different dsDNA sequences were amplified and purified: the correct amplicon sizes were obtained as shown in Supplementary Note 12.

Next, the purified PCR products were selectively digested by T7 exonuclease overnight and purified to remove the polymerase, concentrated and resuspended the ssDNA in water discharging undesired ions. The resulting ssDNA sequences of lengths 2410 nt (triangle scaffolds) and 2484 nt (rectangle scaffolds) showed higher gel mobilities compared to the nondigested dsDNA, as reported in Supplementary Note 12. A faint higher band was also observed in each scaffold variant and was considered as non-digested dsDNA.

Sanger sequencing verified the correct composition of each ssDNA scaffold and underlined no off-target degradation of sense-DNA as previously shown^37^.

### Triangle and rectangle DNA origami variants: folding and pre-screening by gel electrophoresis

The origami self-assembly reactions were run in tris-acetate-EDTA (TAE) buffer containing 12.5 mM of magnesium acetate and with a 10-fold excess of staple strands, a common protocol for 2D origami folding^1,19^.

For each origami variant, 4 different thermal annealing protocols were considered. The mixtures were heated at 95°C for a short period of time and then gradually cooled down to 20°C following a slow (5h 40 min) or a fast (1h 15min) temperature ramp. Isothermal folding protocols were also considered, testing constant annealing temperature (37°C) with or without initial denaturation at 95°C.

To identify potential nanostructure side products (partially assembled and/or mis-folded intermediates) and aggregates, folding solutions incubated at different temperature conditions were first analysed by agarose gel electrophoresis (AGE), a method commonly used to initially assess origami assembly performance. The migration distances, the presence of smearing or multiple bands, and band sharpness were considered indicators of the folding quality, allowing us to select a specific folding condition for subsequent purification and characterisation by atomic force microscopy (AFM). Scaffold, staple strands, and scaffold mixed with non-complementary staples set were considered as negative controls. The formation of DNA origami nanostructures should lead to a mobility shift compared to the scaffold band, which is expected to disappear, while a scaffold mixed with a non-complementary staples set should migrate as the scaffold alone.

The Triangle DNA origami variants *T*_1_, *T*_2_ and *T*_3_ folded in isothermal condition without initial denaturation showed similar band patterns characterised by a main band, a second fainter and diffuse band, and a smearing (Supplementary Figure 26(a),(b) and (c), lane 7). While *T*_1_ and *T*_2_ variants folded in isothermal conditions with initial denaturation were characterised by similar band patterns and migration distance (Supplementary Figure 26(a) and (c), lane 8), the *T*_3_ variant showed aggregates visible in the loading well as a non-migrating band (Supplementary Figure 26(b), lane 8).

Gel electrophoretic analysis of the same variants folded following a fast ramp revealed different band patterns compared to the profiles reported above. In detail, sharp leading bands without intense smearing were visible ( Supplementary Figure 26(a), (b) and (c)), lane 9), while aggregates were still formed in the *T*_3_ variant ( Supplementary Figure 26(b), lane 9). The scaffold mixed with a non-complementary staple set showed the same migration distance as the scaffold alone ( Supplementary Figure 27(b) and (c), lane S).

After 4 days (slow ramp) or 5 days (isothermal folding and fast ramp) at 4°C, the structural stability of the variant assemblies was evaluated. *T*_1_ and *T*_2_ variants showed a similar band intensity ( Supplementary Figure 28(a) and (c)), lane 11) as on day 1 ( Supplementary Figure 26(a), (b) and (c), lane 9), indicating higher stability when compared to the *T*_3_ sample (fast ramp) and other origami samples folded at different temperature conditions.

Based on these results and considering the smearing and fainter bands noticeable in samples folded through a slow temperature ramp ( Supplementary Figure 27(a), (b) and (c), lane 12), we selected the fast ramp for further purification and characterisation by AFM (see below).

As in the triangle DNA origami gel image analysis, rectangle variants *R*_1_, *R*_2_, *R*_3_ folded in isothermal condition without initial denaturation showed a similar band pattern characterised by a main band, a less intense second band and smearing ( Supplementary Figure 29(a),(b) and (c), lane 7). With initial denaturation, *R*_1_ and *R*_2_ variants were characterised by the same band pattern and migration distance with less visible smearing ( Supplementary Figure 29(a),(c), lane 8), while the *R*_3_ variant showed aggregates in the loading well, wide smearing and a main band migrating slightly slower ( Supplementary Figure 29(b), lane 8).

*R*_1_, *R*_2_ and *R*_3_ samples annealed with a fast ramp had a similar electrophoretic behaviour when compared with both isothermal processes, but with a less pronounced smearing and slightly different migration distances ( Supplementary Figure 29(a), (b) and (c), lane 9). The negative controls (scaffold alone and scaffold with non-complementary staple set) showed the same migration distances between them ( Supplementary Figure 30(a),(b) and (c), lane S).

Considering the above results and the fainter bands corresponding to the slower ramp ( Supplementary Figure 30(a), and (c), lane 12), assemblies obtained from the fast ramp were selected, purified, and imaged by AFM (see below). As noted for triangle variants, the rectangle assemblies showed different structural stability when stored at 4°C: after 4/5 days, *R*_1_ and *R*_2_ samples (fast ramp) were more stable ( Supplementary Figure 31(a) and (c), lane 11) compared to the scaffold and samples folded with a slow temperature ramp (S upplementary Figure 31(a), (b) and (c), lane 12), while the *R*_3_ sample (fast ramp) band disappeared ( Supplementary Figure 31(b), lane 11), underlying the assembly instability. The *R*_1_, *R*_2_ and *R*_3_ samples folded in isothermal conditions were relatively stable ( Supplementary Figure 31(a), (b) and (c), lanes 9 and 10).

### Triangle and rectangle DNA origami variants: purification and AFM imaging

Triangle and rectangle DNA assemblies from a fast ramp annealing were purified by centrifugal filtration to remove the low molecular weight excess of staple strands and to concentrate the samples.

The purified reaction mixtures were analysed by AGE and compared with non-purified samples. The centrifugal filters efficiently separated the higher molecular weight DNA assemblies from staple strands. Purified samples (Figures 4 and 5, gel lanes ‘P’) showed the same migration distances as the non-purified samples (Figures 4 and 5, gel lanes ‘A’), suggesting that the purification was damage-free.

**FIG. 4.**
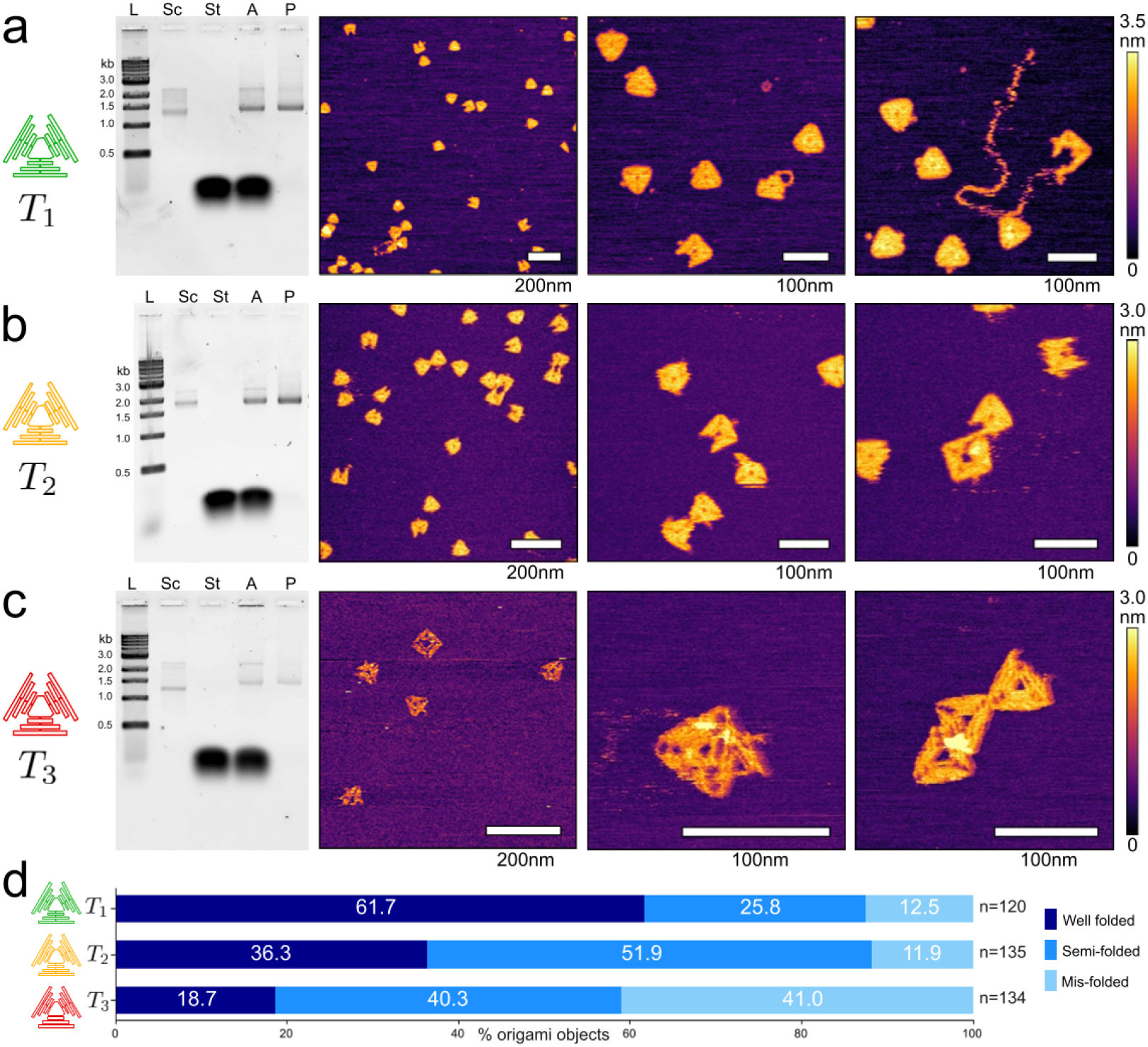
Triangle DNA origami sequence variants: agarose gel and AFM images. **(a)** Triangle variant *T*_1_. **(b)** Triangle variant *T*_2_. **(c)** Triangle variant *T*_3_. In (a), (b) and (c) the first panel shows laser scanned image of SYBR®Gold-stained 1% TBE agarose gel with lanes: L = 1kb Ladder; Sc = scaffold only; St = staples only; A = Origami self-assembly reaction mix (scaffold + staples); P = Purified assembly reaction mix used for AFM imaging. The assembly reaction was performed under a fast temperature ramp of 95°C to 20°C in 1h 15min (−0.1°C per cycle, each cycle 6 seconds). The remaining panels show representative high-resolution AFM images of the purified DNA self-assembly sample (P). **(d)** Manual quantification of origami objects on AFM images into “well-folded”, “semi-folded” and “mis-folded” categories. The percent of origami objects in each category is shown with a mean sample size of 130 objects.

**FIG. 5.**
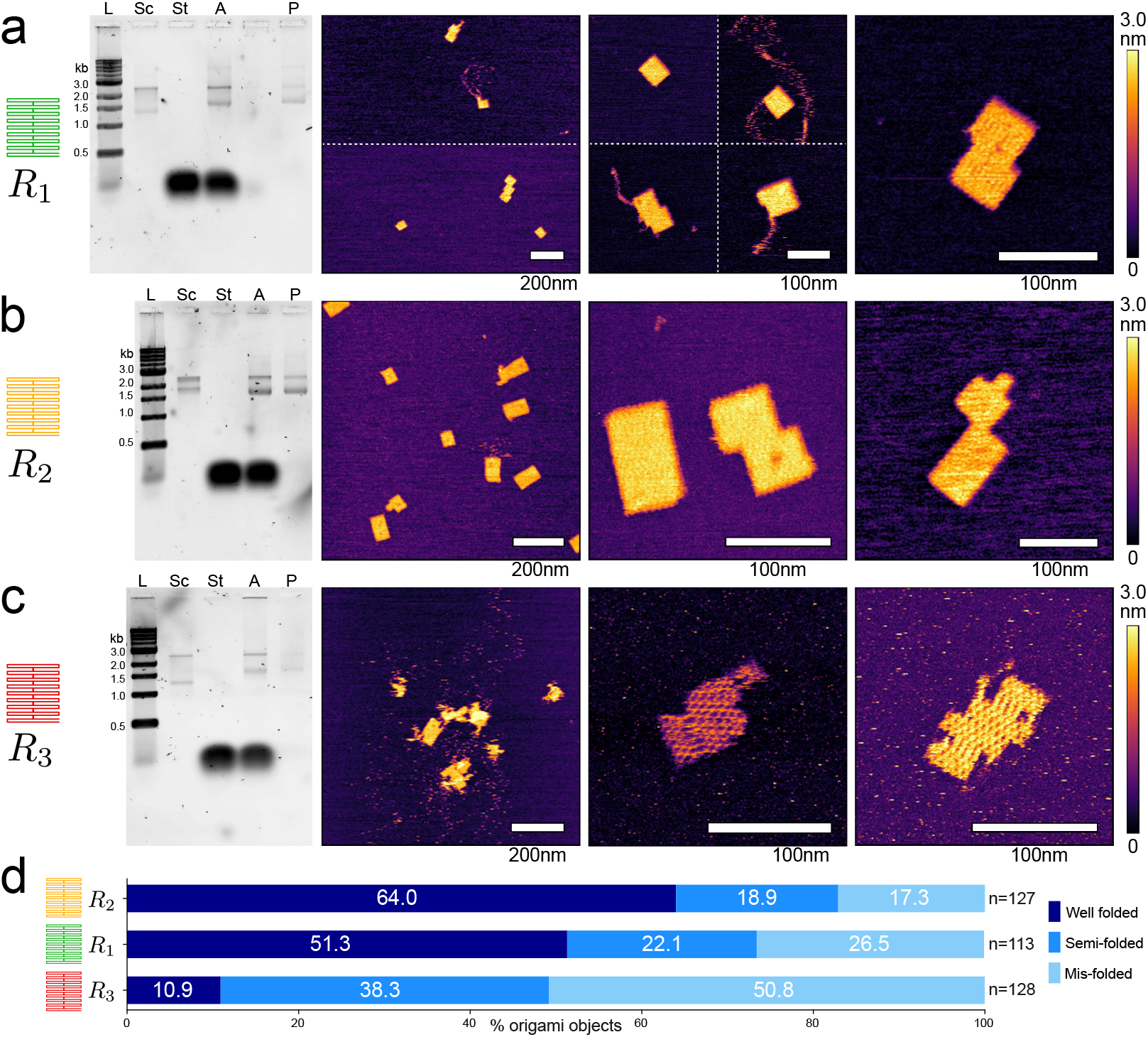
Rectangle DNA origami sequence variants: agarose gel and AFM images. **(a)** Rectangle variant *R*_1_. **(b)** Rectangle variant *R*_2_. **c)** Rectangle variant *R*_3_. In (a), (b) and (c) the first panel shows the laser-scanned image of SYBR®Gold-stained 1% TBE agarose gel with lanes: L = 1kb Ladder; Sc = scaffold only; St = staples only; A = Origami self-assembly reaction mix (scaffold + staples); P = Purified assembly reaction mix used for AFM imaging. The assembly reaction was performed under a fast temperature ramp of 95°C to 20°C in 1h 15min (−0.1°C per cycle, each cycle 6 seconds). The remaining panels show representative high-resolution AFM images of the purified DNA self-assembly sample (P). In (a), dotted lines on AFM images indicate a composite image made of different sample areas to account for low origami adsorption density. **(d)** Manual quantification of origami objects on AFM images into “well-folded”, “semi-folded” and “mis-folded” categories. The percentage of origami objects in each category is shown with a mean sample size of 123 objects.

We classified structures under AFM into three categories: ‘well-folded’ structures resembled the folded shape, ‘semi-folded’ structures resembled the folded shape with small defects, and ‘misfolded’ structures were fragments or aggregates without a specific shape.

The origami triangle AFM images showed that *T*_1_ and *T*_2_ variants (Figure 4(a) and 4(b) respectively) folded with the highest frequency into well-folded triangles (61.7%) and triangles with opened vertices (51.9%), respectively. The *T*_3_ variant (Figure 4c) revealed the lowest percentage of well-folded nanostructures (18.7%) characterised by low stability during the AFM imaging. Semi-folded and misfolded assemblies represented 81.3% of the *T*_3_ origami population. The side average lengths of the *T*_1_ and *T*_2_ variants were compatible with the design, as reported in Supplementary Table 6.

The *R*_1_ and *R*_2_ variants (Figure 5(a) and 5(b) respectively) exhibited higher percentages of well-folded rectangles under AFM (Figure 5d). Conversely, the *R*_3_ variant (Figure 5c) showed a high percentage (about 90%) of semi-folded and misfolded assemblies.

It should be noted that blunt end stacking interactions leading to origami dimers and multimers were observed by AFM imaging on *R*_1_ and *R*_2_ variants (Figure 5(a) and 5(b) respec-tively). In detail, the *R*_1_ variant often formed longer stacked step-like chains compared to the *R*_2_ variant with more frequent dimers. Methods to avoid aggregation based on stacking interactions were not present in the origami designs used. The measured length and width of *R*_1_ and *R*_2_ variants (Supplementary Table 6) were consistent with theoretical values (length and width expected values were 62 nm and 47 nm, respectively).

### Notes on blind experimental protocol and independent replication

The experimental results described above were obtained following a “blind” experimental protocol. Namely, the computational predictions of which scaffold sequence was an rfolder (or not) were obscured from the experimentalists at Newcastle. They later carried out the folding experiments treating all scaffolds in the same manner and analysed their results without an a priori expectation of which scaffold should lead to better AFM results. This blind approach was also replicated independently at the University of Bonn. Bonn’s blind experimental AFM results are in agreement with Newcastle’s for both the triangle and rectangle origami variants (see Supplementary Note 14). Examples of “semi-folded” and “mis-folded” categories were shared by Newcastle to allow a correct quantification of assemblies (see Supplementary Note 16).

### Probing the triangle DNA origami variants: singlemolecule unfolding using optical tweezers

To probe their mechanical response with OT (Supplementary Note 15), triangle origamis were prepared using the fast ramp annealing protocol described above but with two modified staples extending from the 5’ and 3’ ends of the scaffold (Methods). These staples were designed with overhangs that facilitate their ligation to two 2kbp dsDNA “handles”, each modified at the opposite end with either digoxigenin or biotin. The modified dsDNA handles were then attached to two polystyrene beads, coated with anti-digoxigenin and streptavidin, respectively. Each bead was held in a separate optical trap, with one trap fixed in position and the other steerable (Figure 6a). By moving the steerable trap at a constant rate, the force applied to the handles and, consequently, to the ends of the origami structure increased. Initially, only the dsDNA handles were stretched. At higher forces, sufficient to perturb the origami structure, we observed a series of sudden force drops accompanied by increases in extension (Figure 6b, Supplementary Figure 35), likely the result of the release of a single staple or the cooperative release of a number of them, leading eventually to full disassembly of the origami structure. At this point, the ssDNA scaffold was fully stretched, as confirmed by measuring subsequent force-extension curves for those traces that did not break ( ∼ 50%, Supplementary Figure 36a) which are consistent with the size of the scaffold and the elasticity of ssDNA (Supplementary Figure 38). The forces required to unfold the origami structures are generally above 30 pN (Supplementary Figure 36b-c), suggesting that force-induced melting as a result of applying a shearing force between complementary segments (i.e. pulling in the 5’-5’ or 3’-3’ configuration) or stretching one of the complementary segments (i.e. pulling between the 5 and 3’ ends of the same segment) are responsible for the disruption of the corresponding scaffold-staple interactions^38^. However, we observed a small number of events at forces below 20 pN, mainly at the start of the unfolding reaction (Supplementary Figure 36b), indicating that some interactions may involve unzipping events, where the 5’ and 3’ ends of the opposite segments are pulled apart^39^. Comparing the different triangle variants (Figure 6e), reveals a correlation with the construct’s GC contect. *T*_3_ (*GC* = 59%) exhibited the highest unfolding forces, *T*_2_ (*GC* = 44.8%) the lowest, and *T*_1_ (*GC* = 50.4) had intermediate unfolding forces.

**FIG. 6.**
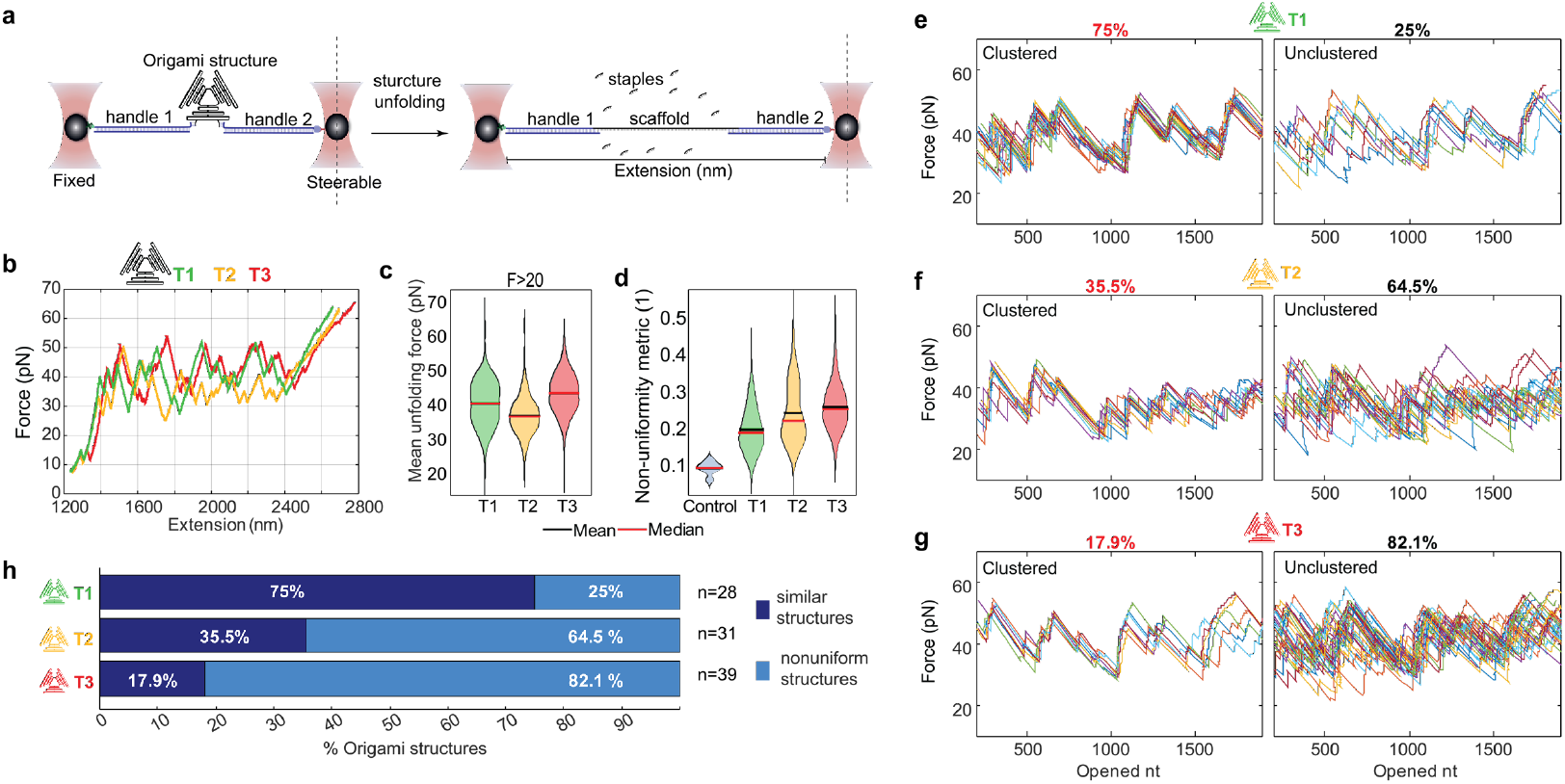
Single-molecule unfolding of DNA origami using optical tweezers. **(a)** Schematic of the optical tweezers setup. The origami structure is anchored at two points to double-stranded DNA handles, which are attached to polystyrene beads held in optical traps. The distance between the traps is gradually increased, unfolding the origami structure, resulting in a single-stranded DNA (ssDNA) scaffold with staples irreversibly lost to the solution. **(b)** Representative unfolding force curves are shown for each triangle variant, with different colours representing each, plotted as a function of the extension. **(c)** Violin plot of unfolding forces above 20 pN, derived from all unfolding curves for each variant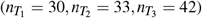. p-values obtained from Students T-tests are shown in **Supplementary Table 8. (d)** Violin plot of the non-uniformity metric, comparing a DNA unzipping control group with the triangle variants, derived from all the unfolding curves that exceed 50% of the scaffold length.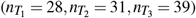. p-values obtained from Students T-tests are shown in **Supplementary Table 9. (e)**,**(f)**,**(g)** Force vs. opened nucleotides (nt) curves for clustered structures (left) and unclustered structures (right) for each T variant: (f) *T*_1_, (g) *T*_2_, and (h) *T*_3_. (h) Bar plot showing clustering percentages for each group. The threshold for the clusters displayed in this figure is set to 0.14.

Notably, there is a significant degree of non-uniformity in the force-extension curves, as evidenced by comparing with a set of curves obtained from unzipping a long DNA hairpin (Supplementary Figure 35). A higher disorder for the origami is expected, since in contrast to the vectorial DNA unzipping where there is only a single unfolding pathway possible, multiple pathways likely exist for a DNA origami (Supplementary Figure 40). However, structural variations resulting from the possibility of off-target binding reactions are likely to contribute to the disorder too, so we expect curves for *T*_1_ and *T*_2_ to be more uniform than those for *T*_3_. To assess the structural nonuniformity of a given origami variant, we defined a nonuniformity metric (NUM) that quantifies the similarity between all possible pairs of traces by calculating the cumulative difference in the position-dependent force (see Methods). As a control, we used DNA unzipping data, which exhibits high uniformity across experiments. NUM values for all origami variants were higher than those of the control, with *T*_1_ showing the lowest mean NUM, followed by *T*_2_, and then *T*_3_ (Figure 6d). Using the NUM values, and a threshold, we then clustered similar traces (see Methods). Remarkably, this analysis revealed that while ∼ 75% of *T*_1_ variants could be clustered into a group of similar traces, and ∼ 36% of the *T*_2_ variants, only ∼ 18% of the traces for *T*_3_ could be clustered (Figure 6e–h), and this trend was insensitive to the choice of threshold (Supplementary Figure 37d). These results indicate that *T*_1_ structures are assembled with higher uniformity than the other variants, followed by *T*_2_, and then *T*_3_, consistent with the AFM data.

## DISCUSSION

The AGE and AFM results we obtained from sequence variants of the triangle and rectangle DNA origami, and the optical tweezers (OT) results for the triangle ones, confirm our computational predictions of good/mid/bad folding scaffold sequences, and broadly support our main hypothesis that increased off-target binding possibilities have a negative effect on origami assembly yield.

In the extreme cases, both the *T*_3_ and *R*_3_ variants (with many potential off-target reactions of all four types) nearly completely failed to fold into the intended triangle and rectangle shapes (only 10%-20% well-folded AFM yield), despite the fact that the staple strands were perfect Watson-Crick complements to the scaffold in both cases. Moreover, the REVNANO tool^40^, which relates to metric M1, had lower performance while attempting to reverse engineering T3 and R3 designs when compared with the others.

*T*_3_ variants also showed a high degree of non-uniformity in OT experiments, and almost completely failed to cluster into a single uniform group.

As mentioned, the *T*_3_ and *R*_3_ variants also displayed well aggregates in AGE when only scaffold strands were present (Supplementary Note 13) and also for the full assembly reactions (Figure 4c and Figure 5c respectively, lane ‘A’), suggesting that for these origamis, scaffold strands were hybridised together either directly and/or indirectly during assembly.

Therefore, while origami self-assembly undoubtedly has a number of self-correcting error mechanisms to recover from moderate off-target binding ‘noise’ (like strand displacement of off-target staple-scaffold and scaffold-scaffold bindings by invading on-target staple strands), our results support the view that it is still possible to kinetically trap origami folding if off-target reactions are too prevalent. This view challenges received wisdom in the DNA nanotechnology community that, an origami will always self-select the minimum free energy structure (with all staples bound in their designed complementary positions) from any initial scaffold/staple strand cocktail, for reasonably heterogeneous sequences. Our results support the view that scaffold sequences (and their complementary staples) that fail to fold into the target origami structure can be found as unlucky regions of commonly used sequences like that of the Lambda DNA phage.

Independent replication of folding reactions and AFM experiments at the University of Bonn, Germany (Supplementary Note 14) generally supported the results presented here. Triangle *T*_1_ (de Bruijn sequence) folded with the highest AFM yield, followed by *T*_2_ and then *T*_3_. The same anomaly was also observed for the rectangle origami: *R*_2_ folded with the highest AFM yield, followed by *R*_1_ and then *R*_3_. This anomaly, where *R*_2_ with more off-target reactions assembles at a higher yield than the de Bruijn sequence control *R*_1_ may point to the fact that there may exist other, hidden sequence design principles that can increase assembly yield in the case of specific origami shapes.

At this stage, it is instructive to reflect on some aspects of our computational method to select origami scaffold sequences, and then proceed to discuss some extensions and limitations of our approach.

A first reflection is that our method is a heuristic, pragmatic approach to increase folding yield for a pre-specified origami design. It indicates which sequences should probabilistically fold better, relative to others, rather than directly predicting yield percentage, or directly specifying where fold defects will be located. Our approach works with static sequences, not simulation trajectories of the origami folding pathway, and as such cannot predict classic assembly features such as structure melting temperature, speed of folding, or potential cooling/re-heating hysteresis. However, we note that because our approach rewards sequences which essentially obey domain-level behaviour (i.e. where the only binding events are the intended on-target bindings between staples and scaffold), then our method could be used to select scaffold/staple sequences which are better described by current origami folding models^6,7,30^.

A second important note is that, in the absence of better information, we assume that the four scoring metrics used in Figure 1 have equal weightings in the final MCDM score for a scaffold sequence. That is, we assume that each different type of off-target binding (staple-scaffold, intra-scaffold, inter-staple, and intra-staple) can equally disrupt the origami folding pathway. Equal weights were sufficient to allow us to predict ‘good’ and ‘bad’ scaffold sequence regions from the lambda DNA sequence for the triangle and rectangle DNA origamis in this study. However, we expect this weighting could be refined in future if large data sets became available relating sequences and buffer conditions to a uniform measure of origami folding yield. In particular, it seems *M*_2_ would be a suitable candidate to weight more heavily, given that scaffold-scaffold bindings are an intra-molecular reaction that occurs at high effective concentration (j-factor), and also because our results evidence that scaffolds can bind inter-molecularly and aggregate.

A third point is the observation that more simple metrics like scaffold GC content and *k* − *mer* subsequence repeats are indeed correlated with scoring metrics *M*_1_ to *M*_4_, but not tightly enough to substitute any of them (Supplementary Note 7). Unlike the latter metrics, which are just based on scaffold sequence, scoring metrics *M*_1_,*M*_3_ and *M*_4_ are specific to a certain origami design and all four metrics precisely enumerate potential off-target binding sites. However, GC content could still be leveraged to pre-screen scaffold sequences: those sequences outside a 45-55% GC content range could be omitted from the scaffold sequence pool as they are likely to lead to high numbers of sub-sequence repeats and thus higher *M*_1_ and *M*_2_ scores.

Our approach has considerable scope for novel extension and refinement. Most obviously, the number and type of scoring metrics could be further investigated.

For calculating off-target binding sites with greater accuracy, metrics *M*_1_ and *M*_2_ could use finer-grain averaged energies of base-pairing. Instead of a single per-base pair average binding energy Δ*G*_bp_, two different average free energies Δ*G*_bp_(*GC*) *<* Δ*G*_bp_(*AT*) could be pre-calculated and employed. This modification could be particularly useful for RNA origami (see below).

Metric *M*_3_ could be modified such that essential staples (e.g. those performing functions like dangling end docking points on origami memory tiles^41^) provide a higher contribution to the *M*_3_ score, thus ensuring that essential staples have a low probability of being co-folded. Additionally, the effect of making *M*_3_ a sum rather than a maximum could be rigorously explored.

To reduce the dimensionality of the objective space and to narrow the Pareto front of solutions, Metric *M*_4_ could conceivably be omitted from the selection step and instead applied post-selection as a filter. This is because strong intra-staple hairpins are actually relatively rare due to (i) the relatively small number of individual staples per structure (with respect to, for example, the number of possible staple-staple co-folds) and (ii) the typically short size of staples. If candidates on the three-dimensional {*M*_1_, *M*_2_, *M*_3_} Pareto front had a high *M*_4_ filter score, they would simply be discarded.

A new benefit metric which rewards stronger on-target staple bindings could potentially be added; then sequence selection would boost favourable interactions (positive design) as well as reduce unfavourable off-target interactions (negative design). Alternatively, a metric to assist to folding of dense 3D structures could be specified, whereby on-target binding sites in the interior of the object could be rewarded if they were rich-GC, as to increase the probability that they hybridise first before hybridisation is blocked by steric hindrance from folded outer parts.

As each sequence pool considered by our method only provides a sparse sampling of the combinatorially vast sequence space, and considering that even under this sparse sampling significant variation in the metric scores was observed, a new sequence sampling technique that couples e.g., Design of Experiments or generative AI sampling methods with our various metrics might lead to better reliable folders.

Finally, it is useful to consider the limitations of our method. The first main limitation is computation time. Even though the method works with static sequences, the time to compute the objective space for large origamis (e.g. 7knt scaffold) can be significant on a single CPU (on the order of days) because of the exact enumeration of all off-target binding sites. However, the codebase does allow use of parallel computing in a cluster environment to speed up calculations. In future, smart caching of energy landscapes and/or optimisation of metric *M*_1_ and *M*_2_ scoring code could accelerate performance.

A second limitation is applying our method to larger origamis where the scaffold sequence is fixed and only the scaffold strand can be rotated (a common scenario). As mentioned in the results, strand rotation for larger origamis (thousands of scaffold bases) typically causes negligible variance in the metric scores. As stated, our method is best applied to select origami scaffold sequences from regions of biological sequences which are longer than the origami scaffold.

The final challenge is application to RNA origami. An RNA energy model was also implemented and it used a higher average energy per base pair than for DNA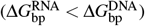, in addition to considering GU pairs as standard base pairs. However, these two factors lead the RNA energy model to identify significantly more off-target sites (in all four categories) for the RNA sequence equivalent version of a DNA origami. A future RNA energy model would benefit from (i) employing not one but three base-pair binding energies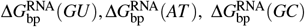, each averaged over all possible nearest neighbour combinations, to make off-target site identification more accurate and (ii) experimental testing of RNA origami self-assembly, as done for DNA origami in this study. A hybrid DNA/RNA energy model suitable for origami assembling *in vivo* could also be parameterised in the future^42^.

## CONCLUSION

Computational efforts to improve DNA origami nanotechnology have traditionally focussed on optimising origami scaffold and staple strand routings in the origami design, rather than on primary base sequence optimisation. Nevertheless, base sequences are able to cause long-range off-target interactions between multiple components of the system which complicate and potentially trap the self-assembly process.

In this study, we developed a multi-objective computational method to select scaffold sequences which minimise four different types of off-target reactions. We demonstrated that off-target side reactions during self-assembly can significantly affect the final AFM yield of two different target origami shapes. We also showed that scaffold sequences for the same designed shape, which are optimised to different degrees by our algorithms, have, under single-molecule optical tweezer experiments, distinct degrees of uniformity in their force profiles, which are linked to our computational predictions.

Overall, this work proposes a change in perspective about DNA origami base sequence. Rather than a top-down engineering view, we advance a more bottom-up systems view of the origami self-assembly process. Instead of the base sequence being seen as a benign low-level implementation of the desired staple-scaffold domain-level bindings, this study suggests that potential off-target interactions inherent in the particular base sequences employed should be taken more seriously, particularly when biological scaffold sequences are used.

## METHODS

### Blinded experiments

DNA origami folding reactions were carried out with blinded samples to mitigate experimental bias. In the methods below, origami variants are referred to by the following names: *T*_1_=DEER, *T*_2_=LION, *T*_3_=BEAR, *R*_1_=GOAT, *R*_2_=LAMB, *R*_3_=MOLE.

### Materials and reagents

All DNA oligonucleotides and gBlocks Gene Fragments were purchased from IDT. The DNA oligonucleotides were resuspended in UltraPure DNase/RNase-free distilled water (Thermo Fisher Scientific) to give stock solutions of 100*µ*M and stored at −20°C. The gBlocks Gene Fragments were resuspended in Tris-EDTA (TE) buffer at a final concentration of 10 ng/*µ*L and stored at −20°C.

EDTA 0.5M pH 8.0, DNase/RNase-free Tris 1M pH 8.0, SYBR^TM^ Gold nucleic acid gel stain, UltraPure 10X TBE buffer, 10X TAE buffer molecular biology grade (400 mM Tris and 0.01 M EDTA), Ambion®10X gel loading solution were purchased from Thermo Fisher Scientific.

Q5®High-Fidelity 2X Master Mix, Monarch®PCR & DNA Clean-up kit, 1 kb and 1 kb Plus DNA ladders, Gel loading dye purple (6X), T7 exonuclease and Lambda DNA (500*µ*g/mL) were purchased from NEB.

Agarose, Nancy-520, magnesium acetate BioUltra 1M in water, Amicon®Ultra 100 kDa centrifugal filters were purchased from Sigma-Aldrich.

### Double-stranded DNA amplification, purification, and agarose gel electrophoresis

Double-stranded gBlocks Gene Fragments and Lambda DNA were amplified using Q5®High-Fidelity DNA polymerase 2X master mix, 10*µ*M of 5’-phosphorothioate modified forward primer and 10*µ*M of reverse primer (Supplementary Table 4): the final concentration of each primer was 0.5*µ*M in 50*µ*L of final reaction volume. When the origami centrifugal purification and imaging were considered, a total of 20 PCR tubes were prepared for each DNA origami variant. The cycling conditions and the template quantity (ng in 50*µ*L) for each variant are reported in Supplementary Table 5. The PCR product was purified using Monarch®PCR & DNA Cleanup kit: PCR solutions diluted with DNA cleanup binding buffer were loaded onto 4 distinct columns, washed, and eluted in 10*µ*L of UltraPure DNase/RNase free distilled water.

The DNA concentration of the total eluted volume (≈32*µ*L) was measured using a NanoDrop One/OneC spec-trophotometer. After the addition of the purple gel loading dye (2*µ*L) to 10*µ*L of TE containing 100ng of purified dsDNA, the size of purified amplicons were evaluated on 1% agarose gel in TBE for 1h at 110V: the gels were pre-stained with Nancy-520, visualized using Typhoon laser scanner (*λ*_exc_ 488nm, *λ*_em_ 532nm, PMT voltage 380, normal sensitivity, GE Healthcare Life Sciences) and ImageQuant TL software (GE Healthcare Life Sciences). The 1 kb DNA ladder (NEB) was used as molecular weight marker.

### Purified double-stranded DNA digestion using T7 exonuclease, purification, and agarose gel electrophoresis

Amplified and purified double-stranded DNA (dsDNA) template was used for single-stranded DNA (ssDNA) preparation, as previously described^37^ with some modifications. Briefly, 6 digestion mixes were prepared for each variant: one reaction mix contained 20 units (Deer, Lion), 25 units (Bear, Lamb) or 30 units (Mole, Goat) of T7 exonucleases, 5*µ*L NEB Buffer 4, 500ng of purified dsDNA template and UltraPure DNase/RNase free distilled water (up to 50*µ*L). After overnight incubation at 25°C, the ssDNA product was purified using Monarch®PCR & DNA Clean-up kit: digested solutions diluted with DNA cleanup binding buffer were loaded onto 3 distinct columns, washed, and eluted in 12*µ*L of UltraPure DNase/RNase free distilled water. The ssDNA concentration of the total eluted volume (≈ 30*µ*L) was measured using a NanoDrop One/OneC spectrophotometer. After the addition of the purple gel loading dye, the purified ssDNA was run on a 1% agarose gel in TBE and imaged as described above. The ssDNA solution was kept at −20°C until the origami folding reaction (the day after). Each purified ssDNA scaffold was sent for Sanger sequencing (Eurofins Genomics) and aligned using the SnapGene®software.

### Triangle and rectangle DNA origami folding reactions and pre-screening by agarose gel electrophoresis

For each DNA origami variant folding (3 triangle DNA origami variants and 3 rectangle DNA origami variants), each ssDNA staple strands set (final concentration 16nM each) was mixed in a 10-fold excess with complementary ssDNA scaffold (1.6nM) in 50*µ*L of folding buffer (1x TAE molecular biology grade buffer, 40mM Tris Acetate, pH 8.3, 1mM EDTA, supplemented with 12.5mM BioUltra magnesium acetate) as previously reported^19^. Folding solutions and negative controls (only scaffold and only staple strand samples) for each variant were incubated at 4 different temperature conditions and then held at constant temperature (4°C) to stop the reaction. The tested temperature conditions were: i) isothermal folding at 37°C for 1 h 15 min without initial denaturation; ii) isothermal folding at 37°C for 1 h 15 min with an initial denaturation at 95°C for 1 min; iii) fast temperature ramp from 95°C to 20°C in 1 h 15 min (−0.1°C/cycle, each cycle 6 sec); iv) slow temperature ramp from 95°C to 20°C in 5 h 40 min (−0.1°C/cycle, each cycle 27 sec).

Scaffold, staple strands, and DNA origami samples (10*µ*L) were run on a 1% agarose gel in 1x TBE buffer at 90 V for 1 h 20 min at low temperature (below 10°C). Samples annealed considering the first 3 temperature ramps (isothermal folding with or without initial denaturation and fast ramp) were kept at 4°C and run again with samples annealed with a slower ramp the day after. The gels were stained in 1x TBE containing SYBR®Gold for 15 min under gentle agitation at 15 rpm (Gyro-rocker SSL3, Stuart) and visualized using a Typhoon laser scanner. The 1 kb and 1 kb Plus DNA ladders were used as molecular weight markers. To monitor the structural stability of the assemblies over the time, the gel electrophoretic analysis was repeated after 4/5 days on all samples stored at 4°C.

### DNA origami purification and atomic force microscope (AFM) imaging

The folded constructs at a specific selected temperature condition were purified from staple strands excess using the Amicon®Ultra 0.5 mL 100 kDa centrifugal filters. Samples were added to the filter device and centrifuged at 14000g for 1 min at 11°C. The flowthrough was removed and the filter was washed 4 times (14000g for 1 min at 11°C) with 450*µ*L of 1x folding buffer. To recover the purified and concentrated sample, the filter was turned upside down and centrifuged at 1000g for 5 min at 11°C. Scaffold, staple strands, non-purified and purified DNA origamis were run on 1% agarose gel in 1x TBE buffer, stained and visualized using a Typhoon laser scanner as previously reported.

Purified DNA origamis samples were stored at 4°C and imaged by AFM as follows. Freshly cleaved mica (Mica Grade V-4 12 mm Discs x 0.15 mm, Azpack Ltd., Loughborough, UK) was passivated using 1x folding buffer. After three washing steps with folding buffer (60*µ*L each), the purified construct solution was added to the passivated mica surface, allowed to absorb for 1-2 min and imaged immediately (≈ 100*µ*L of the folding buffer were added after the origami absorption). The purified assemblies were characterised in tapping mode in liquid using VRS Cypher ES AFM (Asylum Research, Oxford Instruments, Santa Barbara, CA, USA). The vertical oscillation of the BioLever Mini probe (spring k of 0.09 N/m, Asylum Research, Oxford Instruments, Santa Barbara, CA) was controlled by photothermal excitation (Blue Drive) and the imaging was conducted at 25.0 *±* 0.1°C.

Large scan size (about 1*µ*m x 1*µ*m) and zoomed scans were considered to count a total of about 100 nano-objects for each variant, considering well folded, folded with defects and unfolded assemblies. All the images were corrected for tilt (line or plane flattening) and lightly low pass filtered to remove high-frequency noise, using the Asylum Research analysis package for Igor Pro(R) (Wavemetrics, OR, USA)^43^.

### Triangle and rectangle DNA origami: folding reactions, purifications and AFM imaging conducted at University of Bonn, Germany

To validate our approach, data from another nano biotech laboratory was considered. The blind experiments protocol was uploaded in protocols.io^44^ to share an up-to-date version: few changes have been considered and tracked. In detail, the amplified and purified dsDNA templates were digested for 4 hours instead of overnight, and the AFM imaging was modified as follows. Freshly cleaved mica was passivated using poly-L-ornithine instead of 1X folding buffer. After samples deposition, high-resolution AFM images were obtained using a JPK NanoWizard®3 Ultra AFM (JPK Instruments, Berlin). The USC-F0.3-k0.3-10 ultra-short AFM tips (NanoWorld Innovative Technologies, Switzerland) with a nominal force constant of 0.3 N/m were used.

### Optical-tweezers experiments

#### Molecular constructs

Experiments were conducted using a custom-built double-trap optical tweezers setup, as previously described^45^. Constructs to probe the force response of the origamis were based on previous designs aimed at probing protein-DNA interactions^46,47^, with modifications. We generated two 2000 bp DNA handles, each uniquely tagged with double digoxigenin or biotin, by performing PCR on bacteriophage lambda DNA. The biotin handle was created using a biotin-tagged forward primer, while the digoxigenin handle used a forward primer containing a BglI restriction site to create an overhang for ligation with two annealed, commercially purchased oligos featuring 3’ and 5’ digoxigenin terminal modifications (IDT, Supplementary Table 11). The reverse primers were designed with restriction sites for NcoI and BglI for the biotin- and digoxigenin-tagged handles, respectively. Digestion with NcoI-HF and BglI (New England Biolabs) was done following the manufacturer’s recommended protocol, resulting in distinct overhangs for each handle. For the optical tweezers experiment, origami structures were designed with complementary overhangs at both scaffold ends, matching the overhangs on the two DNA handles. The origami structures were first assembled as described above. Subsequently, the handles were ligated to the origami structures at an equimolar ratio using a rapid ligase system (Promega), with incubation at 16°C for 30 minutes. Force-extension curves acquisition. The full construct (handles + origami structure) was incubated for 15 minutes on ice with 0.8 *µ*m polystyrene beads (Spherotech) coated with anti-digoxigenin antibodies. The reaction was then diluted 1000-fold in a working buffer (10 mM Tris-Cl pH 7.4, 150 mM NaCl, 1.5 mM MgCl_2_, 3% glycerol, and 0.01% bovine serum albumin). Tether formation was conducted in situ within the experimental chamber at room temperature by capturing an anti-digoxigenin bead bound to the DNA construct in one optical trap and a 0.9 *µ*m streptavidin-coated polystyrene bead in the other trap. The two beads were brought into close proximity to allow binding between the biotin tag on the DNA and the streptavidin on the bead (Figure 6a). Next, the distance between the two beads was gradually increased at a constant rate until reaching a maximal force of 65 pN to fully unfold the origami structure. If the tether remained intact, the distance between the beads decreased and then increased again, to reveal the stretching of the scaffold and the existence of any possible residual structure (Supplementary Figure 38). Data analysis. Data were digitized at a 2500 Hz sampling rate and saved to disk, with all subsequent data processing performed in MATLAB (MathWorks). From the measured tether extension and force, the extension due to dsDNA handles was calculated and subtracted at each time point (Supplementary Figure 39b). The resulting extension was then divided by the extension of one ssDNA base (calculated from the measured force using the worm-like-chain model) to yield the number of opened nucleotides (Supplementary Figure 39b). Data collection statistics are summarized in Table 1. The nonuniformity metric (NUM) was calculated for each pair of traces obtained within the same variant type. For each pair, we low-pass filtered the data to 10 Hz and focused on the overlapping region of the force versus num-ber of opened nucleotides (nt) curves. The NUM (Eq. 1) was defined as the root of the mean squared force difference, normalized by the mean peak-to-peak force difference for each variant (Supplementary Table 10):

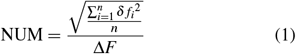

To account for instrumental variability between different unfolding curves, we scanned the NUM for each pair by shifting the two traces along the open nt axis (±150 nt in 5-nt increments) and along the unfolding force axis (±3 pN in 0.5-pN increments). The lowest NUM value was assigned to each pair of curves. Clustering of similar traces is achieved by first connecting traces with a NUM below a specified threshold (0.14 in Figure 6e-h), and then grouping traces that share at least one overlapping connection. To qualify as a cluster, a group must consist of at least three traces; groups with fewer traces are not considered clusters (Figure 6e-g). If more than one cluster existed, the largest resulting cluster was chosen and traces that belonged to other groups were classified as unclustered. The unfolding curves shown in Figure 6(e)-(g) were aligned to a randomly chosen reference curve from the similar trace cluster, using the shift along the open nt and unfolding force axes that yielded the lowest NUM value.

## ACKNOWLEDGEMENTS

This work was supported by the Department for Science, Innovation and Technology (DSIT) and the Royal Academy of Engineering Chair in Emerging Technologies award to N.K. It was also partly supported by a Horizon 2020 project “AI-enabled RNA nanotechnology DElivery SysTem for INfor-mATION transfer into cells.” under grant agreement 899833 to N.K and E.T, and a Royal Society International Exchanges grant IES/R1/180080 to B.S-E and J.E. A.K. acknowledges the Israel Science Foundation for grant 937/20. We are very grateful to Prof. J. Bacardit for facilitating access to Newcastle University ROCKET cluster for high-performance simulations and we thank Prof. K. Voitchovsky for discussions on experimental aspects of this work.

## CODE AVAILABILITY

Python source code (Academic Noncommercial License) and documentation is located at https://scaffoldselector.readthedocs.io. A permanent snapshot of the source code, and scadnano designs of the origami variants used is located at DOI 110.5281/zenodo.14749729.

## SUPPLEMENTARY INFORMATION

Supplementary information associated with this article can be found online.

