## Supplementary material for "Optimizing DNA origami assembly through selection of scaffold sequences that minimise off-target interactions": Suplementary Information

#### Contents

**I Formulae and Definitions** **3**

**Supplementary Note 1**
**DNA Approximate Energy Model for Metrics 1 and 2** **3**

**Supplementary Note 2**
**Definitions of Metrics 1, 2, 3, 4** **9**

†These authors have contributed equally to this work.

Supplementary Note 3
Pareto Front Multi-Criteria Decision Making 16

### II Computational Results 21

Supplementary Note 4
Test Set of 14 DNA Origamis 21

Supplementary Note 5
Scaffold Sequence Selection from Biological Vectors, de Bruijn and Random Sequences 23

Supplementary Note 6
Metric Pair Correlations 29

Supplementary Note 7
Metric Score Correlation with Repeats and GC Content 30

### III DNA Origamis Selected for *In Vitro* Assembly 31

Supplementary Note 8
Triangle Origami Variants 31

Supplementary Note 9
Rectangle Origami Variants 34

Supplementary Note 10
Off-target Binding Site Energy Distributions 37

### IV Additional Experimental Results 38

Supplementary Note 11
PCR Details 39

Supplementary Note 12
Single Stranded DNA Scaffold Purification 41

Supplementary Note 13
Gel Electrophoresis Pre-Screening 44

Supplementary Note 14
Independent Replication of Results at University of Bonn 50

Supplementary Note 15
Optical Tweezers 54

Supplementary Note 16
AFM Examples of "Semi-Folded" and "Mis-Folded" Origamis 59

#### Part I

### Formulae and Definitions

In this part, we detail the main technical aspects of scaffold sequence selection: identification of on-target and off-target binding sites with a sequence-averaged energy model, definitions of scoring metrics 1-4 and MCDM methods used to choose an 'optimal' candidate from the Pareto front in objective space.

#### Supplementary Note 1 DNA Approximate Energy Model for Metrics 1 and 2

A fast sequence-averaged energy model is used to identify and calculate the (approximate) free energy of on-target and off-target binding sites in metrics $M_1$ and $M_2$.

Within a sliding window region, the energy model treats the two aligned strands as a duplex with permitted symmetric interior loops and calculates an energy landscape based only the pattern of base matches and mismatches between the two strands (Supplementary Figure 1(a)). An average base-pairing energy is used instead of a sequence-dependent nearest-neighbours approach. Distinct binding sites are then identified along the energy landscape as explained below.

##### Energy Landscape Construction

An approximate energy landscape is constructed from only the dot parens string of the sliding window duplex by using sequence-averaged base pairing parameters and sequence-averaged interior loop parameters (Supplementary Table 1). Specifically, the landscape free energy at base $i$ is calculated as:

$$\Delta G^0_{\text{landscape}}(i) = b_i \Delta G^0_{\text{bp}} + \sum_{l \epsilon L(i)} \Delta G^0_{\text{sym internal loop}}(l) \tag{1}$$

The first RHS term is the sum of the base pairing enthalpies up to point $i$, where $b_i$ is the number of basepairs up to and including base $i$ of the duplex and $\Delta G^0_{\text{bp}}$ is the average per-basepair hybridisation energy for DNA or RNA. G-U pairs are counted as basepairs in RNA.

The second RHS term is the sum of entropic penalties of all closed symmetric interior loops up to point $i$. Set $L(i)$ contains the length of all symmetric interior loops up to point $i$ and $\Delta G^0_{\text{sym internal loop}}(l)$ is the free energy penalty of an interior loop of size $l$. To minimise calculations, $\Delta G^0_{\text{sym internal loop}}(l)$ is read from a lookup table for loop sizes $1 \leq l \leq 30$, and different lookup tables are used for DNA and RNA. In the $l > 30$ case, for both DNA and RNA, the Jacobson-Stockmayer loop entropy approximation [? ] is used instead:

$$\Delta G^0_{\text{sym internal loop}}(l) = -T \Delta S^0_{\text{loop}} + \gamma RT \ln\left(\frac{l}{30}\right) \tag{2}$$

where gas constant $R = 1.9872036 \times 10^{-3}$ kcal K$^{-1}$ mol$^{-1}$ and $T = 310.15$ K. See Supplementary Table 1 for parameters.

Note that:

- Absolute energies along the energy landscape are not meaningful: only energy differences between two points on the energy landscape are meaningful.

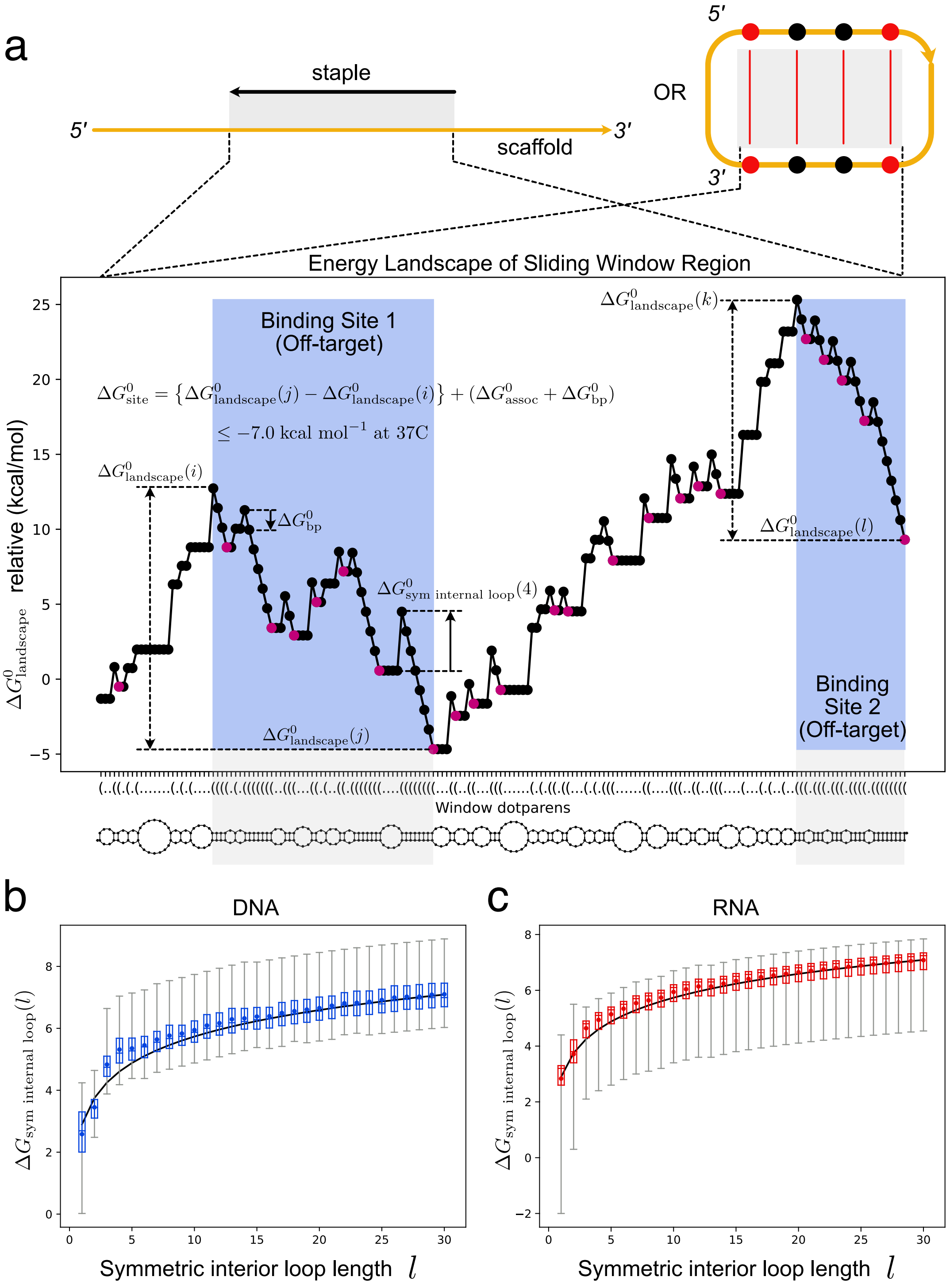


Supplementary Figure 1: Energy Model for metrics $M_1$ and $M_2$. **(a)** A sliding window region (staple-scaffold, or scaffold-scaffold) is treated as a helix duplex with symmetric interior loops. An energy landscape is constructed from the dot parens binding pattern of the sliding window region. Two distinct binding sites are identified in this example. **(b)** and **(c)**: DNA and RNA free energy penalties for different interior loop sizes at 37C, respectively. Free energy penalty distribution at each loop size computed from $n = 500$ samples of NUPACK `structure_energy()`, see text. Dots show mean free energy penalty values used as the lookup value for each loop size. Boxes show median and interquartile free energy range. Grey lines show minimum and maximum free energy values. Black lines show the Jacobson-Stockmayer approximation of equation (2).

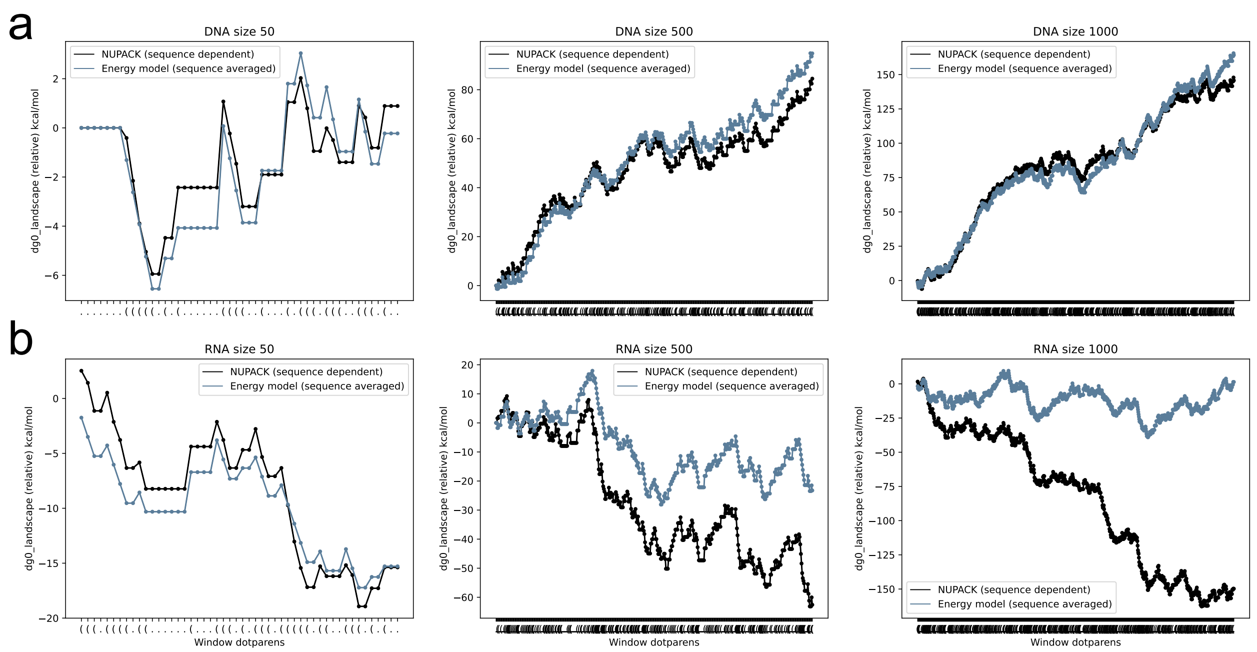


Supplementary Figure 2: Validity of the energy landscape approximation. Plots show the energy landscape created by sequence-averaged energy model (blue) versus the “true” sequence-dependent energy landscape created base-by-base using NUPACK (black) for helices of random sequence obeying the dot parens pattern on the x-axis. Both a) DNA and b) RNA windows are shown in three sizes. Agreement is fair over short window sizes (50nt). As window size increases agreement for DNA remains good whereas RNA diverges. However, in the latter case, the approximate landscape still retains the same profile features, including binding sites (downward dips) even if the landscape energy diverges.

- Landscape energy within interior loops remains constant, in line with the ’no stacking’ ensemble in NUPACK.
- Sequence-averaged energy landscape construction is 50-600 times faster than constructing an equivalent sequence-dependent energy landscape base-by-base using NUPACK (from windows of length 50-1000nt respectively). Figure 2 shows the approximation is good for DNA.

#### Binding Site Identification

The energy of a binding site between points $i$ and $j$ of the energy landscape (inclusive) is calculated as:

$$\Delta G^0_{\text{site}} = \Delta G^0_{\text{landscape}}(j) - \Delta G^0_{\text{landscape}}(i) + (\Delta G^0_{\text{assoc}} + \Delta G^0_{\text{bp}}) \tag{3}$$

and takes account of the entropy penalty in co-localising the (independent) strands when the first basepair is made.

Moving along an energy landscape, a unique binding site is considered to exist between a basepair at position $i$ and a local minimum on the energy landscape at position $j > i$ when:

1. The minimum at position $j$ is the *deepest* minimum on the energy landscape for which 2 and 3 below hold

2. At no point $x$ along the binding site $i < x < j$ does the landscape free energy equal or exceed the energy of the initial point $i$:

$$\Delta G^0(i < x < j) < \Delta G^0(i) \quad \forall x \tag{4}$$

3. The free energy of the binding site between $i$ and $j$ is equal to or less than -7 kcal mol$^{-1}$ at 37C:

$$\Delta G^0_{\mathrm{site}} \leq -7.0\,\mathrm{kcal\,mol^{-1}\ at\ 37C} \tag{5}$$

Once a binding site is found, the search for the next binding site begins at $i = j + 1$. Binding sites in the same energy landscape do not overlap, but binding sites belonging to different sliding window positions (different energy landscapes) can overlap.

Note that:

- Purple dots in Supplementary Figure 1(a) denote local minima along the energy landscape.
- Practically, Criterion 2 above means that if, for example, a short region of hybridisation is followed by a region of many interior loops, which are in turn followed by a long region of perfect complementary hybridisation (giving a deep energy minimum on the landscape), then the two regions of hybridisation are still classed as independent binding sites (not one long contiguous binding site).
- The energy landscape can also be constructed in the opposite direction (scaffold 3' to 5' direction) and the same binding sites are identified.
- For efficiency, if the maximum energy difference along the energy landscape is insufficient for a binding site, i.e. when

$$\Delta G^0_{\mathrm{landscape}}(\mathrm{min}) - \Delta G^0_{\mathrm{landscape}}(\mathrm{max}) + (\Delta G^0_{\mathrm{assoc}} + \Delta G^0_{\mathrm{bp}}) > -7.0\,\mathrm{kcal\,mol^{-1}}$$

  then no binding sites are checked for and the next sliding window region is moved to immediately.

#### NUPACK Sampling of Parameters

The $\Delta G^0_{\mathrm{bp}}$ parameters and the $\Delta G^0_{\mathrm{sym\ internal\ loop}}(l)$ lookup function were obtained by sampling the NUPACK v4.0.0.27 (Ref [? ]) `structure_energy()` function.

$\Delta G^0_{\mathrm{bp}}$ was obtained by sampling the free energy of a perfectly complementary random helix of DNA or RNA (G-U wobble pairs were included as complements in RNA). The helix was set to 50bp and then incremented to 100bp in 1bp steps. At each length, the helix hybridisation energy was calculated. $\Delta G^0_{\mathrm{bp}}$ was calculated as the average energy *difference* between 1bp increments of the helix at 37C (note that this difference calculation allowed convenient cancellation of the helix initiation energy). This was repeated for 500 helices to gain a total of $n = 25000$ samples. $\Delta G^0_{\mathrm{bp}}$ was set to the sample mean and yielded a notably higher value for RNA than for DNA (Supplementary Table 1).

The $\Delta G^0_{\mathrm{sym\ internal\ loop}}(l)$ lookup function was obtained by creating a perfectly complementary random helix of DNA or RNA between 6bp and 65bp. The free energy of the helix was recorded as $\Delta G^0_{\mathrm{no\ loop}}$ at 37C. Then a random insertion point for a symmetric interior loop was chosen after the first base pair and before the last base pair of the helix. A loop of a single basepair mismatch was added to the helix and the free energy of the complex recorded as $\Delta G^0_{\mathrm{loop}}(1)$ at 37C. The loop penalty for a length 1 symmetric interior loop was calculated as $\Delta G^0_{\mathrm{loop}}(1) - \Delta G^0_{\mathrm{no\ loop}}$. The symmetric interior loop was then incremented from 1 to 30 mismatching basepairs and the loop penalty calculated at each loop length.

| Parameter | Description | Value | Unit |
|---|---|---|---|
| $\Delta G^0_{\text{bp}}$ (DNA, 37C) | Per-basepair sequence-averaged hybridisation energy DNA | -1.31 | kcal mol$^{-1}$ |
| $\Delta G^0_{\text{bp}}$ (RNA, 37C) | Per-basepair sequence-averaged hybridisation energy RNA | -1.75 | kcal mol$^{-1}$ |
| $\Delta G^0_{\text{assoc}}$ (37C) | Entropic penalty for two strands co-localising | 1.619 | kcal mol$^{-1}$ |
| $\Delta S^0_{\text{loop}}$ | Loop entropy | -0.0228567 | kcal K$^{-1}$ mol$^{-1}$ |
| $\gamma$ | Loop exponent | 2.0 | |

Supplementary Table 1: Energy model parameters for for metrics $M_1$ and $M_2$. Note that for RNA $\Delta G^0_{\text{bp}}$ (RNA, 37C) includes G-U wobble pairs. For RNA, during sampling, every G base was paired with C 50% of the time and U 50% of the time. Every U base was paired with A 50% of the time and G 50% of the time.

This was repeated for 500 helices, to gain a total of $n = 500$ samples of the energy penalty of each loop length. $\Delta G^0_{\text{sym internal loop}}(l)$ was set to the sample mean at each loop length $l$. Supplementary Figure 1(b) and (c) shows that the loop penalties for DNA and RNA helices are similar. Parameters $\Delta S_{\text{loop}}$ and $\gamma$ for the Jacobson-Stockmayer approximation (2) were found by manually fitting the data.

#### Energy Model General Notes

1. The energy landscape assumes that two strands are paired (or mismatched) base-for-base in the sliding window region. Thus, it considers that only base pairings and symmetric interior loops exist. Conceivably, a staple could also hybridise to the scaffold with a binding site also containing small bulge loops and/or an asymmetric interior loops. However, these latter cases (which would require more complex sequence alignments) are ignored for the benefit of speed: the aim here is to perform a fast and approximate identification of possible binding sites where some basic mismatching is taken into account.

2. Binding energies returned by the energy model are sequence-averaged and also ignore dangle and co-axial stacking contributions. However, in Metrics $M_1$ and $M_2$, energy values are summed and cumulative energy values are only ever compared *relatively*. Hybridisation energies are not used in a absolute sense to e.g. set reaction equilibrium constants, and thus approximated energies are sufficient.

3. The energy model assumes that both participating strands in a sliding window region have no internal secondary structure. In some cases, secondary structure in the sliding window strand pair could make some binding sites energetically non-viable. Therefore, off-target binding sites detected by the energy model should be considered as off-target binding sites in the *worst case*.

4. RNA has a stronger $\Delta G^0_{\text{bp}}$ than does DNA, but has similar interior loop penalty costs. Also, RNA has the additional G-U complementary pair. These factors mean that the energy landscape for RNA-RNA bindings has a downward, rather than upward trend (Figure 2), and the RNA energy model thus tends to detect more binding sites and longer binding sites as compared to the DNA energy model.

5. Even though detected off-target binding sites are labelled as 'beginning' at basepair *i* and 'ending' at basepair *j*, off-target binding sites are typically *fuzzy objects* without sharply defined boundaries. In most cases, an off-target site features one or more nucleation centres where an incoming strand can start to bind. Once bound, the incoming strand can continuously hybridise

and melt along the extent of the site in a dynamic manner. This is in contrast to **on-target** binding sites where: (i) runs of complementary bases are flanked by runs of base mismatches and (ii) hybridisation can be approximated with a two-part model of binding.

### Supplementary Note 2 Definitions of Metrics 1, 2, 3, 4

Note that all metrics defined below take the *absolute value* of the computed free energy quantity. This allows the metrics to be costs, which are minimised to zero from a positive number. Metrics 1 and 2 derive sequence-averaged free energies whereas Metrics 3 and 4 derive sequence-dependent free energies.

#### Metric 1

***Premise:*** *The yield of the target origami is more likely be maximised if potential off-target staple binding positions on the scaffold are minimised, leaving only on-target staple binding positions.*

Metric 1 indicates to what extent staples can initially bind the scaffold in non-designed i.e. "off-target" locations. Metric 1 is the (absolute) total free energy of initial off-target staple binding locations on the scaffold.

It should be emphasised that Metric 1 is concerned with minimising the number of *initial* off-target binding sites where staples *first* hybridise with the scaffold: it is not concerned with calculating and minimising binding sites for the second, third, fourth etc. staple legs, which are additionally determined by entropic loop penalties dependent on the current fold state of the structure.

Minimising Metric 1 reduces potential kinetic traps in origami self-assembly by simultaneously minimising both (i) the number and (ii) the strength of initial off-target binding sites.

Metric 1 is defined as:

$$M_1 = |\sum_{s_i \in \mathcal{S}} \min\left(0, \Delta G^0_{\text{offtarget}}(s_i)\right)| \qquad \text{optimised as} \quad M_1 \to 0 \tag{6}$$

where $\mathcal{S}$ is the set of staples $\{s_1, s_2, ...s_n\}$ and

$$\Delta G^0_{\text{offtarget}}(s_i) = \Delta G^0_{\text{total}}(s_i) - \Delta G^0_{\text{ontarget}}(s_i) \tag{7}$$

is the total free energy of initial off-target binding sites for staple $s_i$ on the scaffold. Defining the terms in Eq. (7):

- $\Delta G^0_{\text{total}}(s_i)$ is the total free energy of all initial **on-target** and **off-target** binding sites for staple $s_i$ on the scaffold. It is calculated algorithmically by positioning staple $s$ anti-parallel to the scaffold and moving it along the scaffold as a "sliding window" as indicated in Supplementary Figure 3. At each position of the sliding window the energy of all potential binding sites between the staple bases and the aligned scaffold bases is calculated by the energy model, as explained in Supplementary Note 1. $\Delta G^0_{\text{total}}(s_i)$ is the sum of all binding site energies when the staple window has been moved all the way along the scaffold.

- $\Delta G^0_{\text{ontarget}}(s_i)$ is the total free energy of the initial **on-target designed binding sites** for staple $s_i$ on the scaffold. It is the sum of the (sequence averaged) hybridisation energies for each of the individual staple leg binding sites, namely:

$$\Delta G^0_{\text{ontarget}}(s_i) = d_{s_i} \Delta G^0_{\text{init}} + b_{s_i} \Delta G^0_{\text{bp}} \tag{8}$$

  where $d_{s_i}$ is the number of staple legs that $s_i$ binds to the scaffold, and $b_{s_i}$ is the total number of bases on staple $s_i$ that hybridise to the scaffold. This is derived by considering that each staple section initially binds with energy $\Delta G^0_{\text{init}} + b\Delta G^0_{\text{bp}}$, where $b$ is the number of bases on the staple section in question.

#### Metric 1 Notes

- When all initial staple binding sites detected are exactly the designed staple binding sites, then $\Delta G^0_{\text{total}}(s_i) = \Delta G^0_{\text{ontarget}}(s_i)$ and thus $\Delta G^0_{\text{offtarget}}(s_i) = 0$
- The most common case is for extra off-target binding sites exist for a staple. Then $\Delta G^0_{\text{total}}(s_i) < \Delta G^0_{\text{ontarget}}(s_i)$ and thus $\Delta G^0_{\text{offtarget}}(s_i) < 0$
- Very short staple sections (e.g. less than 6nt) are **not detectable by the energy model** and therefore are not included in $\Delta G^0_{\text{total}}(s)$. However, they are included in $\Delta G^0_{\text{ontarget}}(s_i)$, making it more negative. Staple sections less than 6nt therefore have the effect of making $\Delta G^0_{\text{offtarget}}(s_i)$ less negative overall and make it seem (falsely) like less off-target sites exist.
- The $\min$ function is included for completeness, to ensure that $\Delta G^0_{\text{offtarget}}(s_i)$ never becomes positive in the rare case that an origami design has most staples with very short sections.
- Extended binding regions "spilling over" designed binding sites also (correctly) contribute to the total energy of off-target binding sites.

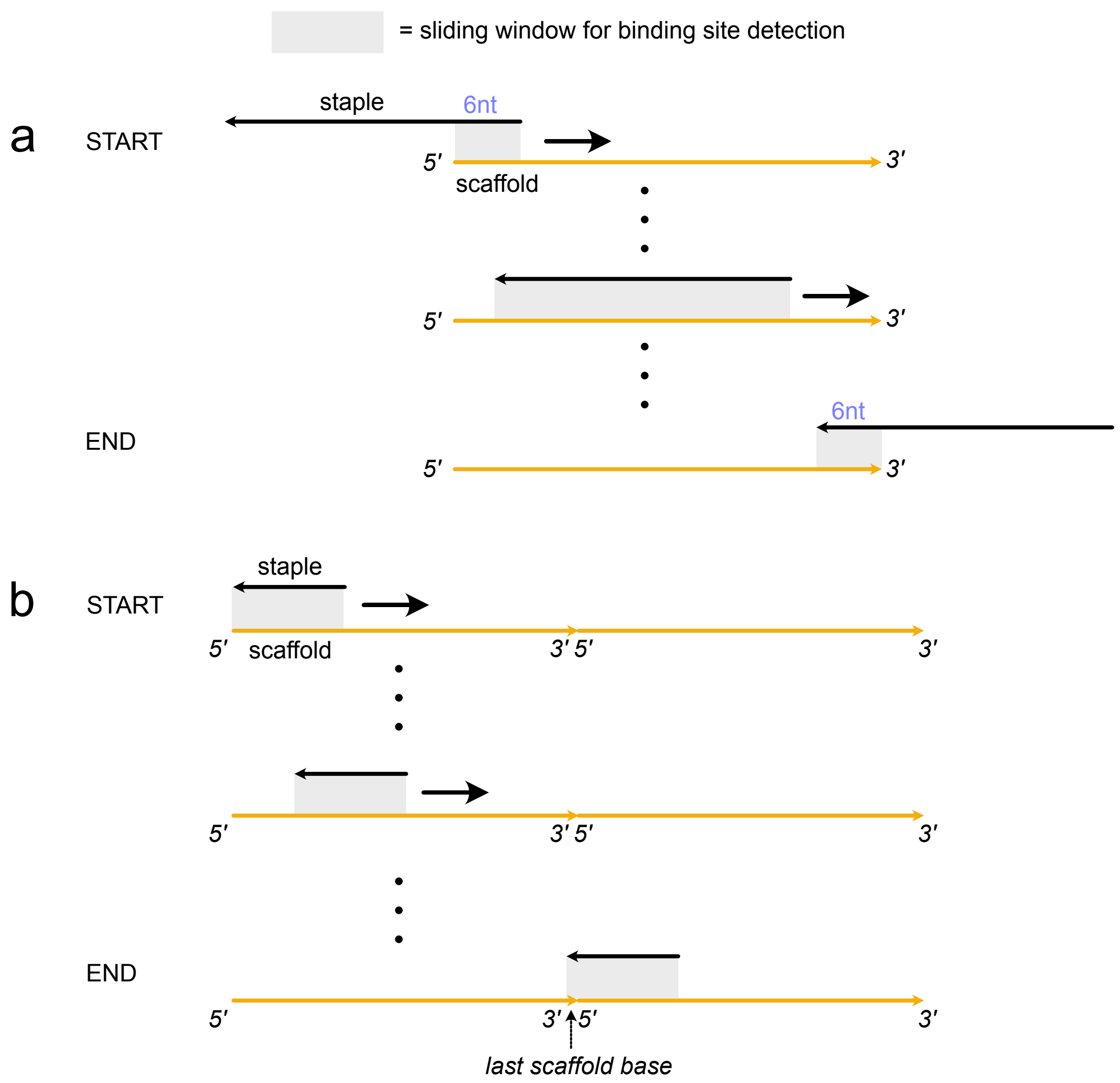


Supplementary Figure 3: Metric 1 sliding staple window for (a) linear and (b) circular scaffolds. For linear scaffolds (a) a staple always overlaps by at least 6 bases; windows smaller than this are not computed. In (b), the flanking scaffold to the right is a repeat of the scaffold to the left (circular boundary conditions). Window size for a linear scaffold is variable at the beginning and end; window size for a circular scaffold is constant.

#### Metric 2

**Premise:** *The yield of the target origami is more likely be maximised when potential scaffold-scaffold binding sites are minimised, as such sites can slow or prevent staple hybridisation by partially or completely blocking staple binding sites.*

Metric 2 computes the prevalence of scaffold-scaffold binding sites. The scaffold is checked for pairing regions systematically, following a binding permutation scheme explained in Supplementary Figure 4. The binding permutation schemes essentially ensure that each subsequence of length $N$ on the scaffold is aligned against every other non-overlapping subsequence of length $N$, at some point.

All scaffold-scaffold binding sites detected are off-target; ideally none would exist. Metric 2 is the (absolute) total energy of scaffold-scaffold binding sites found in all of the distinct binding permutations $\mathcal{P}$.

Metric 2 is defined as:

$$M_2 = |\sum_{p_i \in \mathcal{P}} \Delta G^0_{\text{total}}(p_i)| \qquad \text{optimised as} \quad M_2 \to 0 \tag{9}$$

where $\Delta G^0_{\text{total}}(p_i)$ is the cumulative energy of off-target binding sites in permutation $p_i$ calculated by the energy model (Supplementary Note 1).

#### Metric 2 Notes

- Metric 2 approximates the total energy of all the scaffold-scaffold binding sites, in the absence of loops. Scaffold-scaffold binding events are effectively treated as bi-molecular events where two independent strands come together in solution. In reality, scaffold-scaffold bindings are pseudo-unimolecular reaction events. Instead of a constant initiation penalty, scaffold-scaffold bindings have an entropic loop penalty dependent on the loop size between the hybridising domains (which is, in turn, dependent on the temporal fold state of the origami). Hence, it should be emphasised that Metric 2 **does not** calculate quantitatively accurate hybridisation energies between different complementary regions of the scaffold. Rather, a lower value of Metric 2 simply indicates that one scaffold has *relatively less* off-target scaffold-scaffold binding sites than another scaffold. This is sufficient for the multi-objective selection approach in this work.

- Metric 2 is more suitable than a direct MFE calculation of the scaffold sequence. This is because Metric 2 includes **all** possible scaffold-scaffold binding sites, whereas the MFE only includes scaffold-scaffold bindings in the MFE configuration. Furthermore, MFE calculation time for long scaffolds is considerable in NUPACK and circular scaffolds and pseudo-knots are prohibited.

- Metric 2 also serves to quantify the extent of inter-scaffold bindings, i.e. off-target bindings between two (or more) different copies of the scaffold strand in solution.

- Circular and linear versions of the same scaffold generally have identical $M_2$ scores, except when the circular version contains extra binding sites existing in the region where the virtual 5' and 3' scaffold ends join into a loop.

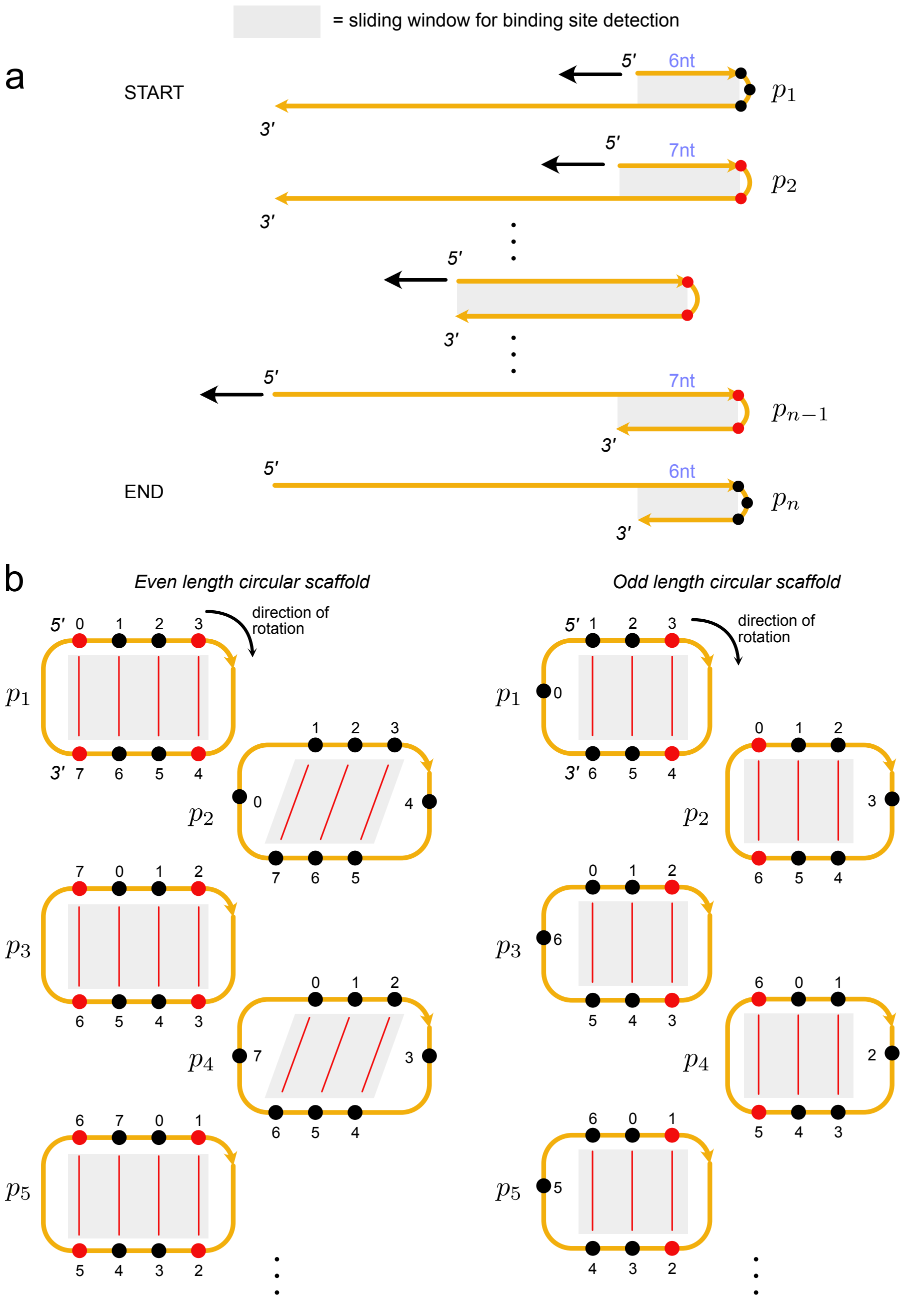


Supplementary Figure 4: Permutation schemes to find all possible scaffold-scaffold binding regions in Metric 2. Scheme for **(a)** linear scaffolds and **(b)** circular scaffolds. Each subsequent configuration in (a) and (b) is a named 'binding permutation', denoted $p_i$. Circular scaffolds have a different calculation scheme to ensure that potential binding sites spanning across the virtual 5' and 3' ends of the scaffold sequence are accounted for. In (a) the 5' end of the scaffold (top strand) is "pulled backwards" (black arrow) from an initial minimum overlap of 6nt and the scaffold rotates until the bottom strand "shrinks" to a minimum overlap of 6bp. Alternate rotations feature one base missed out at the end of the loop; this is necessary to include all possible basepair-basepair bindings. Red circles denote adjacent bases that cannot physically hybridise due to the scaffold back bone. However, for simplicity these bases are assumed able to hybridise. In (b) different schemes are employed depending on whether the circular scaffold is even or odd length. In each case, the scaffold is rotated through alternate full and half turns. Note that each circular permutation in (b) can be rotated 180 degrees to yield to a permutation which is rotationally equivalent. For efficiency, rotationally equivalent permutations are not double counted.

#### Metric 3

***Premise:*** *The yield of the target origami is more likely be maximised if staples are able to engage in hybridisation reactions with the scaffold as singular entities, rather than being sequestered in strong staple-staple co-folds or in multi-stranded staple complexes.*

Metric 3 calculates the worst extent of staple-staple co-folding in the staple set. It is reasoned that if staples are less likely to co-fold in pairs then multi-stranded staple complexes ($\geq$ 3 strands) are also less likely. Metric 3 is the (absolute) energy of the strongest staple-staple co-fold at 55C as assessed by NUPACK `mfe()`. See Table 2 for NUPACK parameters. Minimising Metric 3 brings down the ceiling value for all staple-staple co-fold MFE energies at 55C.

The number of distinct staple-staple pairs in the set of all co-folds $\mathcal{C}$ is given by:

$$|\mathcal{C}| = \frac{1}{2}|\mathcal{S}|\left(|\mathcal{S}| + 1\right) \tag{10}$$

where where $|\mathcal{S}|$ denotes the total number of staples. Note that all staple dimers $(s_n, s_n)$ are included in $\mathcal{C}$.

Metric 3 is defined as:

$$M_3 = \left|\min\left(\Delta G^0_{s_1,s_2} - \Delta G^0_{s_1} - \Delta G^0_{s_2} \quad \forall (s_1, s_2) \in \mathcal{C}\right)\right| \qquad \text{optimised as} \quad M_3 \rightarrow 0 \tag{11}$$

where $\Delta G^0_{s_1,s_2}$ is the minimum free energy of formation of staple co-fold $(s_1, s_2)$ at 55C and $\Delta G^0_{s_n}$ is the minimum free energy of formation of staple $s_n$ at 55C. Note that term $\Delta G^0_{s_1,s_2} - \Delta G^0_{s_1} - \Delta G^0_{s_2}$ gives the overall free energy of binding of the two staple strands, taking into account the secondary structure of the individual staples themselves. Staple pairs which are completely orthogonal (with no co-fold MFE structure defined) are omitted from the minimum calculation in (11). For computational efficiency, Metric 4 is calculated before Metric 3 and, once computed, $\Delta G^0_{s_n}$ values are saved for re-use by Metric 3.

#### Metric 3 Notes

- Staple co-fold binding energies are dependent on three energies $\Delta G^0_{s_1,s_2}$, $\Delta G^0_{s_1}$ and $\Delta G^0_{s_2}$, each with their own temperature dependence. The choice is made to evaluate $M_3$ at 55C in the middle of the origami folding temperature range.

- Staple concentrations are also relevant for the formation of staple complexes. Metric 3 works on the assumption that minimising staple co-fold energies minimises the likelihood of staple complexes at any given concentration.

- If some staples are intended to hybridise together as a feature of the origami design, then these hybridisation regions should be removed from the staples before Metric 3 is run.

| Staple Material | Na (M) | Mg (M) | Ensemble | Temperature (C) |
|---|---|---|---|---|
| DNA | 0.05 | 0.0125 | No Stacking | 55 |
| RNA | 1.0 | 0 | No Stacking | 55 |

Supplementary Table 2: NUPACK `mfe()` parameters for metrics $M_3$ and $M_4$. DNA parameters attempt to mimic a typical origami folding buffer (12.5mM magnesium). RNA parameters are constrained to 1.0M sodium by lack of RNA salt corrections in NUPACK. The "No Stacking" ensemble disregards energetic contributions from dangling ends and co-axial stacking.

#### Metric 4

***Premise:*** *The yield of the target origami is more likely be maximised when individual staples possess less internal secondary structure as internal base pairing can sequester staple bases designed to hybridise with the scaffold.*

Metric 4 calculates the worst extent of intra-staple secondary structure in the staple set.

Metric 4 is the (absolute) energy of the staple with the strongest hairpin-like fold at 55C, as assessed by NUPACK `mfe()`. Minimising Metric 4 minimises the worst-case hairpin fold in the staple set. An evaluation temperature of 55C is used for consistency with Metric 3.

Metric 4 is defined as:

$$M_4 = \left|\min\left(\Delta G^0_{s_i} \quad \forall s_i \in \mathcal{S}\right)\right| \qquad \text{optimised as} \quad M_4 \to 0 \tag{12}$$

where $\Delta G^0_{s_i}$ is the minimum free energy of formation of staple $s_i$ at 55C and $\mathcal{S}$ is the set of staples.

#### Filter $G$

Filter $G$ is not a metric, but may help in deciding which sequence set to order. It counts the number of staples on an Pareto front origami which contain four consecutive G's in their sequence. Such motifs may lead to stacked G-quadruplex structures which can cause difficulty in staple synthesis (reduced yields) and in determination of staple purity. A lower $G$ score is preferable, ideally $G = 0$.

### Supplementary Note 3 Pareto Front Multi-Criteria Decision Making

When a Pareto front of optimal trade-off solutions has been computed in the objective space defined by the metrics in Supplementary Note 2, it is necessary to select a **single** candidate solution. Defined below are three different multi-criteria decision making (MCDM) schemes we used to rank individuals on the Pareto front.

We note that the solution selected from the Pareto front by an MCDM method is influenced by the choice of normalisation scheme used [1]. We used a simple max-min normalisation method and considered the range of each metric across the whole of objective space, not just across the Pareto front region.

#### Normalised and Weighted Decision Matrix

The Pareto front of optimal trade-off solutions in objective space is first written as a decision matrix $\mathbf{X} = (x_{ij})_{m\times n}$ where the $m$ rows represent individuals $i$ on the Pareto front and the $n$ columns give the metric scores of each individual:

| Metrics | $M_1$ | $M_2$ | $M_3$ | $M_4$ |
|---|---|---|---|---|
| | $j = 1$ | $j = 2$ | $j = 3$ | $j = 4 = n$ |
| Alternatives | | | | |
| $i = 1$ | $x_{ij}$ | | | |
| $i = 2$ | | | | |
| $i = 3$ | | | | |
| $i = 4$ | | | | |
| $i = m$ | | | | |

Note: Each element $x_{ij}$ is rounded to 8dp to prevent ranking differences based on minor numerical imprecisions.

The decision matrix is then **normalised** to place all metrics on comparable absolute scales. Each element $x_{ij}$ in the decision matrix is normalised into range $0 \leq r_{ij} \leq 1$ by a linear "Max-Min" normalisation:

$$r_{ij} = \frac{x_{ij} - M_j^{\min}}{M_j^{\max} - M_j^{\min}} \tag{13}$$

whereby each metric score is converted to a fraction of the entire metric score range. In the above $M_j^{\max}$ and $M_j^{\min}$ are the maximum and minimum values of metric $M_j$ respectively, in objective space. When $x_{ij} = M_j^{\min}$ then $r_{ij} = 0$. When $x_{ij} = M_j^{\max}$ then $r_{ij} = 1$. This normalisation method still works when some (but not all) of the elements in a column $j$ of the decision matrix are zero (a possibility for small origamis).

Every metric $M_j$ is assigned a weight $0 \leq w_j < 1$. In this work, all used metrics have equal weights and their weights sum to 1. Unused metrics have a weight of 0. Finally, each element of the normalised decision matrix is multiplied by the weight of the metric to which it pertains, giving the **normalised and weighted decision matrix** $\mathbf{V}$ where each element:

$$v_{ij} = r_{ij} w_j \tag{14}$$

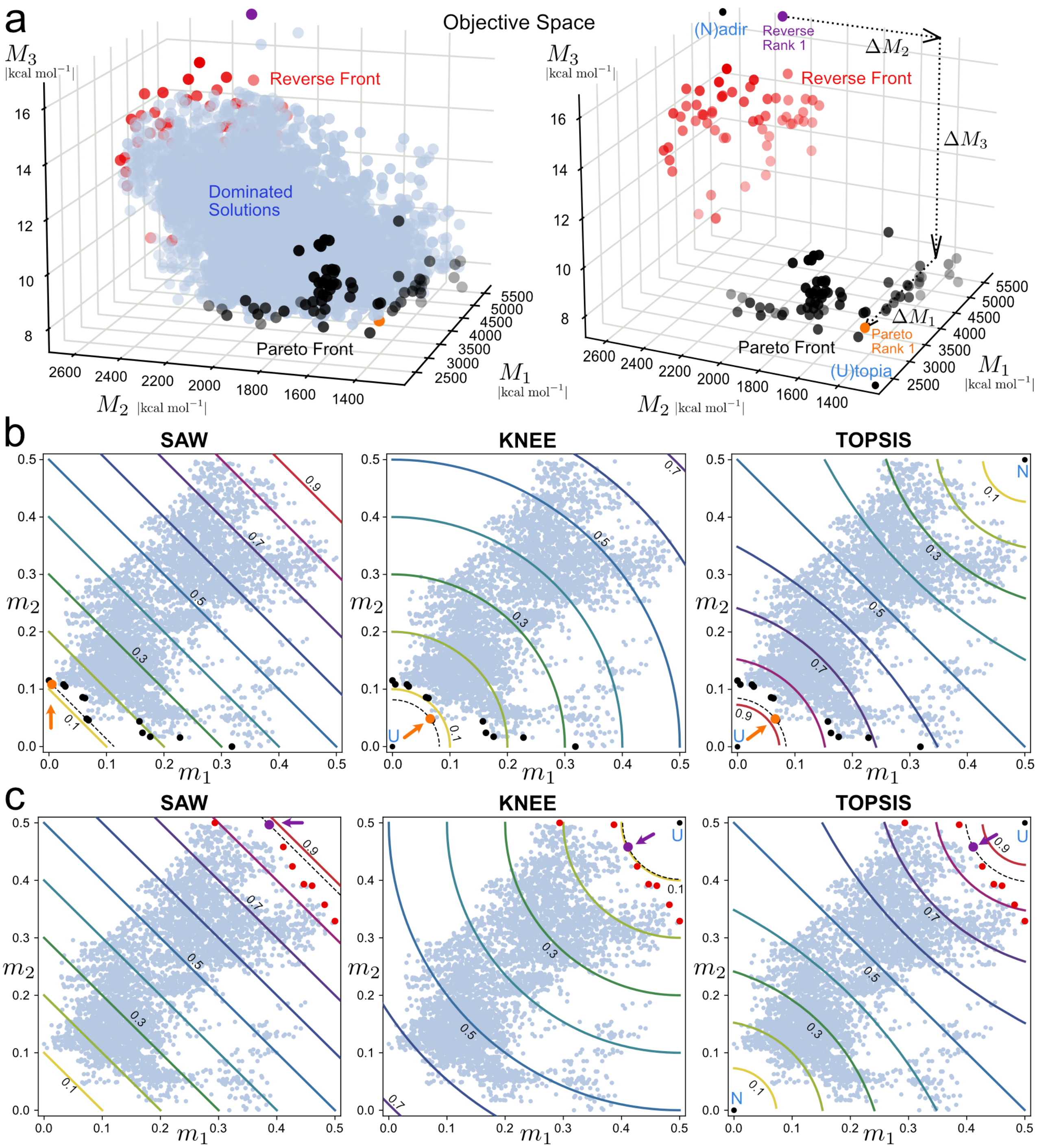


Supplementary Figure 5: Graphical illustration of different MCDM Methods. **(a)** Scoring an origami leads to an objective space of points, where the Pareto front solutions (black dots) and reverse front solutions (red dots) can be identified. Note metric $M_4$ is omitted to give 3 dimensions. Right panel omits dominated solutions to show Pareto and reverse fronts more clearly. **(b)** and **(c)** Reduction of objective space to two dimensions (only $M_1$ and $M_2$) and scaling so that metric scores are normalised and equally weighted (becoming $m_1$ and $m_2$ respectively). The simplification to 2 dimensions allows to super-impose the scoring lines of the different MCDM methods and to graphically illustrate how Pareto front candidate selection differs between these methods: (b) illustrates which candidate (orange) each method selects from the Pareto front (black dots); (c) illustrates which candidate (purple) each method selects from the reverse front (red dots). U denotes utopia point, N denotes nadir point. In practice, the MCDM methods operate in 3- or 4-dimensional space.

In matrix $\mathbf{V}$, we denote $\mathbf{v}_j$ as the column vector of scores for metric $j$.

In Supplementary Figure 5 panels (b) and (c), the $m_1$ and $m_2$ variables comes from matrix $\mathbf{V}$: they are the normalised and weighted versions of the $M_1$ and $M_2$ scores.

#### SAW

The SAW (Simple Additive Weighting) method ranks Pareto candidates simply by the weighted sum of their normalised metric scores.

##### Ranking of Pareto and Reverse Fronts

The SAW score of Pareto individual $i$ is the sum of row $i$ in the normalised and weighted decision matrix $\mathbf{V}$:

$$\mathrm{SAW}(i) = \sum_{j=1}^{n} v_{ij} \tag{15}$$

- The top ranked individual on the Pareto front has the **lowest** SAW score.
- The top ranked individual on the reverse front (the worst solution) has the **highest** SAW score.

Graphed in two dimensions (e.g. with 2 metrics) SAW is a series of linear contour lines across normalised-weighted objective space (Supplementary Figure 5).

#### KNEE

The KNEE method ranks Pareto candidates by their proximity to a "utopia" point U. The shorter the euclidean distance to the utopia point, the lower (better) the KNEE score. As Ref [2] comment, "The knee point [the solution closest to the utopia point] is furthest from the extreme points of the Pareto front and represents a good compromise among the various objectives".

##### Ranking of Pareto Front

For the Pareto front, the utopia point is at the 0 origin in normalised-weighted objective space. The KNEE score of Pareto individual $i$ is simply the euclidean distance:

$$\mathrm{KNEE}(i) = \sqrt{\sum_{j=1}^{n} (v_{ij} - 0)^2} = \sqrt{\sum_{j=1}^{n} (v_{ij})^2} \tag{16}$$

- The top ranked individual on the Pareto front has the **lowest** KNEE score according to Equation 16.

Graphed in two dimensions (e.g. with 2 metrics), KNEE is a series of circular arc contour lines across normalised-weighted objective space (Supplementary Figure 5).

##### Ranking of Reverse Front

For the reverse front, the utopia point U is instead defined as where all metrics have their maximal normalised-weighted score. The KNEE score of reverse front individual $i$ is thus:

$$\mathrm{KNEE}(i) = \sqrt{\sum_{j=1}^{n} (v_{ij} - \max(\mathbf{v}_j))^2} \tag{17}$$

- The top ranked individual on the reverse front (the worst solution) has the **lowest** KNEE score according to Equation 17.

#### TOPSIS

The TOPSIS (Technique of Order Preference by Similarity to Ideal Solution) method is similar to the KNEE method. However, whereas the KNEE method just considers proximity of a Pareto candidate to a single utopia point, the TOPSIS method also considers proximity to an undesirable "nadir" point. The score is maximised if a Pareto candidate is both close the the utopia point **and** far from the nadir point.

##### Ranking of Pareto Front

A Pareto candidate $i$ has the following euclidean distance to the 0 origin utopia point:

$$D_i^{\mathrm{utopia}} = \sqrt{\sum_{j=1}^{n} (v_{ij})^2} \tag{18}$$

and the following euclidean distance from the nadir point:

$$D_i^{\mathrm{nadir}} = \sqrt{\sum_{j=1}^{n} (v_{ij} - \max(\mathbf{v}_j))^2} \tag{19}$$

The TOPSIS score is the fraction that the nadir distance is with respect to the total distance to the utopia and nadir points:

$$\mathrm{TOPSIS}(i) = \frac{D_i^{\mathrm{nadir}}}{D_i^{\mathrm{utopia}} + D_i^{\mathrm{nadir}}} \tag{20}$$

When a Pareto candidate is close to the utopia point and far from the nadir point the fraction is dominated by the nadir distance and $\mathrm{TOPSIS}(i) \approx D_i^{\mathrm{nadir}}/D_i^{\mathrm{nadir}} = 1$. Conversely, when far from the utopia point and close to the nadir point then utopia distance dominates and $\mathrm{TOPSIS}(i) \approx 0/D_i^{\mathrm{utopia}} = 0$.

- The top ranked individual on the Pareto front has the **highest** TOPSIS score according to Equation 20.

Graphed in two dimensions (e.g. with 2 metrics), TOPSIS appears as a set of contours resembling magnetic dipole field lines encircling the utopia and nadir points in normalised-weighted objective space (Supplementary Figure 5).

##### Ranking of Reverse Front

The top ranking TOPSIS individual on the reverse front (worst solution) is calculated by swapping the utopia and nadir points around, such that:

$$D_i^{\text{utopia}} = \sqrt{\sum_{j=1}^{n} (v_{ij} - \max(\mathbf{v}_j))^2} \tag{21}$$

$$D_i^{\text{nadir}} = \sqrt{\sum_{j=1}^{n} (v_{ij})^2} \tag{22}$$

and again using Equation 20 to calculate the TOPSIS score for all candidates $i$.

- The top ranked individual on the reverse front (the worst solution) has the **highest** TOPSIS score according to Equation 20 when the utopia and nadir points are defined as in Equations 21 and 22 respectively

### Part II

### Computational Results

In this part, we perform scaffold sequence selection for 14 different DNA origami shapes ranging from 342nt scaffold (7 staples) to 8052nt scaffold (199 staples). The DNA origamis include 2D and 3D shapes in both wireframe and raster designs. For each origami shape, we performed scaffold sequence selection from four qualitatively different sequence pools (described in the main paper) using an HPC computing cluster.

#### Supplementary Note 4 Test Set of 14 DNA Origamis

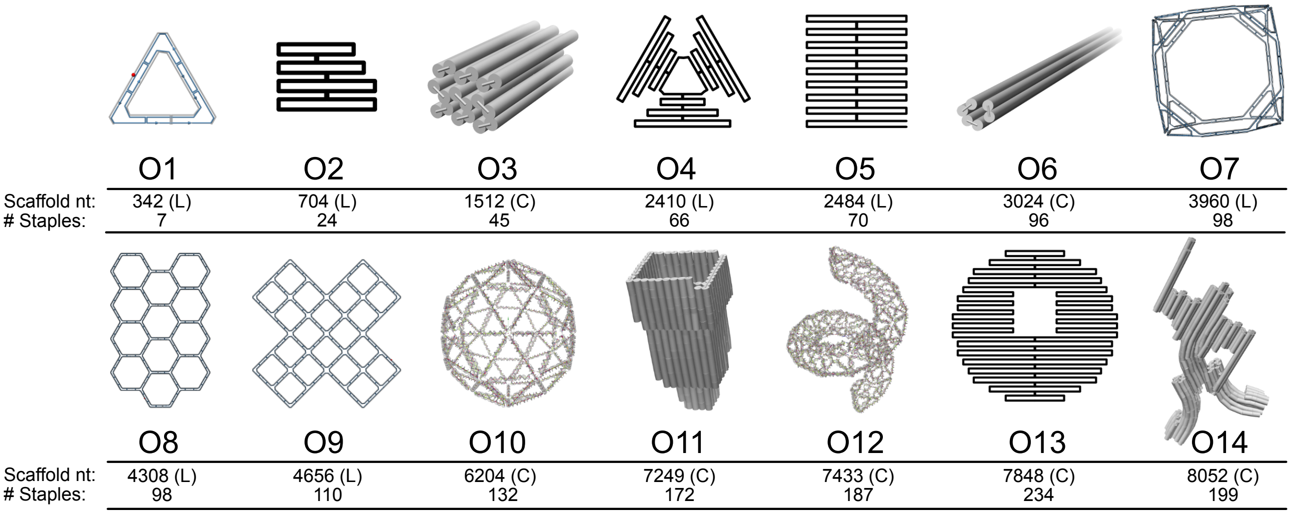


| | Origami | Ref | Scaffold | | | | Staples | | | | | | Warnings |
|---|---|---|---|---|---|---|---|---|---|---|---|---|---|
| | | | A | B | C | D | E | F | G | H | I | J | <7nt |
| O1 | Tiny Triangle | [3] | M13mp18 | **342** | L | 27 | **7** | 0/1/1/5 | 7 | 21→71 | 48.43 | 24 | 2 |
| O2 | Four Finger Tile | [4] | M1.3 | **704** | L | 0 | **24** | 2/13/8/1 | 4 | 18→42 | 29.33 | 0 | 1 |
| O3 | ABrick | [5] | pScaf_1512 | **1512**$^{(N1)}$ | C | 0 | **45** | 0/1/7/37 | 6 | 21→49 | 36.8 | 144 | 10 |
| O4 | Small Triangle | [6] | Mini M13 | **2410** | L | 178 | **66** | 2/13/45/6 | 4 | 20→44 | 34.18 | 24 | 0 |
| O5 | Small Rectangle | [7] | Synthetic DBS | **2484** | L | 0 | **70** | 2/20/48/0 | 3 | 29→48 | 35.49 | 0 | 0 |
| O6 | 6-Helix Bundle | [5] | pScaf_3024 | **3024**$^{(N1)}$ | C | 0 | **96** | 0/23/43/30 | 5 | 20→49 | 31.98 | 46 | 6 |
| O7 | Truncated Cube | [3] | M13mp18 | **3960** | L | 120 | **98** | 0/0/39/59 | 6 | 22→59 | 41.88 | 264 | 0 |
| O8 | Hexagonal Tile | [3] | M13mp18 | **4308** | L | 85 | **98** | 0/0/47/51 | 10 | 32→72 | 46.31 | 315 | 44 |
| O9 | Cross Mesh | [3] | M13mp18 | **4656** | L | 38 | **110** | 0/2/33/75 | 7 | 20→65 | 46.13 | 456 | 48 |
| O10 | Ball | [8] | M13mp18 | **6204**$^{(N2)}$ | C | 0 | **132** | 0/132/0/0 | 2 | 42→50 | 47 | 0 | 0 |
| O11 | Nanopore | [9] | M13mp18 | **7249** | C | 1174 | **172** | 1/39/47/85 | 6 | 18→49 | 35.32 | 0 | 55 |
| O12 | Helix | [8] | p7560 | **7433**$^{(N2)}$ | C | 165 | **187** | 0/79/31/77 | 8 | 26→57 | 39.44 | 107 | 7 |
| O13 | Small Moon | [10] | p8064 | **7848**$^{(N2)}$ | C | 821 | **234** | 10/33/191/0 | 3 | 8→40 | 30.03 | 0 | 8 |
| O14 | Robot | [11] | p8064 | **8052**$^{(N2)}$ | C | 570 | **199** | 0/11/20/168 | 8 | 18→54 | 37.60 | 0 | 167 |

Supplementary Figure 6: 14 Origami Test Set.

**Table Columns:**
A = Scaffold origin in original publication
B = Scaffold length (nt)
C = Scaffold type (**L**inear or **C**ircular)
D = Number of non-hybridised bases on scaffold (nt)
E = Number of staples
F = Counts of staples with 1/2/3/3+ sections respectively
G = Maximum number of sections on a staple
H = Staple length range (nt)
I = Average staple length (nt)
J = Number of unhybridised bases on all staples, in single stranded loop-outs or dangling ends (nt)
<7nt = Number of staple sections binding to the scaffold which are smaller than 7nt (the energy model cannot detect these short bindings when material is DNA)

**Table Notes:**

(N1) pScaf is a custom scaffold sequence derived in Ref [5]. pScaf_1512 and pScaf_3024 sequences were obtained from https://github.com/douglaslab/pScaf.

(N2) Scaffold sequence is assumed to be first <scaffold length> bases of the sequence in column A.

### Supplementary Note 5 Scaffold Sequence Selection from Biological Vectors, de Bruijn and Random Sequences

Supplementary Figures 7, 8, 9 and 10 show the computational results for the 14 origami test set in Supplementary Note 4. Each figure shows the total variance in metrics 1,2,3 and 4, respectively, over the four scaffold sequence pools described in the main paper. Metrics $M_1$ to $M_4$ all have equal weightings for MCDM scoring.

On each of the figures, panel (a) shows the variance in off-target binding sites versus origami scaffold length, for each sequence pool type; panel (b) shows the relative percent decrease in off-target binding sites that can be achieved via scaffold selection, versus origami scaffold length, for each sequence pool type; panel (c) shows a direct comparison of data in panel (a).

Panel (a) for metrics 1 and 2 reports the **density** of off-target binding sites. This is metric score normalised by the scaffold length $L_{\text{scaf}}$. Using density allows cross-comparison of origamis with different scaffold lengths. The limits of each box plot show the minimum and maximum values off-target site density (|kcal / mol| per scaffold base) encountered over all samples; the coloured box section shows the variance of binding site density between the worst individual on the reverse front and the best individual on the Pareto front as selected by TOPSIS.

Panel (b) reports for all metrics the **relative change** in metric scores (binding site prevalence) between the best and worst candidates selected by the MCDM methods in Supplementary Note 3 (coloured lines). This plot is a measure of the effectiveness of scaffold selection; it shows by what relative percentage the presence of off-target binding sites can be reduced. Taking metric 1 (Supplementary Figure 7) as example, the coloured lines show the following relative percentage:

$$r = \frac{M_1^{\text{MCDM\#1,pareto}} - M_1^{\text{MCDM\#1,reverse}}}{M_1^{\text{MCDM\#1,reverse}}} \times 100 \tag{23}$$

where $M_1^{\text{MCDM\#1,pareto}}$ denotes "rank 1 candidate on the Pareto front, chosen by the respective MCDM method" and $M_1^{\text{MCDM\#1,reverse}}$ denotes "rank 1 candidate on the reverse front chosen by the respective MCDM method".

In the metric 1 case, the black dotted lines represent:

$$r_{\max} = \frac{M_1^{\min} - M_1^{\max}}{M_1^{\max}} \times 100 \tag{24}$$

i.e. the relative change between the maximum and minimum instances of the metric score encountered during sampling.

#### Additional Comments on Results Figures

- Selection of synthetic de Bruijn sequences (or random sequences, see below) may be carried out to ensure that no bad staple-staple (Metric 3) or intra-staple (Metric 4) interactions exist and/or to eradicate most off-target binding sites from very small origamis like O1 and O2.

- From an off-target binding site perspective, random synthetic scaffold sequences could be an effective substitute for de Bruijn sequences (and they require minimal computational power to produce).

- For large origamis with a fixed scaffold sequence, running a scaffold rotation selection using just metrics 3 and 4 could be beneficial to reduce unwanted staple-staple and intra-staple interactions.

- In general, metrics 3 and 4 have a different behaviour as compared to metrics 1 and 2 because they measure the strength of the worst binding site, rather than the total strength of all binding sites. Metric 4 can be optimised to nearly zero because: (i) typically there are only relatively few staples per nanostructure (≈200 in the worst case) and (ii) staples are typically short and strong intra-folds are relatively rare outlier cases which can be easily destroyed with change of scaffold sequence.

- Metric 3 (staple-staple co-folds) optimises to a lesser extent than metric 4 (and depends more on origami size) because the number of staple co-folds grows as the square of the staple number (Supplementary Figure 9). Thus, there are many more possibilities for a bad staple co-fold to exist as compared to a bad staple intra-fold. Additionally a staple co-fold has twice the number of bases able to make an off-target binding, as compared to a single staple intra-fold.

- As origami size increases, not only do the total number of off-target staple-scaffold bindings (metric 1) and intra-scaffold bindings (metric 2) increase, but the *local density* of binding sites also increases linearly (Supplementary Figures 7a and 8a). For metric 1, this is expected: if an origami has $s$ staples per base of scaffold (typically $s \approx 0.025$) and each staple has $b$ (assumed equal energy) off-target binding sites per base of scaffold on average, then there are $S = sL$ staples when the scaffold has length $L$ bases and $S(bL) = (sL)(bL) = sb(L^2)$ off-target binding sites in total. Hence the total number of off-target sites scales as the square of the scaffold length and the density $sb(L^2)/L = sbL$ scales linearly with scaffold length. A similar argument can be made for the linear density increase of intra-scaffold binding sites observed in metric 2: if the scaffold strand is divided into equal length segments with 1 segment per $u$ scaffold bases, and if each segment has $b$ binding sites (to other portions of the scaffold) per base of scaffold on average, then there are $U = L/u$ total segments on the scaffold and $U(bL) = (L/u)\,(bL) = (b/u)(L^2)$ intra-scaffold binding sites in total. Intra-scaffold binding site density scales linearly as $(b/u)L$.

- Scaffold sequence selection becomes increasing ineffective as origami size increases (Supplementary Figures 7b and 8b).

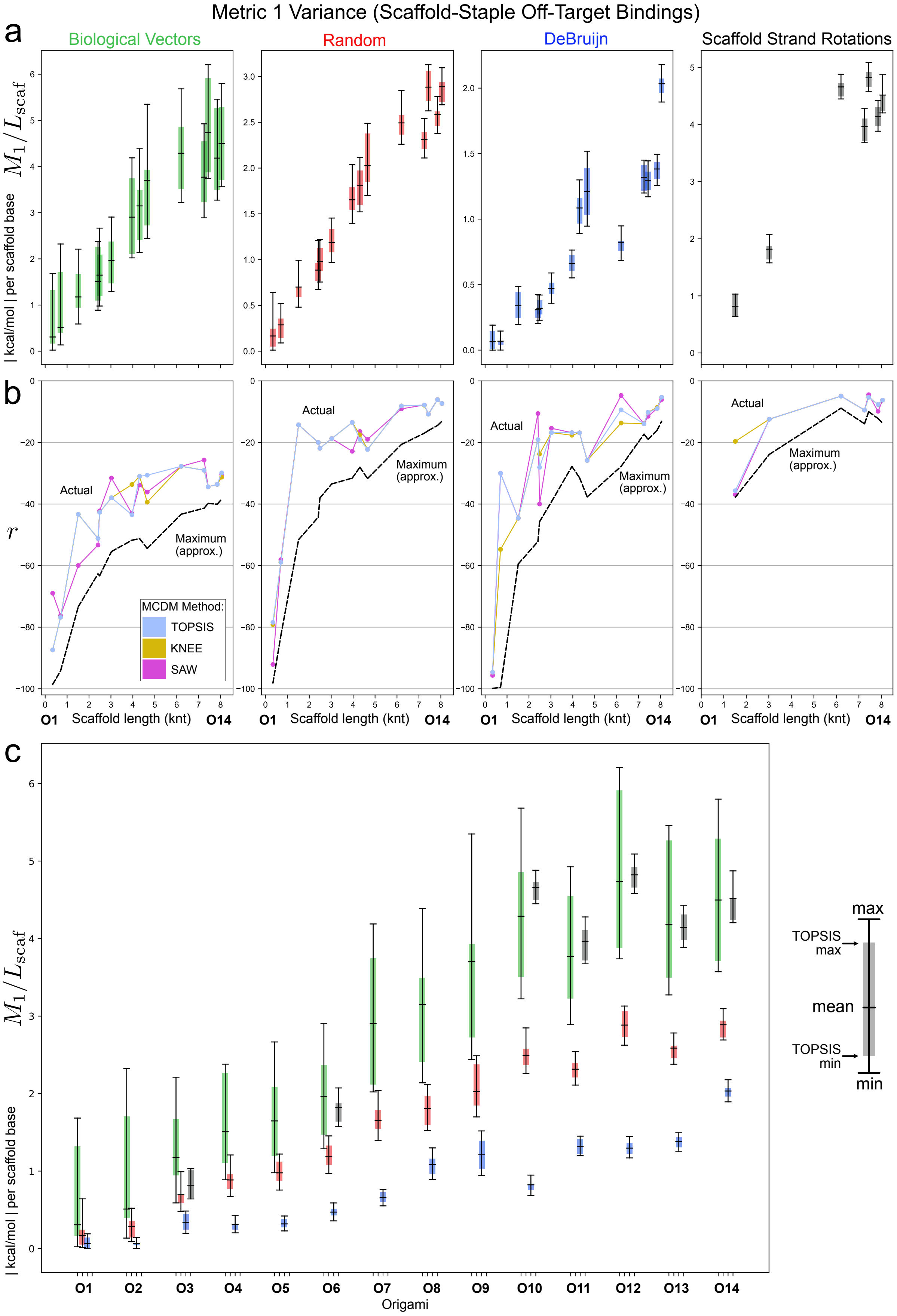


Supplementary Figure 7: Metric $M_1$ variance over 14 origami test set with scaffold sequences selected from four different sequence pool types. See text for description.

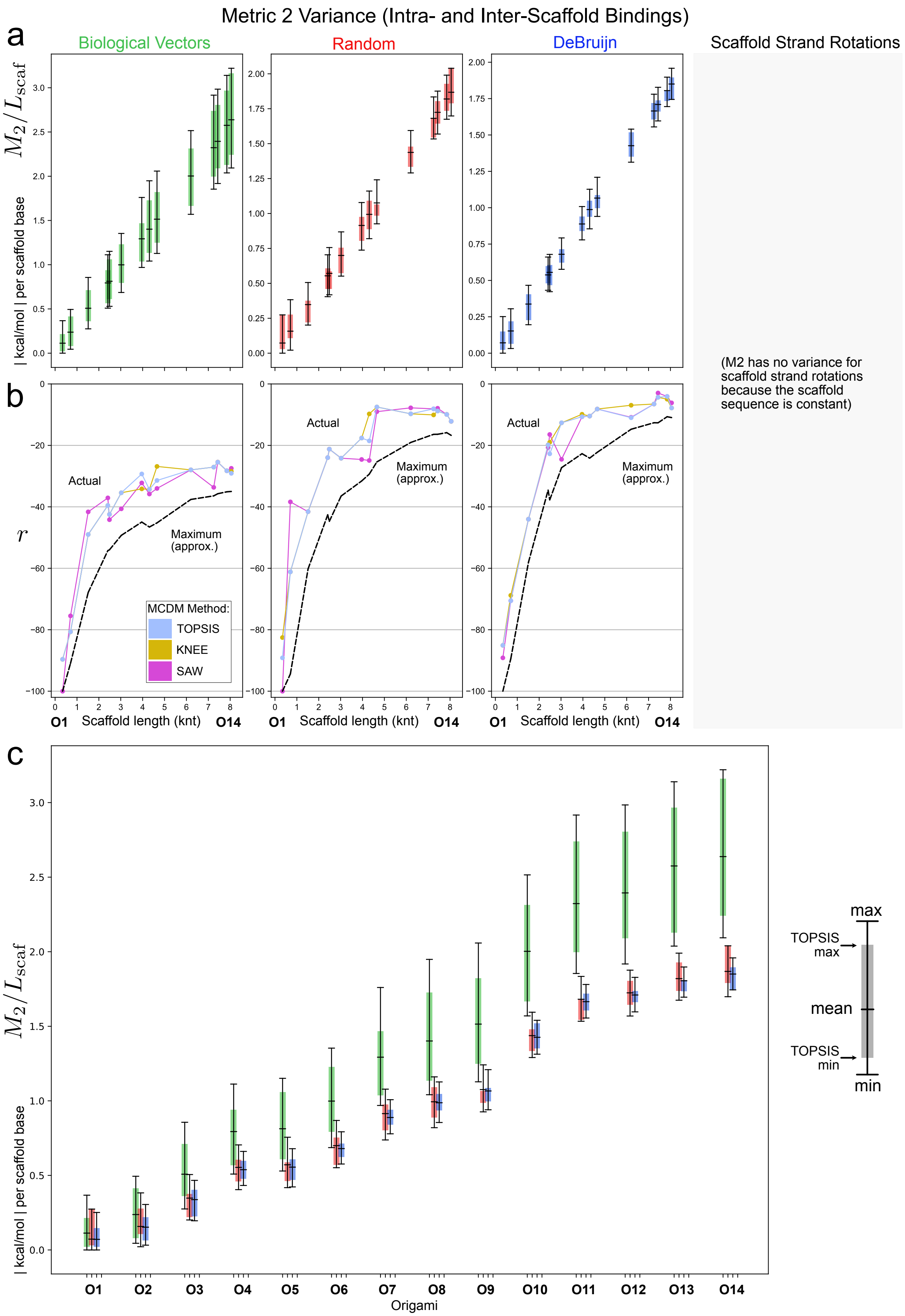


Supplementary Figure 8: Metric $M_2$ variance over 14 origami test set with scaffold sequences selected from four different sequence pool types. See text for description.

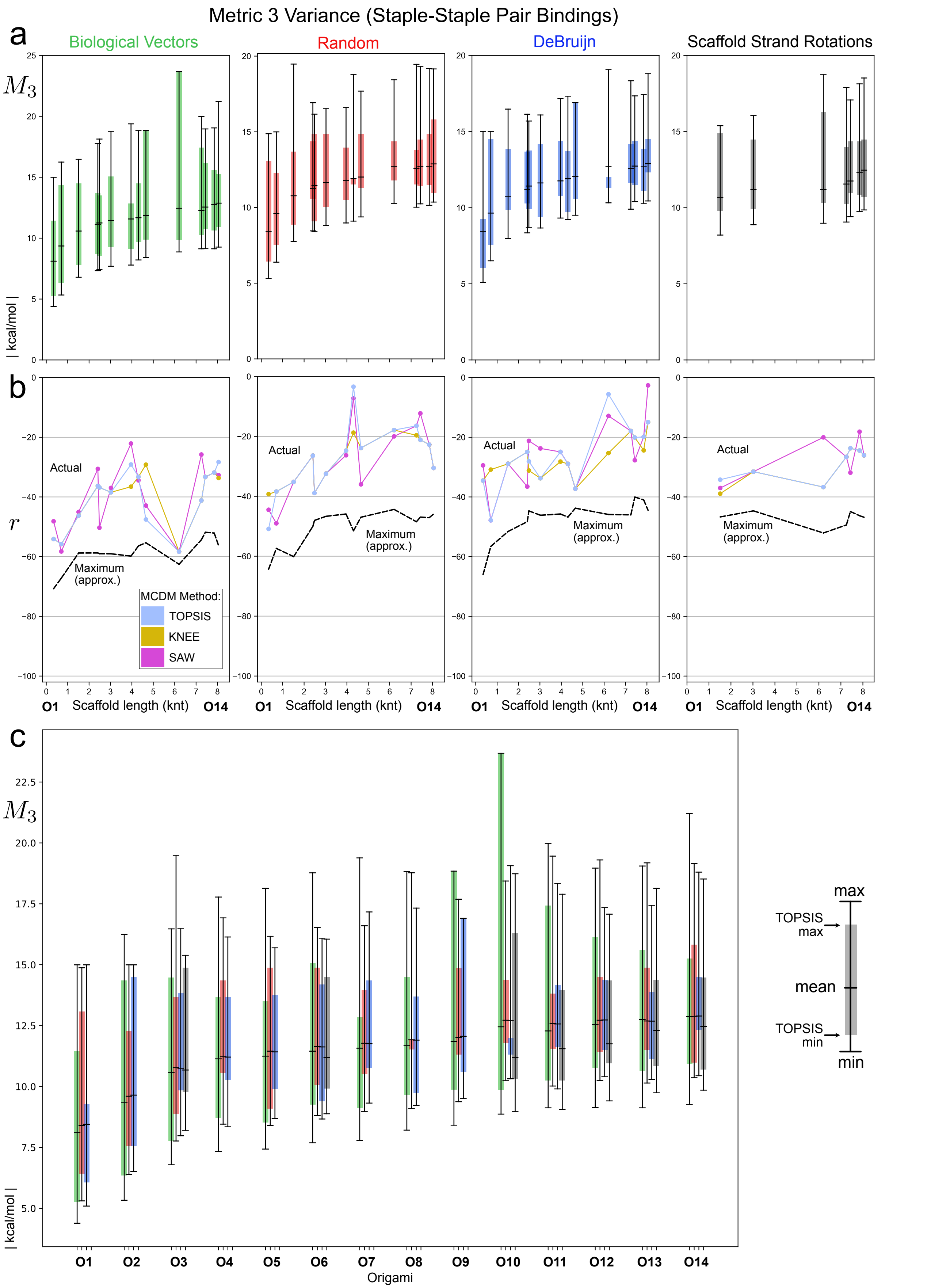


Supplementary Figure 9: Metric $M_3$ variance over 14 origami test set with scaffold sequences selected from four different sequence pool types. See text for description.

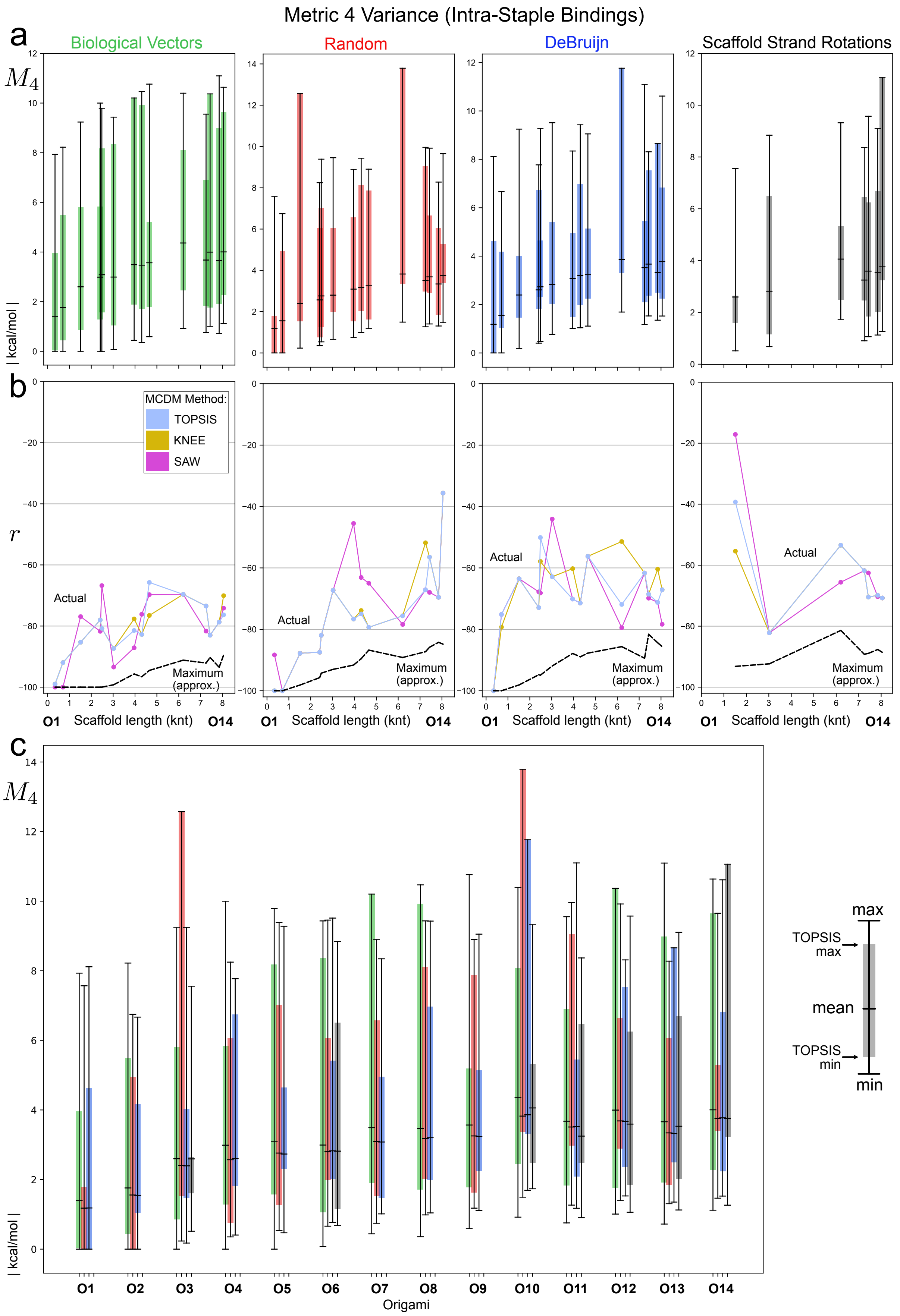


Supplementary Figure 10: Metric $M_4$ variance over 14 origami test set with scaffold sequences selected from four different sequence pool types. See text for description.

### Supplementary Note 6 Metric Pair Correlations

We did not observe Pearson correlation between pairs of metric scores when scaffold sequences in the random, De Bruijn and scaffold strand rotation sequence pools were scored (Supplementary Figure 11).

Pearson correlation across all metric score pairs was observed for scaffold sequences from the biological vector pool (up to $r$ = 0.5), particularly between $M_1$ and $M_2$ for larger origamis ($r \approx$ 0.8). However, rather than being an intrinsic property of the metrics, this correlation is likely a small size effect of the biological vector pool. The biological vector pool was constructed from five commercially available vectors and scaffold sequences "cut-out" from these vectors would have experienced some degree of sequence overlap – particularly for longer scaffold origamis – generating similar yet different metric scores. Therefore, the metric scores are not intrinsically pair-wise correlated.

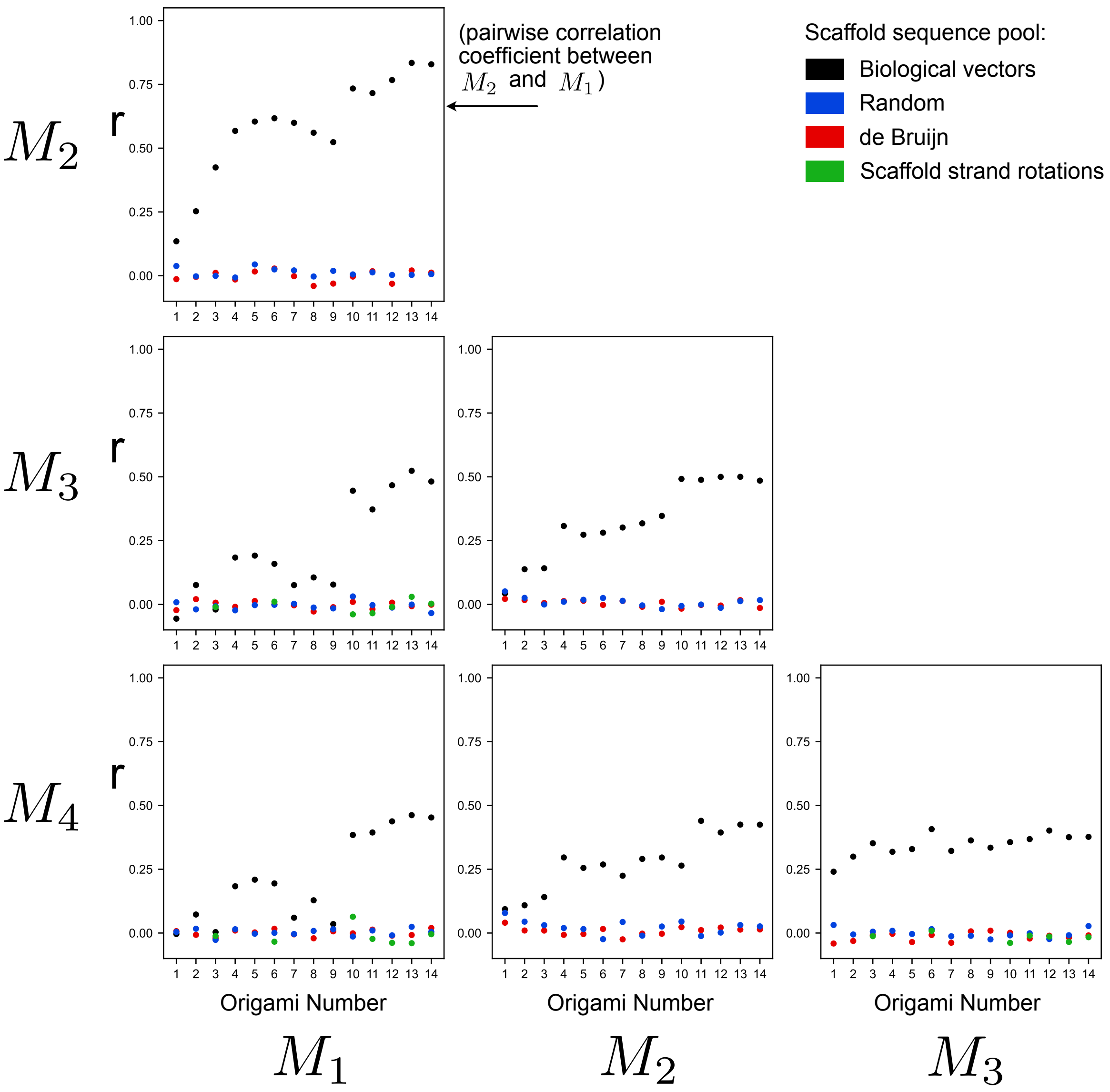


Supplementary Figure 11: The Pearson correlation coefficient $r$ between each pair of metric scores. Calculated for each origami in the test set (Supplementary Note 4) and for each sequence pool type. For the biological vector, random and de Bruijn sequence pools, each correlation value is calculated from $n$=5000 pairs of metric scores. For the scaffold strand rotation pool, the number of samples depends on the scaffold length.

### Supplementary Note 7 Metric Score Correlation with Repeats and GC Content

Supplementary Figure 12 investigates if total kmer repetitions and GC content of a scaffold sequence are correlated with $M_1$ to $M_4$ scores, and thus to what extent the former can be quick predictors of the latter. Analysis is run with the DNA origami triangle in the main paper with a 5000 sequences taken from biological vectors (VEC pool). Similar results were obtained with the DNA origami rectangle. Total kmer repeats is calculated by computing the number of repeated subsequences of length 1, 2, 3, 4.... and taking the sum. We observed that:

- **Total kmer repeats.** Increasing total kmer repeats is strongly correlated with increasing $M_1$ score and correlated (but more weakly) with increasing $M_2$ score, although there are outlier groups. Total kmer repeats appear uncorrelated with $M_3$ or $M_4$ scores and with overall TOPSIS score of the scaffold sequence.

- **GC content.** Low GC content or excessively high GC content are associated with many kmer repeats, and thus $M_1$ and $M_2$ scores are minimised at intermediate values of GC content only ($0.47$ fraction). Metrics $M_3$ and $M_4$ are correlated with GC content and minimise as GC content decreases (because the latter metrics are sequence dependent and low GC content leads to weaker bindings). Overall TOPSIS score is not correlated with GC content.

- **GC content is the best proxy** for predicting lower $M_1$ to $M_4$ scores. A value close to $0.47$ is likely to yield lower $M_1$ and $M_2$ scores. However, at each GC content value there is considerable variation of all metric scores, and hence GC content alone cannot substitute sequence selection (which is specific to an origami design).

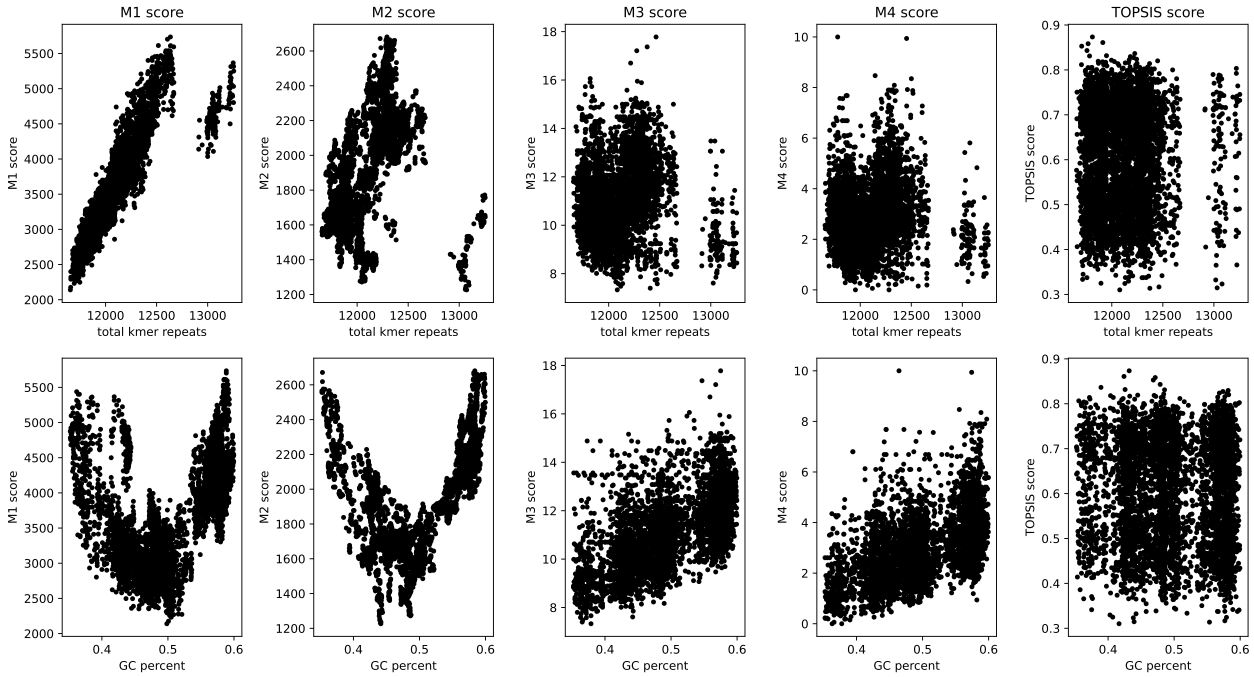


Supplementary Figure 12: Total kmer repetitions and GC content correlation with $M_1$ to $M_4$ scores for DNA origami triangle in the main paper. See text.

### Part III

### DNA Origamis Selected for *In Vitro* Assembly

In this part, we detail the scadnano schematics for the triangle and rectangle DNA origami variants used in the paper. We also detail the off-target binding site energy distributions for the different variants. The schematics below were drawn with scadnano.org [12]. See Zenodo repository https://doi.org/10.5281/zenodo.14191020 for electronic versions and sequences.

#### Supplementary Note 8 Triangle Origami Variants

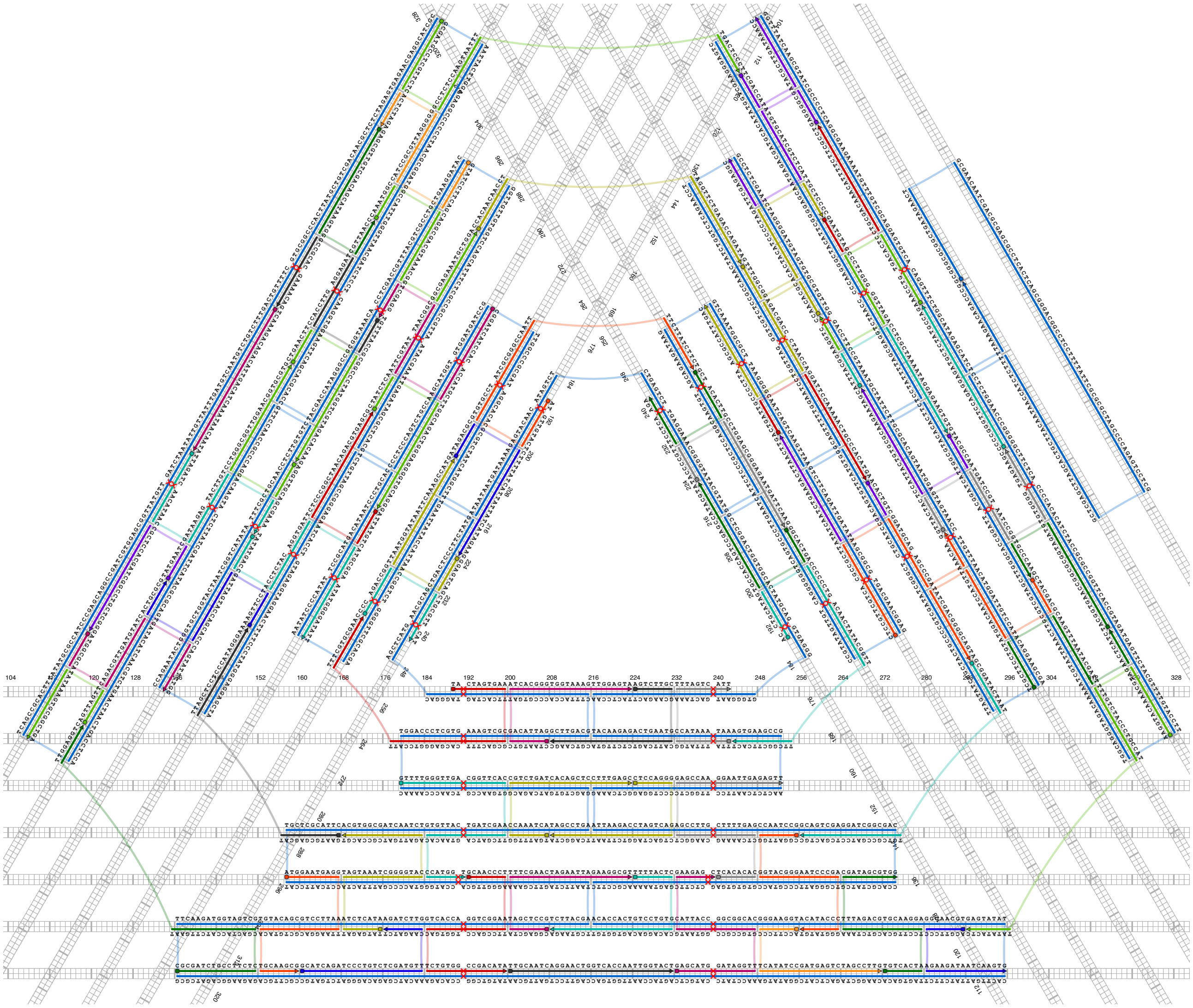

Supplementary Figure 13: Triangle $T_1$ (blinded name: DEER). Scaffold sequence (5'-3', 2410 nt) synthesized from gBlocks Gene Fragment DNA template. Zoom for sequence detail.

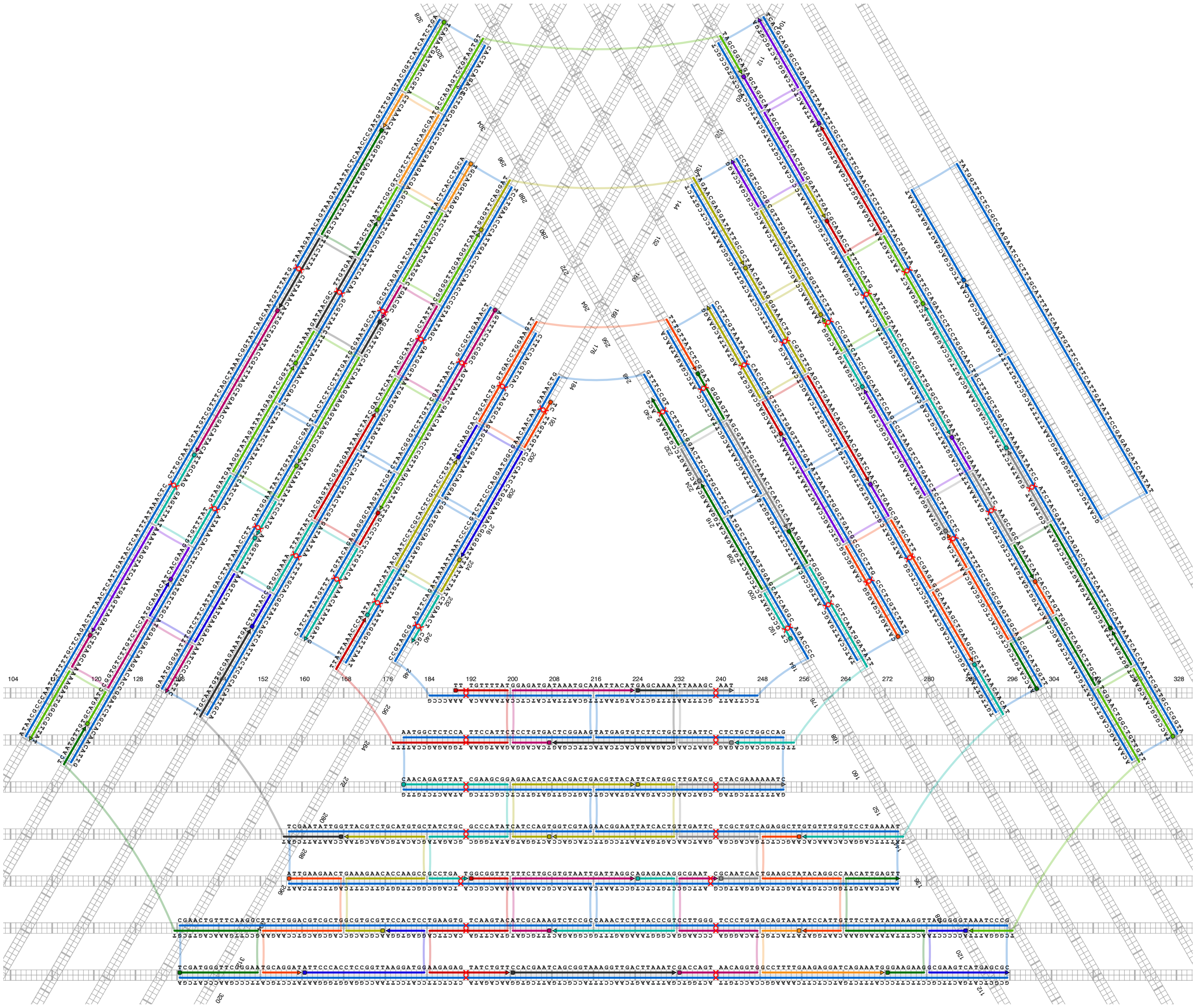

Supplementary Figure 14: Triangle $T_2$ (blinded name: LION). Scaffold sequence (5'-3', 2410 nt) synthesized from Lambda dsDNA template. Zoom for sequence detail.

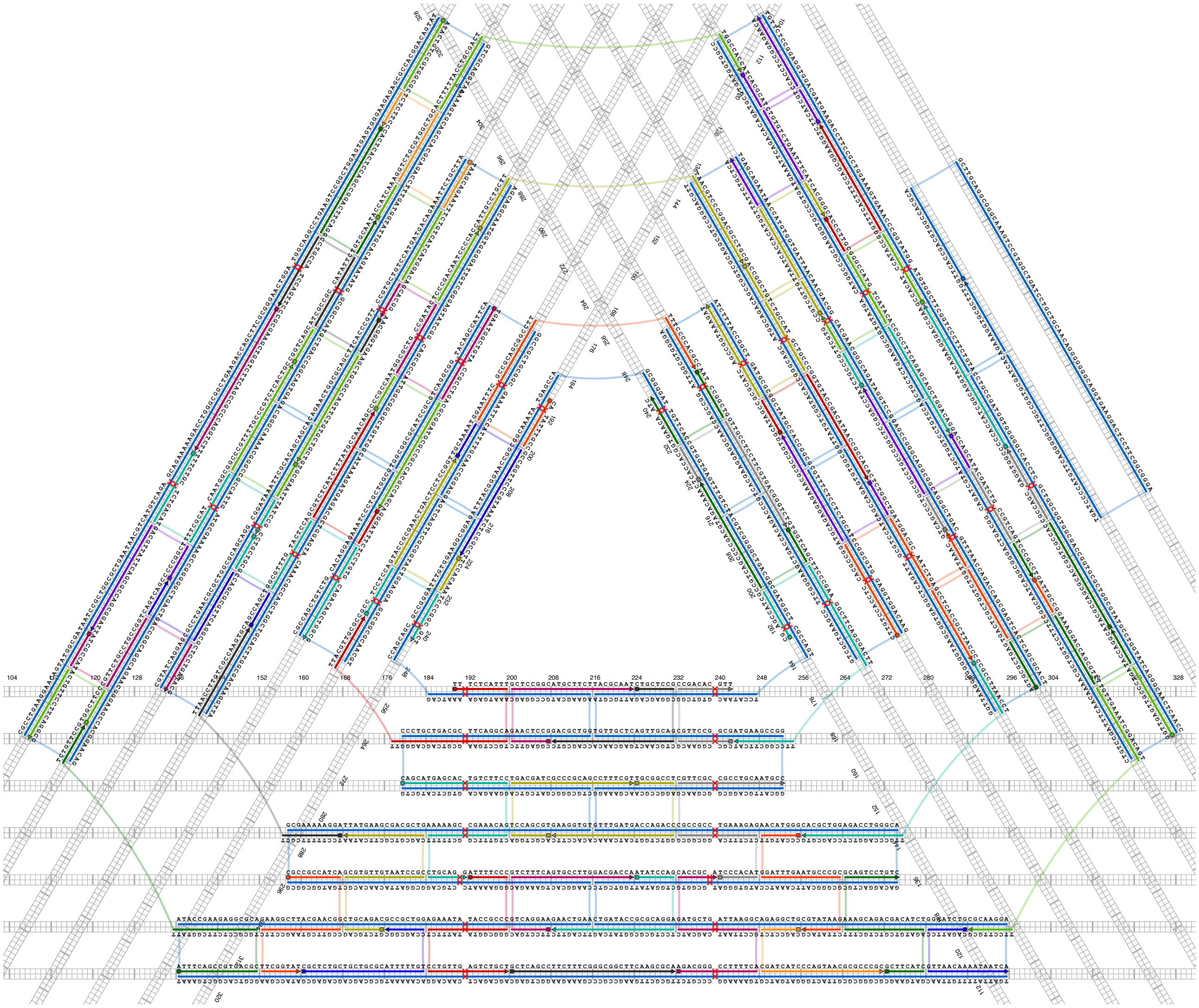

Supplementary Figure 15: Triangle $T_3$ (blinded name: BEAR). Scaffold sequence (5'-3', 2410 nt) synthesized from Lambda dsDNA template. Zoom for sequence detail.

### Supplementary Note 9 Rectangle Origami Variants

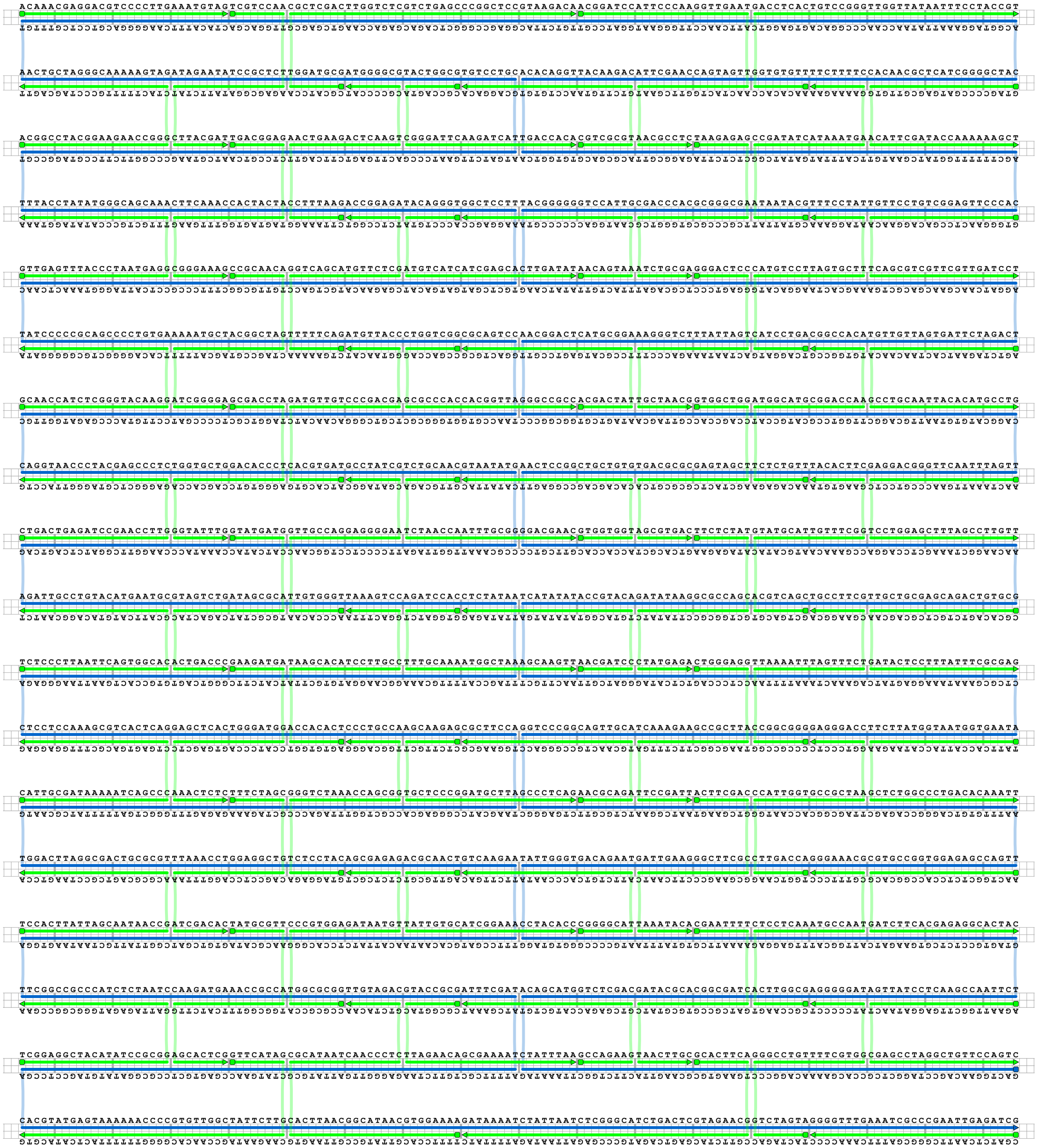

Supplementary Figure 16: Rectangle $R_1$ (blinded name: GOAT). Scaffold sequence (5'-3', 2484 nt) synthesized from gBlocks Gene Fragment DNA template. Zoom for detail.

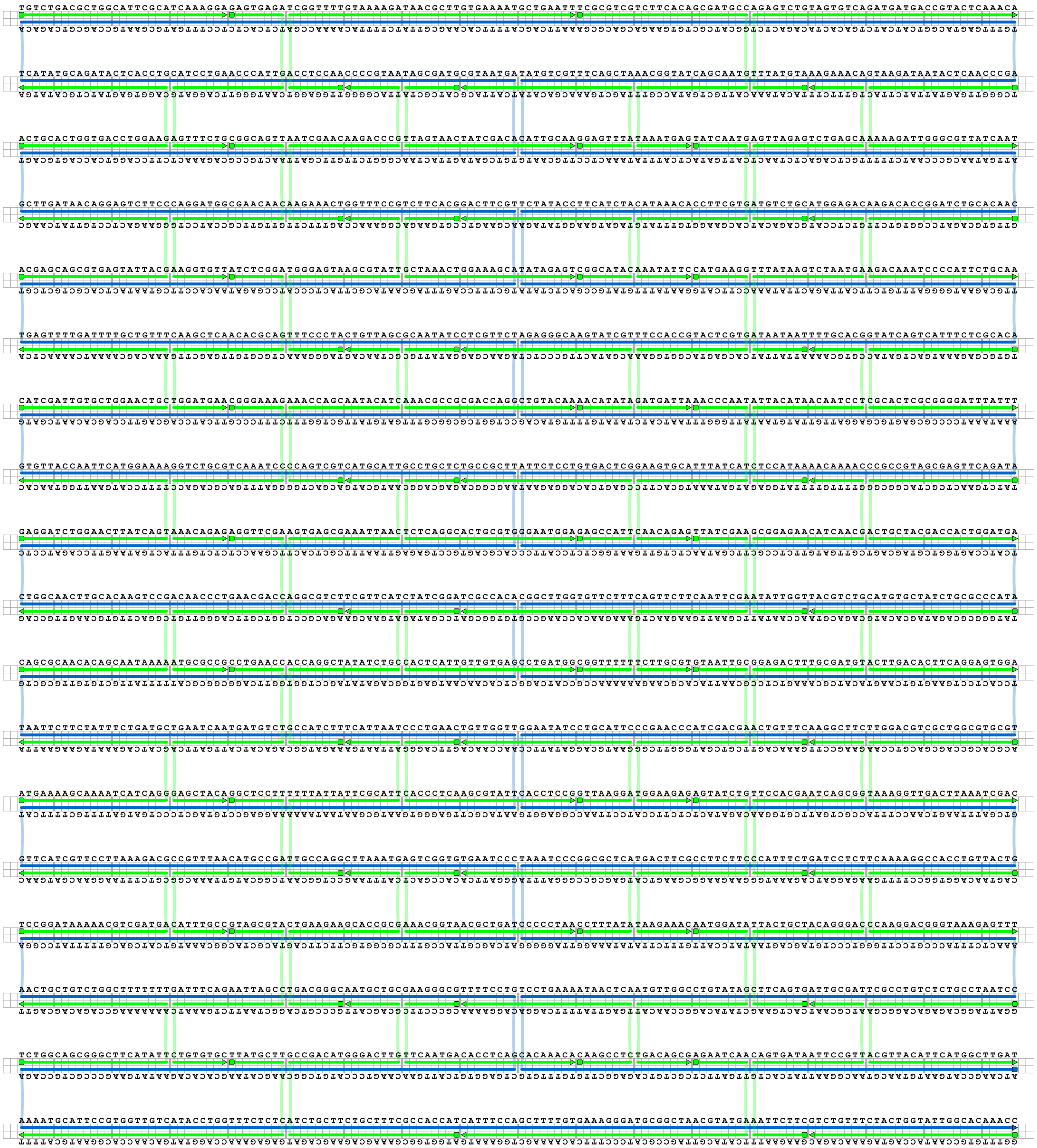

Supplementary Figure 17: Rectangle $R_2$ (blinded name: LAMB). Scaffold sequence (5'-3', 2484 nt) synthesized from Lambda dsDNA template. Zoom for detail.

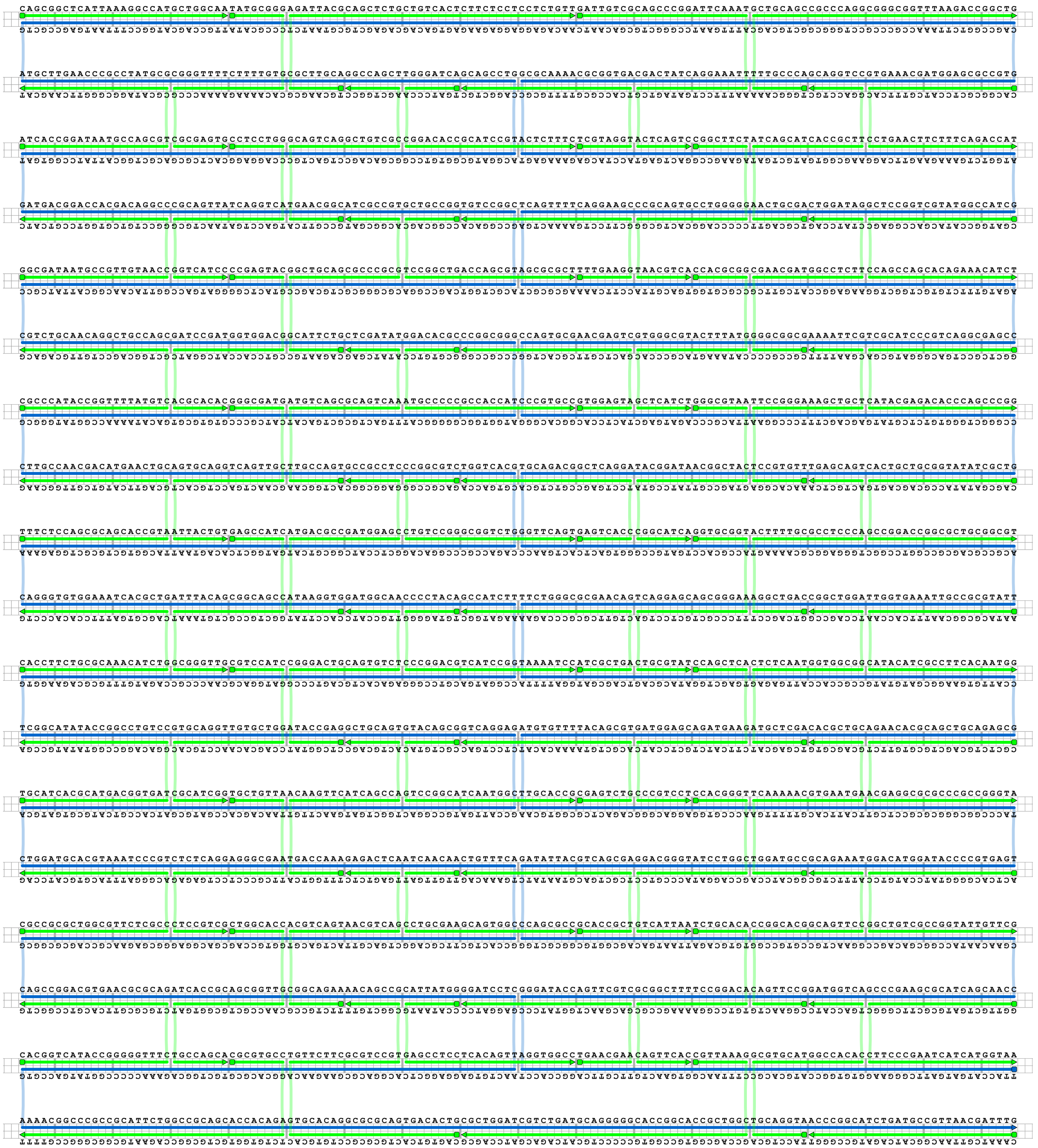

Supplementary Figure 18: Rectangle $R_3$ (blinded name: MOLE). Scaffold sequence (5'-3', 2484 nt) synthesized from Lambda dsDNA template. Zoom for detail.

### Supplementary Note 10 Off-target Binding Site Energy Distributions

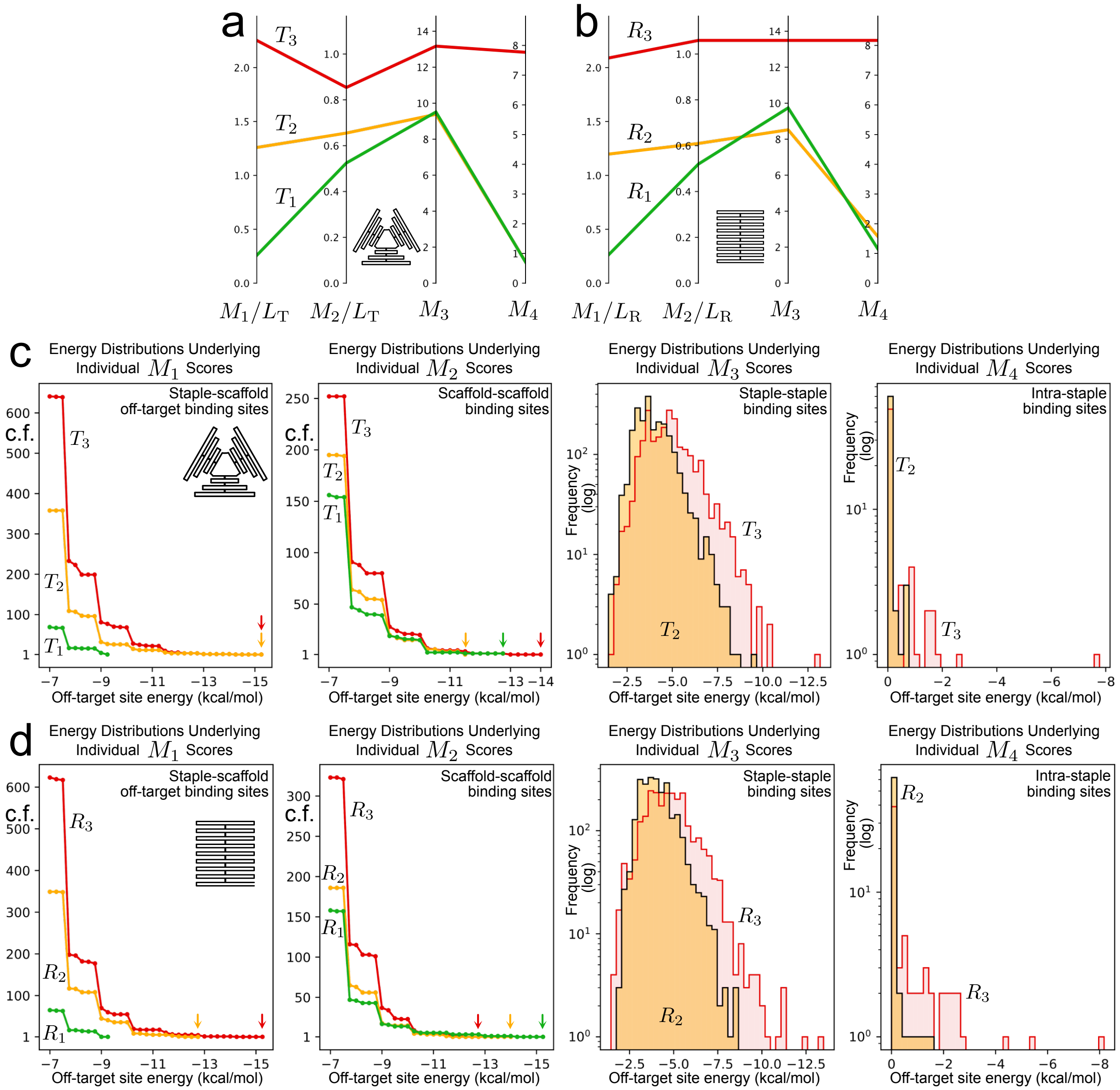


Supplementary Figure 19: **(a)** Metric scores for the three triangle origami variants, compared on a parallel co-ordinate plot. Scores for metrics 1 and 2 are normalised by the length of the triangle scaffold $L_T = 2410$. **(b)** Similar comparison of metric scores for the three rectangle origami variants with $L_R = 2484$. **(c)** Off-target binding site energy distributions underlying the metric scores for each triangle origami variant. Distributions underlying $M_1$ and $M_2$ scores are shown as cumulative frequency (c.f) plots representing number of off-target binding sites with a more negative $\Delta G^0(37C)$ than the value on the x-axis. Coloured vertical arrows indicate where cumulative frequency falls to zero for each variant. Distributions underlying $M_3$ and $M_4$ scores are frequency histograms and compare only $T_2$ and $T_3$ as variant $T_1$ has similar energy distributions to $T_2$. **(d)** Off-target binding site energy distributions underlying each metric score for each rectangle origami variant.

#### Part IV

### Additional Experimental Results

This part contains experimental results relating to the amplification, purification and gel electrophoresis pre-screening of the triangle and rectangle DNA origami variants described in the main paper. Experiments on the triangle and rectangle variants were performed blinded, thus the results below refer to the blinded variant names in Supplementary Table 3.

| **TRIANGLE DNA ORIGAMI** | |
|---|---|
| $T_1$ | DEER |
| $T_2$ | LION |
| $T_3$ | BEAR |
| **RECTANGLE DNA ORIGAMI** | |
| $R_1$ | GOAT |
| $R_2$ | LAMB |
| $R_3$ | MOLE |

Supplementary Table 3: Blinded names for origami variants.

### Supplementary Note 11 PCR Details

| NAME | SEQUENCE (5'-3') |
|---|---|
| **TRIANGLE DNA ORIGAMI** | |
| SC_BEAR_Forward | **G*****T*****T*****C*****A***CCGACGTGGCCACGC |
| SC_BEAR_Reverse | CAAAGCGCACCTTTCTTACAGAAGGGAC |
| SC_LION_Forward | **T*** **T*** **C*** **G*** **C***GACGAGTAGATGCAATTATG |
| SC_LION_Reverse | TTGTCGGACTTGTGCAAGTTG |
| SC_DEER_Forward | **G*** **C*** **G*** **G*** **G***CGCGGCTGATAGAAC |
| SC_DEER_Reverse | CGGGTCTGTTCTTTAAAGATGCTCTATGCC |
| **RECTANGLE DNA ORIGAMI** | |
| SC_LAMB_Forward | **A*** **T*** **C*** **A*** **A***GCCATGAATGTAACGTAAC |
| SC_LAMB_Reverse | GGTTTGTGCCAATACCAGTAG |
| SC_MOLE_Forward | **T*** **T*** **A*** **C*** **C***ATGATGATTCGGGAAGG |
| SC_MOLE_Reverse | CAAATCGTTAACGGCATCAGATG |
| SC_GOAT_Forward | **G*** **A*** **C*** **T*** **G***GAACAGCCTAGGCTC |
| SC_GOAT_Reverse | CGATCTCAATTCGGGCGATTTCAAAC |

Supplementary Table 4: List of primer sequences used to amplify double-stranded DNA template (gBlocks Gene Fragments and Lambda DNA). Phosphorothioated DNA bases at the 5' end (forward primers) are in bold with asterisk.

| DNA TEMPLATE | QUANTITY (ng in 50 μL) | STEP | TEMPERATURE (°C) | TIME (s) | CYCLES |
|---|---|---|---|---|---|
| LAMBDA (BEAR) | 1 | Initial denaturation | 98 | 30 | 1 |
| | | Denaturation | 98 | 10 | 15 |
| | | Annealing/Extension | 72 | 50 | |
| | | Final Extension | 72 | 120 | 1 |
| LAMBDA (LION) | 1 | Initial denaturation | 98 | 30 | 1 |
| | | Denaturation | 98 | 10 | 15 |
| | | Annealing | 67 | 20 | |
| | | Extension | 72 | 25 | |
| | | Final Extension | 72 | 120 | 1 |
| gBlocks (DEER) | 2.5 | Initial denaturation | 98 | 30 | 1 |
| | | Denaturation | 98 | 10 | 15 |
| | | Annealing/Extension | 72 | 50 | |
| | | Final Extension | 72 | 120 | 1 |
| LAMBDA (LAMB) | 1 | Initial denaturation | 98 | 30 | 1 |
| | | Denaturation | 98 | 10 | 22 |
| | | Annealing | 64 | 20 | |
| | | Extension | 72 | 25 | |
| | | Final Extension | 72 | 120 | 1 |
| LAMBDA (MOLE) | 10 | Initial denaturation | 98 | 30 | 1 |
| | | Denaturation | 98 | 10 | 15 |
| | | Annealing | 65 | 20 | |
| | | Extension | 72 | 25 | |
| | | Final Extension | 72 | 120 | 1 |
| gBlocks (GOAT) | 5 | Initial denaturation | 98 | 30 | 1 |
| | | Denaturation | 98 | 10 | 12 |
| | | Annealing | 71 | 20 | |
| | | Extension | 72 | 25 | |
| | | Final Extension | 72 | 120 | 1 |

Supplementary Table 5: Template amount and thermocycling conditions for PCR amplification of double-stranded DNA scaffold.

#### Supplementary Note 12 Single Stranded DNA Scaffold Purification

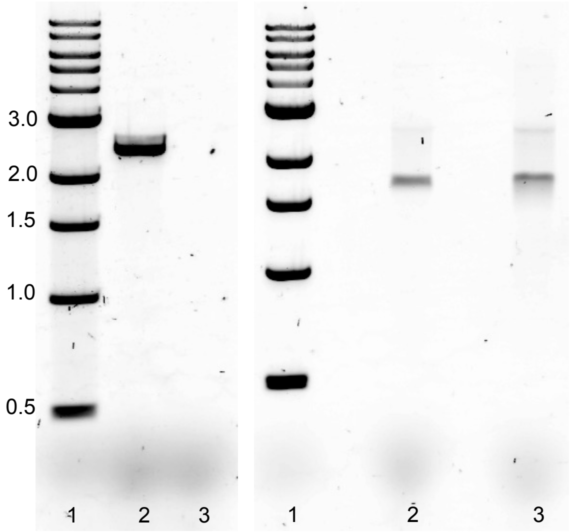


Supplementary Figure 20: Purified single-stranded DNA scaffold sequence (triangle origami, Bear variant). 1% TBE agarose gel electrophoresis image after SYBR® Gold staining. Left image (amplified dsDNA), lanes 1: 1 kb DNA ladder; 2: pcr product; 3: pcr blank. Right image (purified ssDNA after overnight T7 exonuclease digestion), lanes 1: 1 kb DNA ladder; 2: non-purified ssDNA scaffold; 3: purified ssDNA scaffold. Molecular sizes in kilobases are indicated.

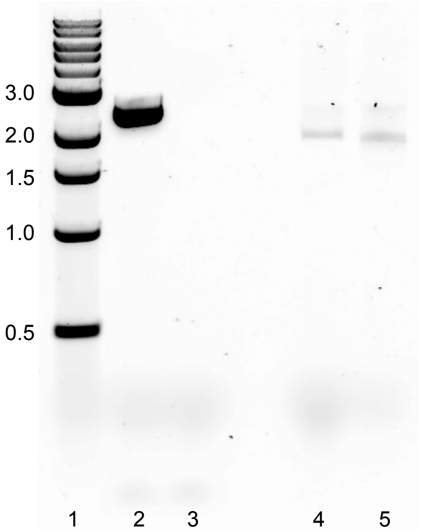


Supplementary Figure 21: Purified single-stranded DNA scaffold sequence (triangle origami, Lion variant). 1% TBE agarose gel electrophoresis image after SYBR® Gold staining. Lanes. 1: 1 kb DNA ladder; 2: pcr product; 3: pcr blank; 4: non-purified ssDNA scaffold after overnight T7 exonuclease digestion; 5: purified ssDNA scaffold. Molecular sizes in kilobases are indicated.

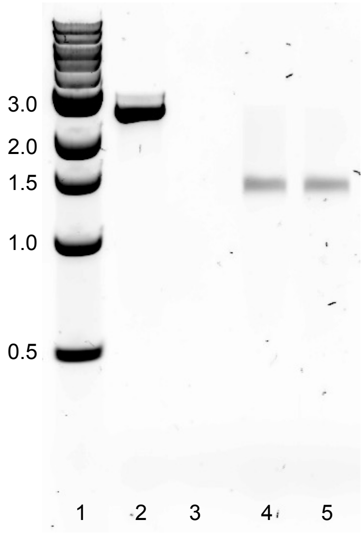


Supplementary Figure 22: Purified single-stranded DNA scaffold sequence (triangle origami, Deer variant). 1% TBE agarose gel electrophoresis image after SYBR® Gold staining. Lanes. 1: 1 kb DNA ladder; 2: pcr product; 3: pcr blank; 4: non-purified ssDNA scaffold after overnight T7 exonuclease digestion; 5: purified ssDNA scaffold. Molecular sizes in kilobases are indicated.

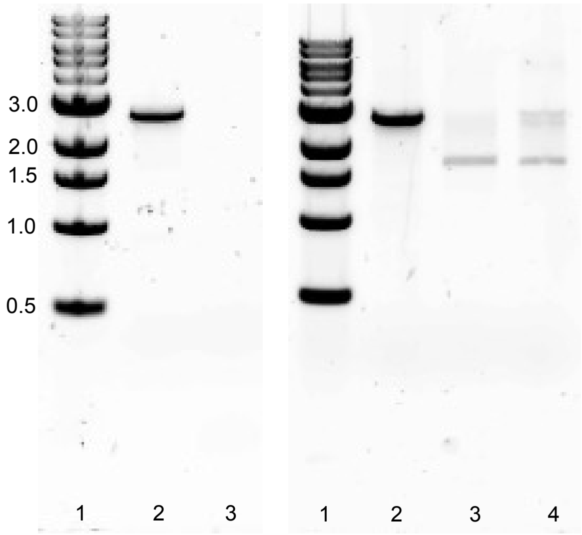


Supplementary Figure 23: Purified single-stranded DNA scaffold sequence (rectangle origami, Lamb variant). 1% TBE agarose gel electrophoresis image after SYBR® Gold staining. Left image (amplified dsDNA), lanes 1: 1 kb DNA ladder; 2: pcr product; 3: pcr blank. Right image (purified ssDNA after overnight T7 exonuclease digestion), lanes 1: 1 kb DNA ladder; 2: purified pcr product; 3: non-purified ssDNA scaffold; 4: purified ssDNA scaffold. Molecular sizes in kilobases are indicated.

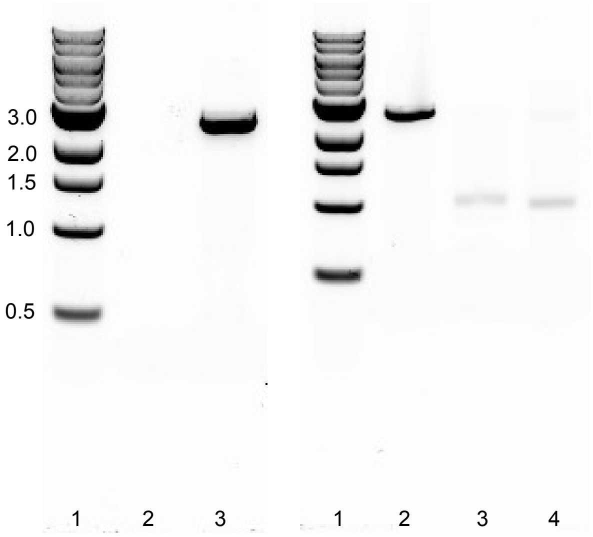


Supplementary Figure 24: Purified single-stranded DNA scaffold sequence (rectangle origami, Mole variant). 1% TBE agarose gel electrophoresis image after SYBR® Gold staining. Left image (amplified dsDNA), lanes 1: 1 kb DNA ladder; 2: pcr blank; 3: pcr product. Right image (purified ssDNA after overnight T7 exonuclease digestion), lanes 1: 1 kb DNA ladder; 2: purified pcr product; 3: non-purified ssDNA scaffold; 4: purified ssDNA scaffold. Molecular sizes in kilobases are indicated.

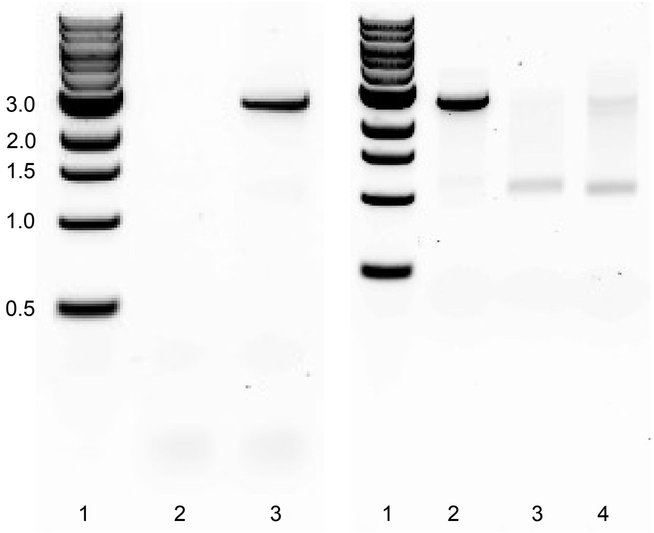


Supplementary Figure 25: Purified single-stranded DNA scaffold sequence (rectangle origami, Goat variant). 1% TBE agarose gel electrophoresis image after SYBR® Gold staining. Left image (amplified dsDNA), lanes 1: 1 kb DNA ladder; 2: pcr blank; 3: pcr product. Right image (purified ssDNA after overnight T7 exonuclease digestion), lanes 1: 1 kb DNA ladder; 2: purified pcr product; 3: non-purified ssDNA scaffold; 4: purified ssDNA scaffold. Molecular sizes in kilobases are indicated.

#### Supplementary Note 13 Gel Electrophoresis Pre-Screening

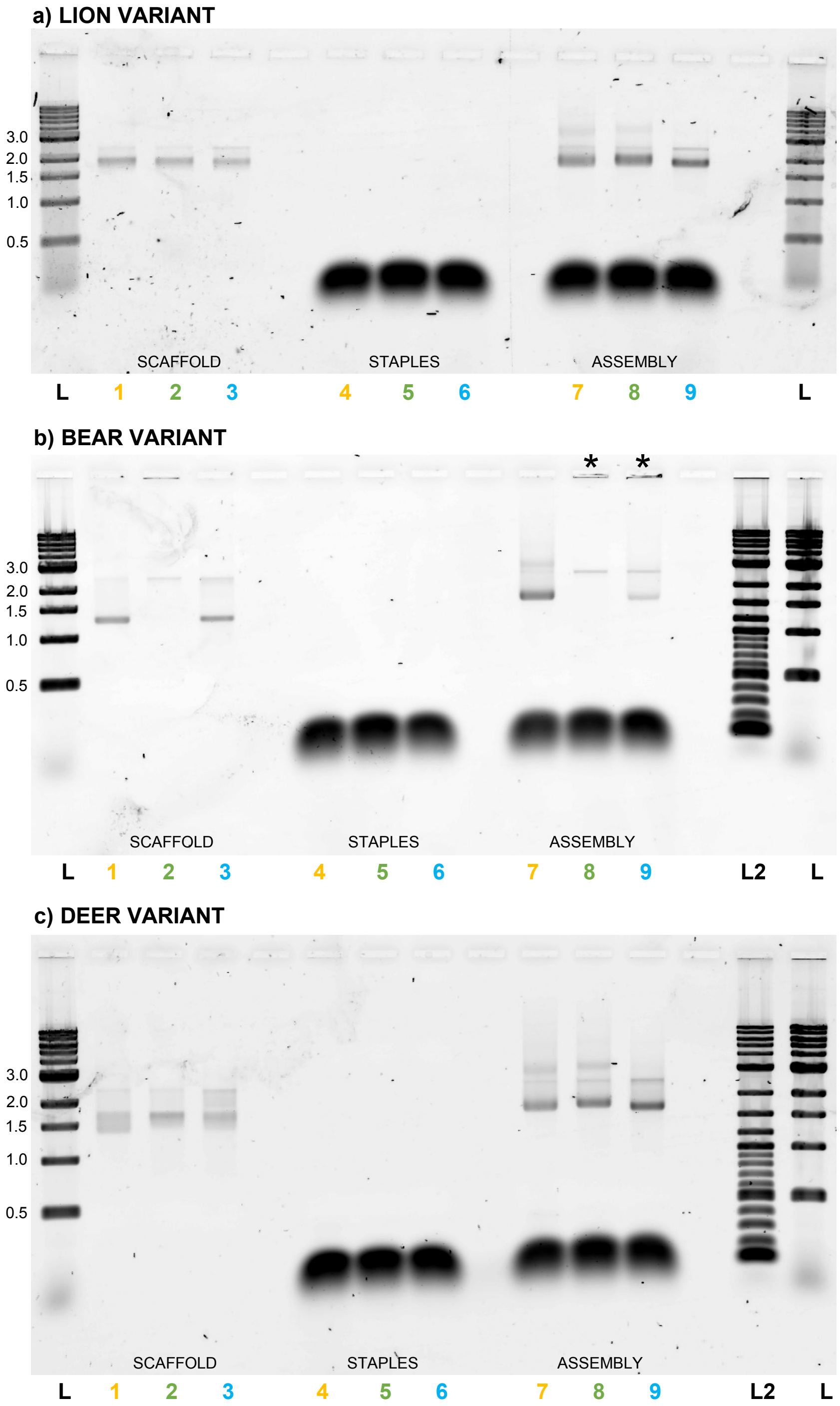


Supplementary Figure 26: Triangle DNA origami variants (Lion, Bear, Deer) and gel electrophoresis pre-screening. Laser-scanned images of SYBR® Gold-stained 1% TBE agarose gels loaded with scaffolds, staple strands mixes and assembly products obtained from isothermal annealing without initial denaturation (numbers in orange), isothermal annealing with initial denaturation (numbers in green) and fast annealing ramp (numbers in blue). a) Lion variant; b) Bear variant; c) Deer variant; L: 1 kb Ladder; L2: 1 kb Plus Ladder; *: aggregates in the loading well. Molecular sizes in kilobases are indicated.

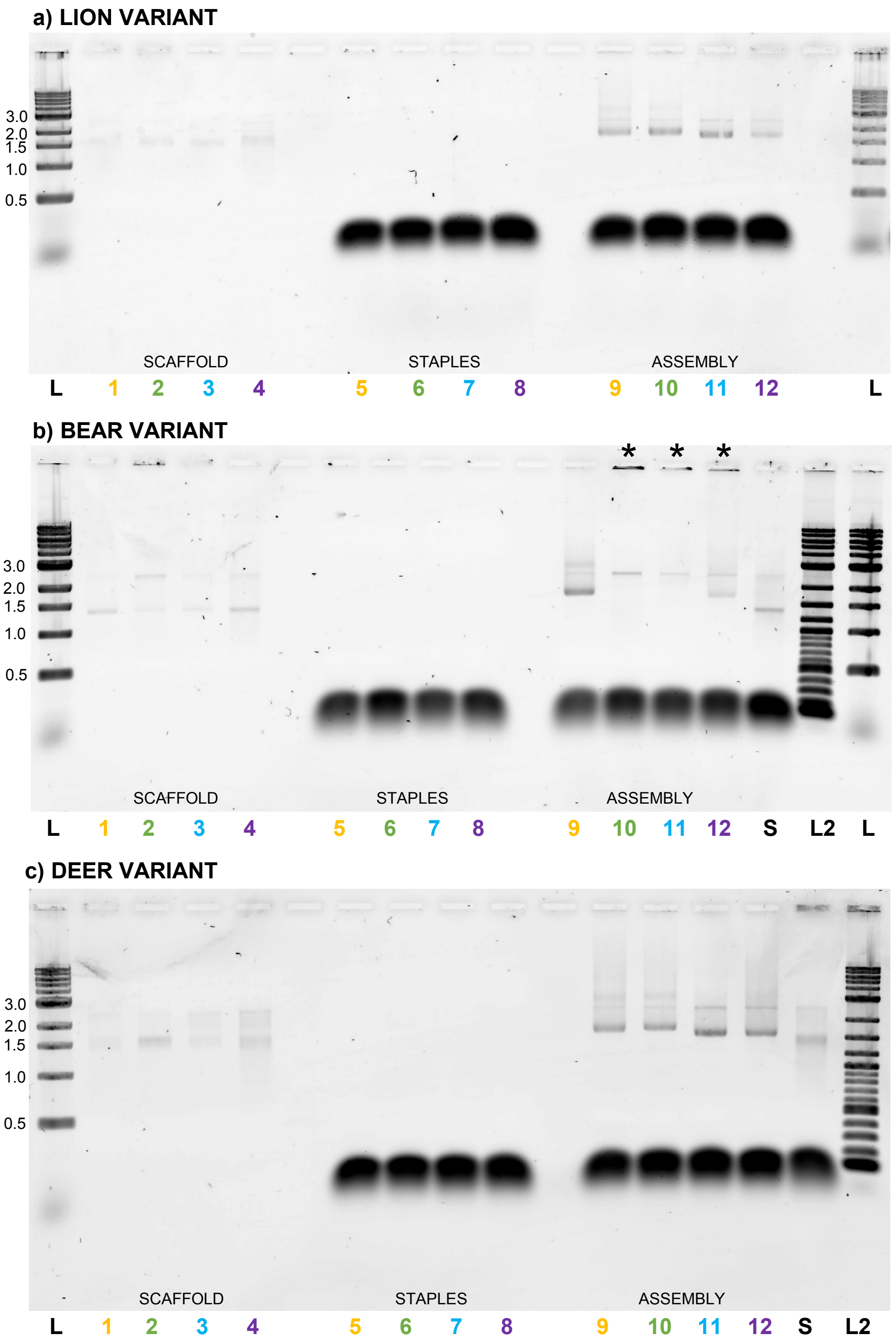


Supplementary Figure 27: Triangle DNA origami variants (Lion, Bear, Deer) and gel electrophoresis pre-screening. Laser-scanned images of SYBR® Gold-stained 1% TBE agarose gels loaded with scaffolds, staple strands mixes and assembly products obtained from isothermal annealing without initial denaturation (numbers in orange), isothermal annealing with initial denaturation (numbers in green), fast annealing ramp (numbers in blue) and slow annealing ramp (numbers in purple). Samples annealed considering the first 3 temperature ramps (as shown in S26) were kept at 4 °C and run again with samples annealed the day after with a slower ramp. a) Lion variant; b) Bear variant; c) Deer variant; L: 1 kb Ladder; L2: 1 kb Plus Ladder; S: Bear scaffold and non-complementary Lion strands set (negative control); *: aggregates in the loading well. Molecular sizes in kilobases are indicated.

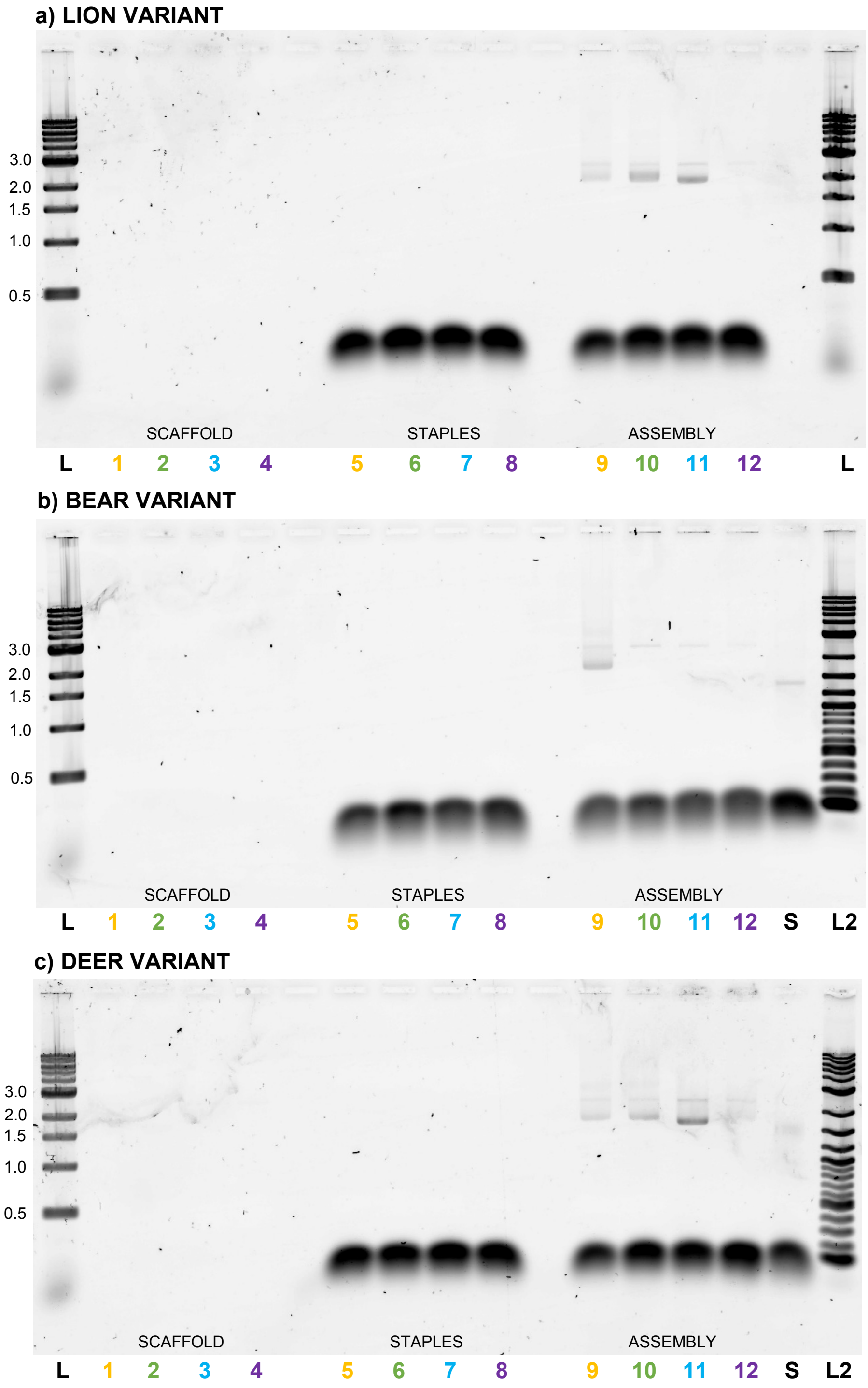


Supplementary Figure 28: Triangle DNA origami variants (Lion, Bear, Deer) and gel electrophoresis pre-screening. Laser-scanned images of SYBR® Gold-stained 1% TBE agarose gels loaded with scaffolds, staple strands mixes and assembly products obtained from isothermal annealing without initial denaturation (numbers in orange), isothermal annealing with initial denaturation (numbers in green), fast annealing ramp (numbers in blue) and slow annealing ramp (numbers in purple). Samples annealed considering 4 different temperature conditions (as shown in S26 and S27) were kept at 4 °C and run again after 4 (slow ramp) or 5 days (isothermal foldings and fast ramp). a) Lion variant; b) Bear variant; c) Deer variant; L: 1 kb Ladder; L2: 1 kb Plus Ladder; S: Bear scaffold and non-complementary Lion strands set (negative control). Molecular sizes in kilobases are indicated.

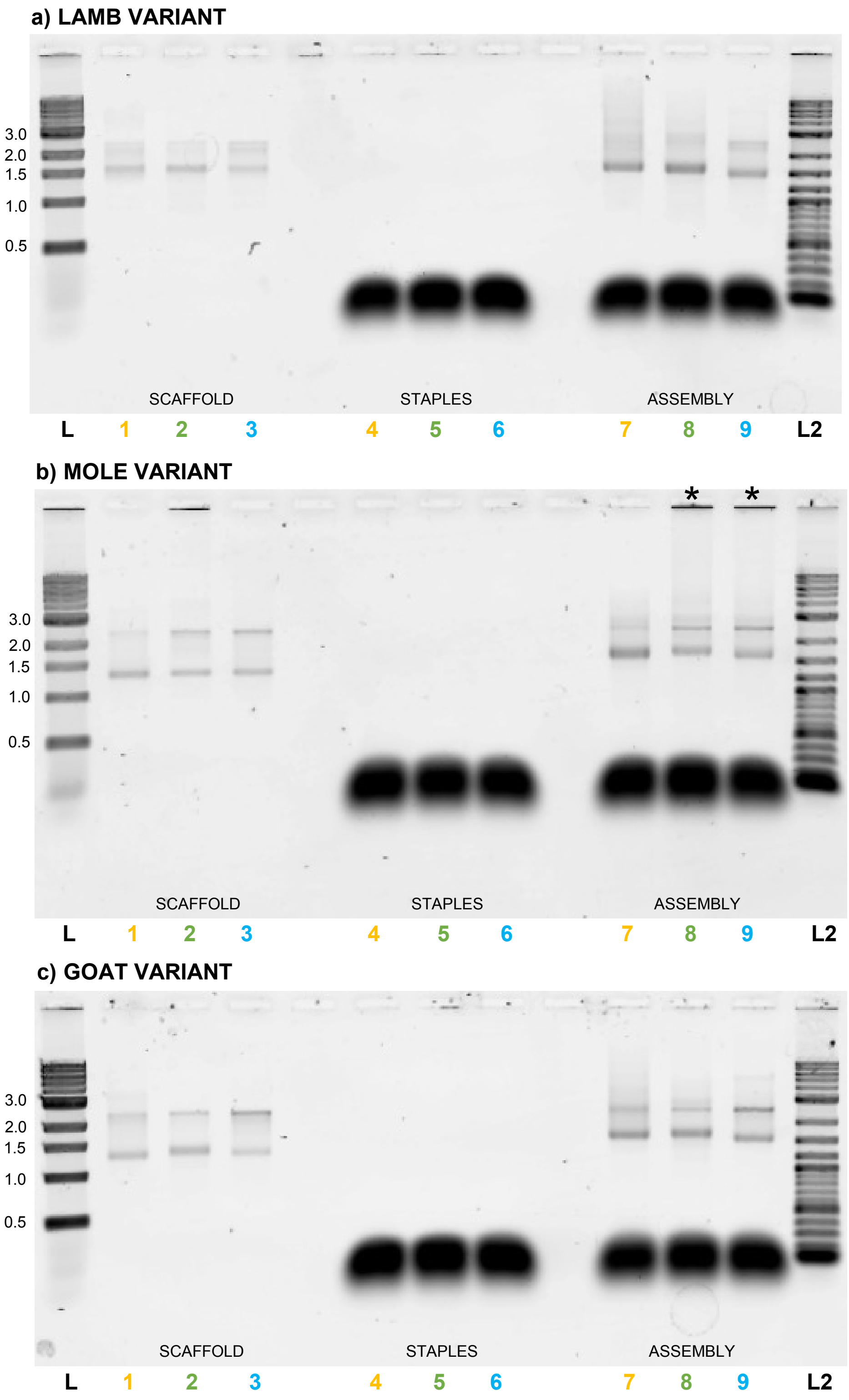


Supplementary Figure 29: Rectangle DNA origami variants (Lamb, Mole, Goat) and gel electrophoresis pre-screening. Laser-scanned images of SYBR® Gold-stained 1% TBE agarose gels loaded with scaffolds, staple strands mixes and assembly products obtained from isothermal annealing without initial denaturation (numbers in orange), isothermal annealing with initial denaturation (numbers in green) and fast annealing ramp (numbers in blue). a) Lamb variant; b) Mole variant; c) Goat variant; L: 1 kb Ladder; L2: 1 kb Plus Ladder; *: aggregates in the loading well. Molecular sizes in kilobases are indicated.

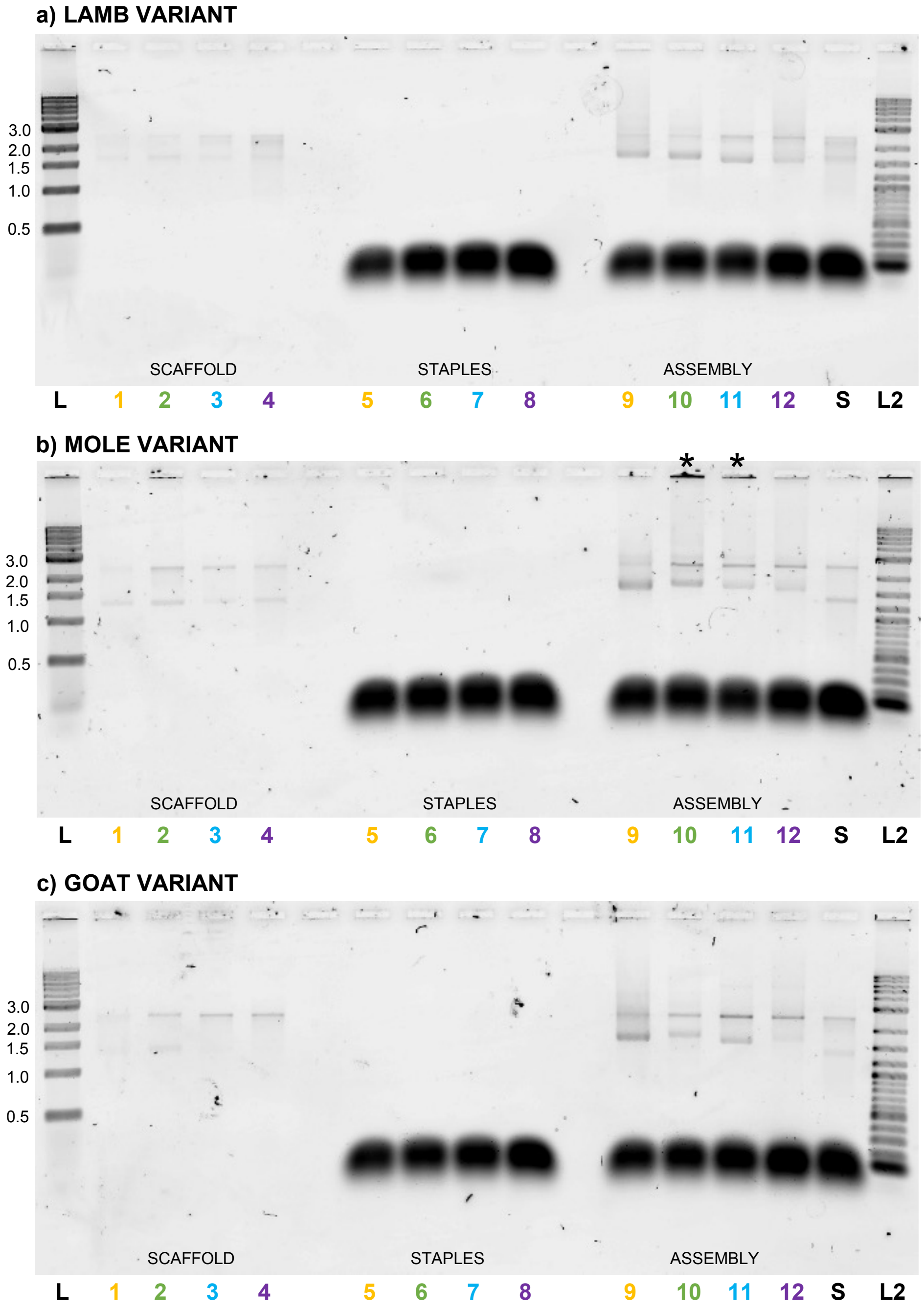


Supplementary Figure 30: Rectangle DNA origami variants (Lamb, Mole, Goat) and gel electrophoresis pre-screening. Laser-scanned images of SYBR® Gold-stained 1% TBE agarose gels loaded with scaffolds, staple strands mixes and assembly products obtained from isothermal annealing without initial denaturation (numbers in orange), isothermal annealing with initial denaturation (numbers in green), fast annealing ramp (numbers in blue) and slow annealing ramp (numbers in purple). Samples annealed considering the first 3 temperature ramps (as shown in S29) were kept at 4 °C and run again with samples annealed the day after with a slower ramp. a) Lamb variant; b) Mole variant; c) Goat variant; L: 1 kb Ladder; L2: 1 kb Plus Ladder; S: Lamb, Mole and Goat scaffolds and non-complementary Goat, Lamb and Mole staple strands set (negative control); *: aggregates in the loading well. Molecular sizes in kilobases are indicated.

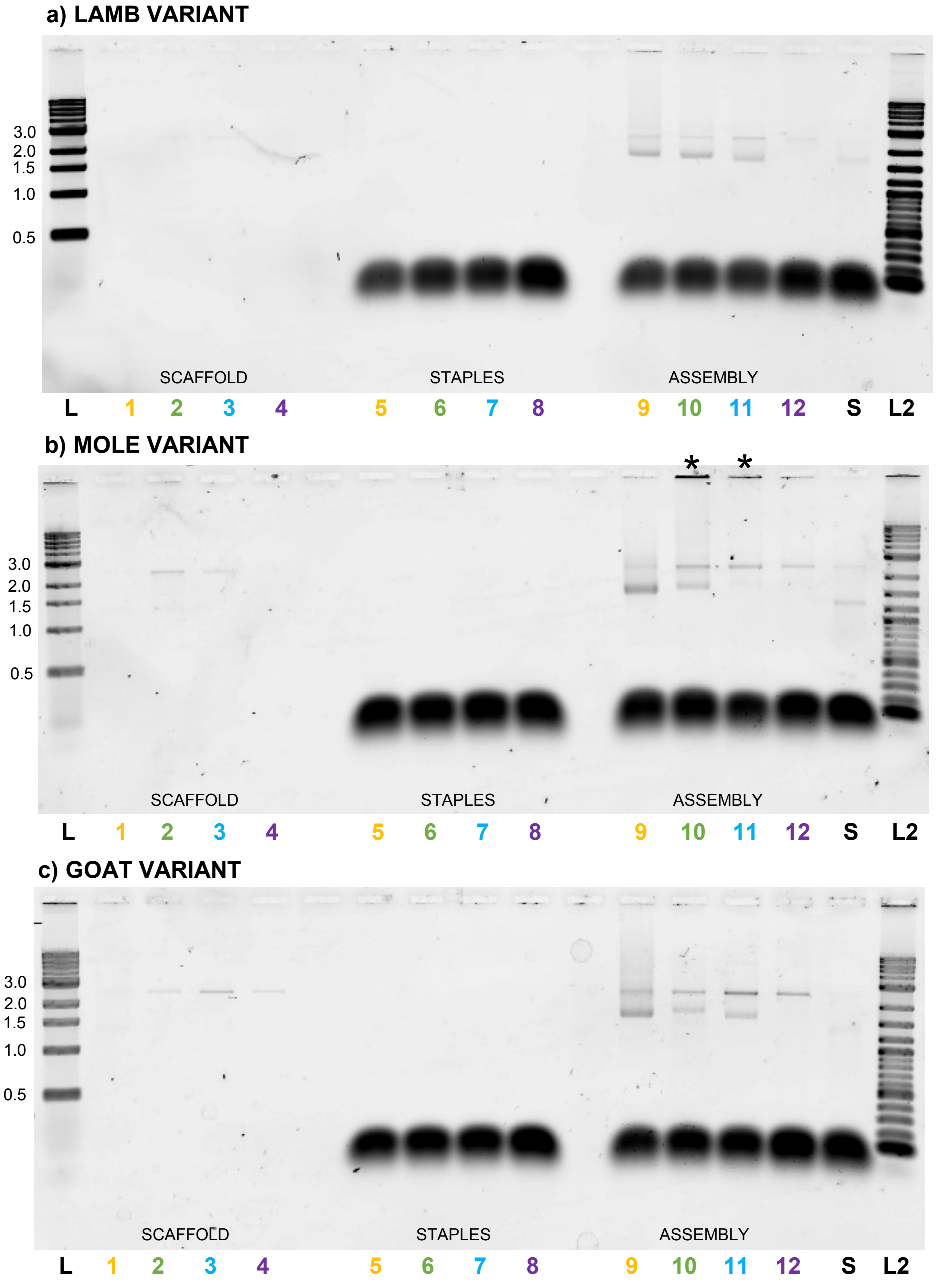


Supplementary Figure 31: Rectangle DNA origami variants (Lamb, Mole, Goat) and gel electrophoresis pre-screening. Laser-scanned images of SYBR® Gold-stained 1% TBE agarose gels loaded with scaffolds, staple strands mixes and assembly products obtained from isothermal annealing without initial denaturation (numbers in orange), isothermal annealing with initial denaturation (numbers in green), fast annealing ramp (numbers in blue) and slow annealing ramp (numbers in purple). Samples annealed considering 4 different temperature conditions (as shown in S29 and S30) were kept at 4 °C and run again after 4 (slow ramp) or 5 days (isothermal foldings and fast ramp). a) Lamb variant; b) Mole variant; c) Goat variant; L: 1 kb Ladder; L2: 1 kb Plus Ladder; S: Lamb, Mole and Goat scaffolds and non-complementary Goat, Lamb and Mole staple strands set (negative control); *: aggregates in the loading well. Molecular sizes in kilobases are indicated.

### Supplementary Note 14 Independent Replication of Results at University of Bonn

This section details a replication of results carried out at the University of Bonn, LIMES, Germany (Famulok research group).

| NEWCASTLE UNIVERSITY | | | | |
|---|---|---|---|---|
| Rectangle variants | Length (nm) | Width (nm) | Triangle variants | Side (nm) |
| Goat | 63±3 | 55±3 | Deer | 66±0.51 |
| Lamb | 63±1 | 51±2 | Lion | 63±0.68 |
| **UNIVERSITY OF BONN** | | | | |
| Rectangle variants | Length (nm) | Width (nm) | Triangle variants | Side (nm) |
| Goat | 53±6 | 44±4 | Deer | 53±4 |
| Lamb | 58±6 | 44±4 | Lion | 56±3 |

Supplementary Table 6: Average length and width of each variant measured from AFM images at Newcastle University and University of Bonn (mean ± standard deviation, n = 25). The triangle side expected dimension is about 55 nm. The rectangle length and width expected values are 62 nm and 47 nm, respectively.

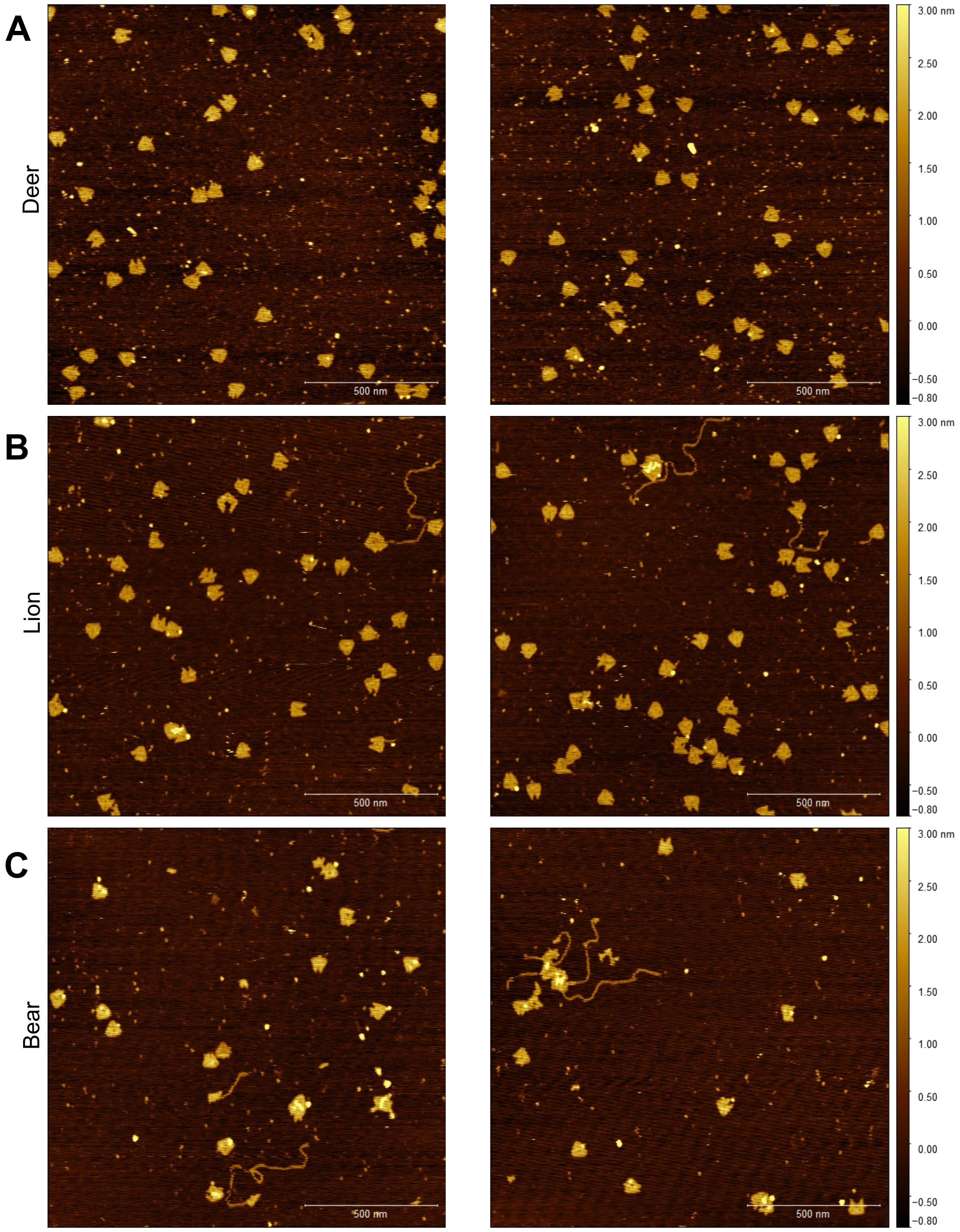


Supplementary Figure 32: AFM images of purified triangle DNA origami variants assembled using a fast temperature ramp (samples prepared and imaged at University of Bonn, LIMES, Famulok research group). High resolution AFM images of three triangle DNA origami variants Deer (A), Lion (B) and Bear (C). The respective purified assemblies were deposited on the Mica passivated with poly-L-ornithine and the images were obtained in AC mode in liquid. Scale bars: 500 nm.

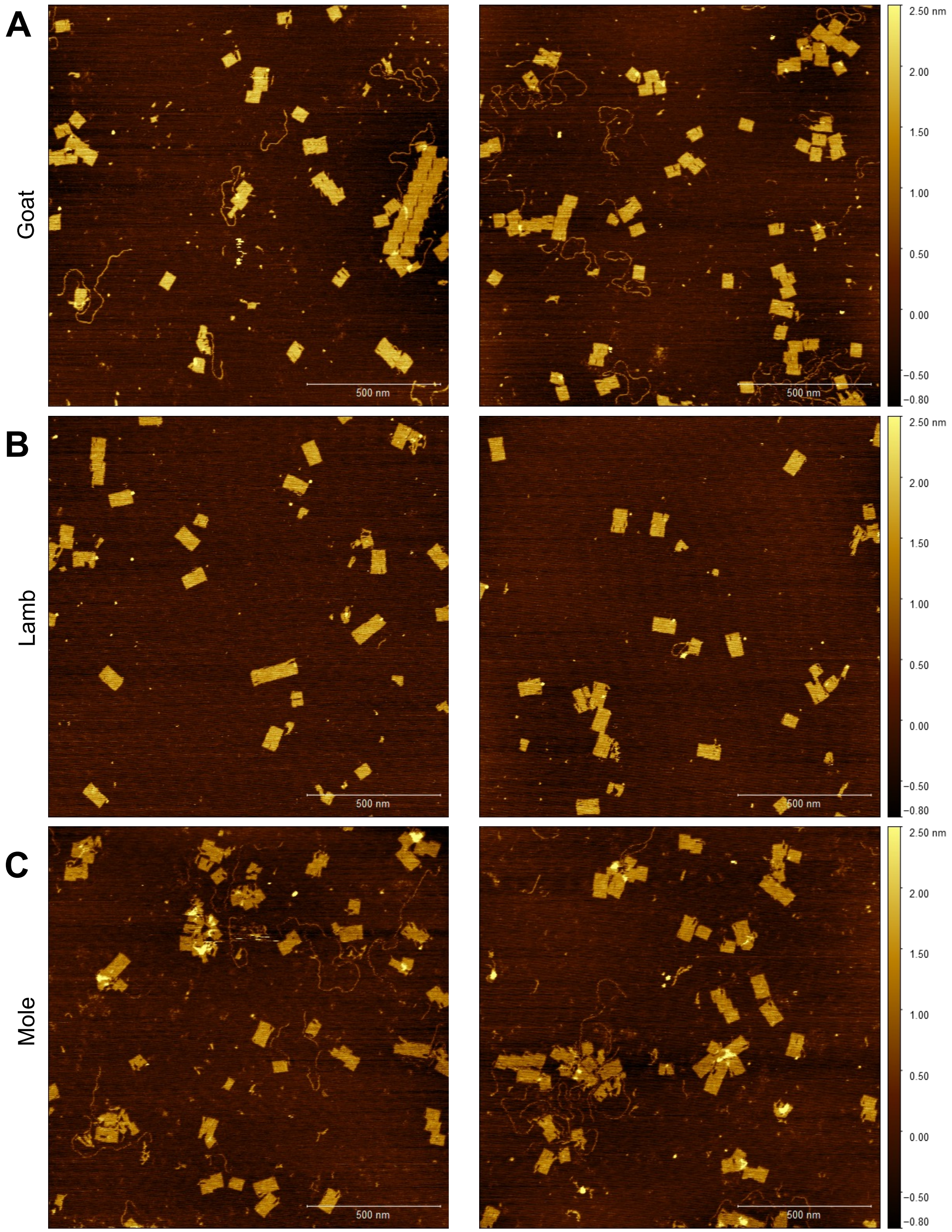


Supplementary Figure 33: AFM images of purified rectangle DNA origami variants assembled using a fast temperature ramp (samples prepared and imaged at University of Bonn, LIMES, Famulok research group). High resolution AFM images of three rectangle DNA origami variants Goat (A), Lamb (B) and Mole (C). The respective purified assemblies were deposited on the Mica passivated with poly-L-ornithine and the images were obtained in AC mode in liquid. Scale bars: 500 nm.

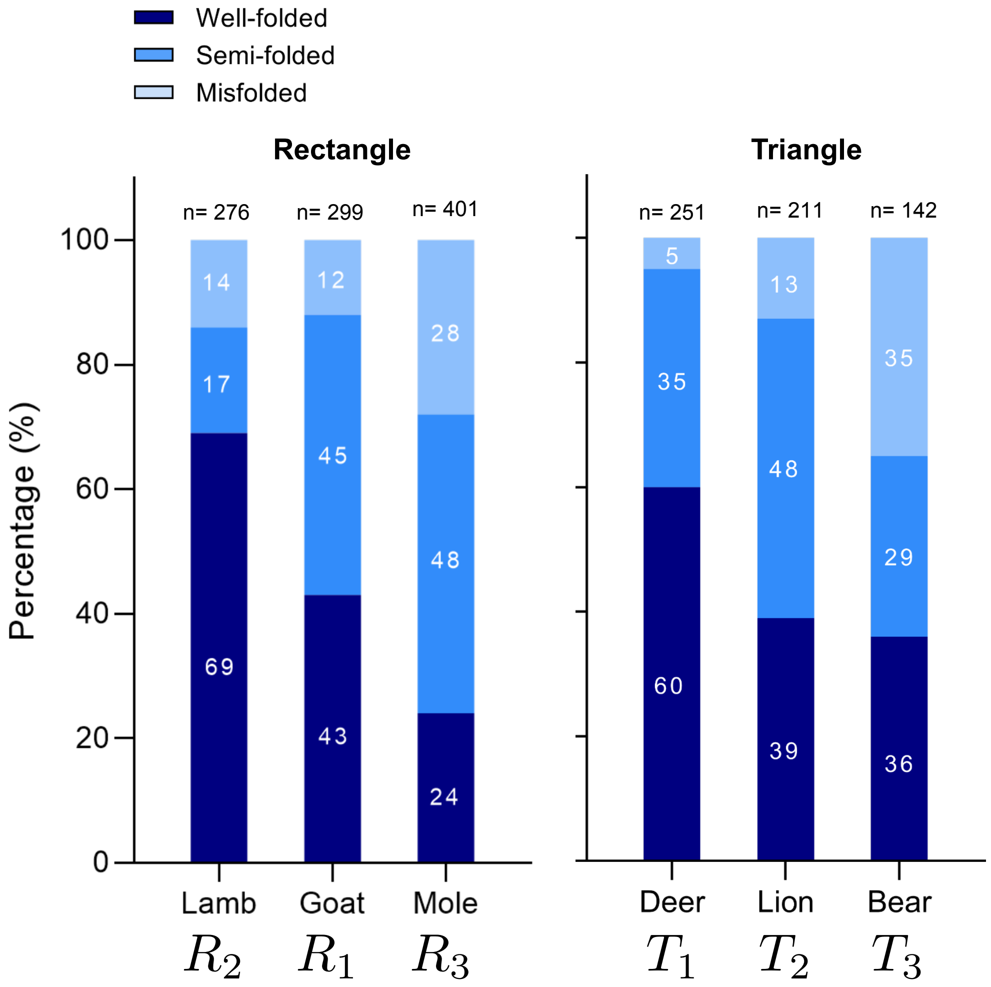


Supplementary Figure 34: Stacked bar charts of DNA origami variant populations assembled using a fast temperature ramp and imaged by AFM (samples prepared and imaged at University of Bonn, LIMES, Famulok research group).

### Supplementary Note 15 Optical Tweezers

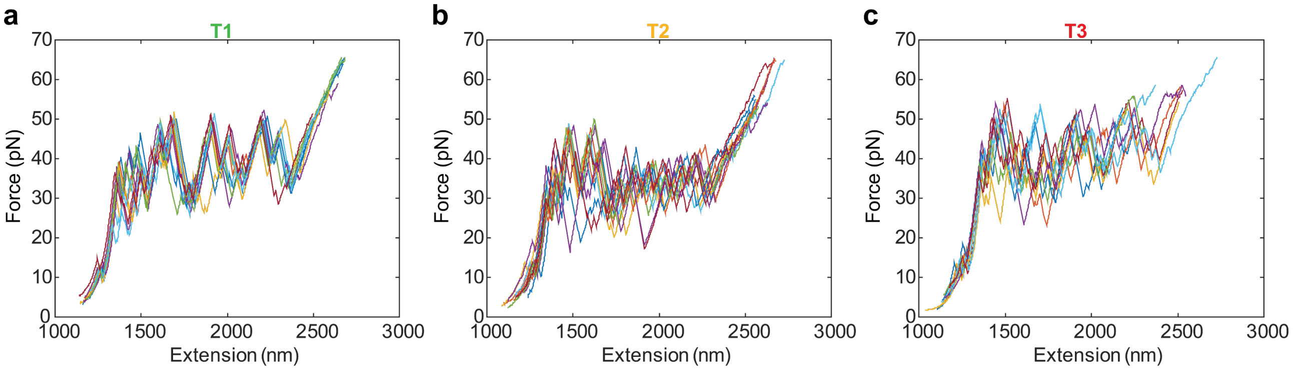


Supplementary Figure 35: Force vs. extension curves for each of the triangle origami variants. Ten curves exceeding 50% of the scaffold length were randomly selected for each variant: (a) T1, (b) T2, and (c) T3.

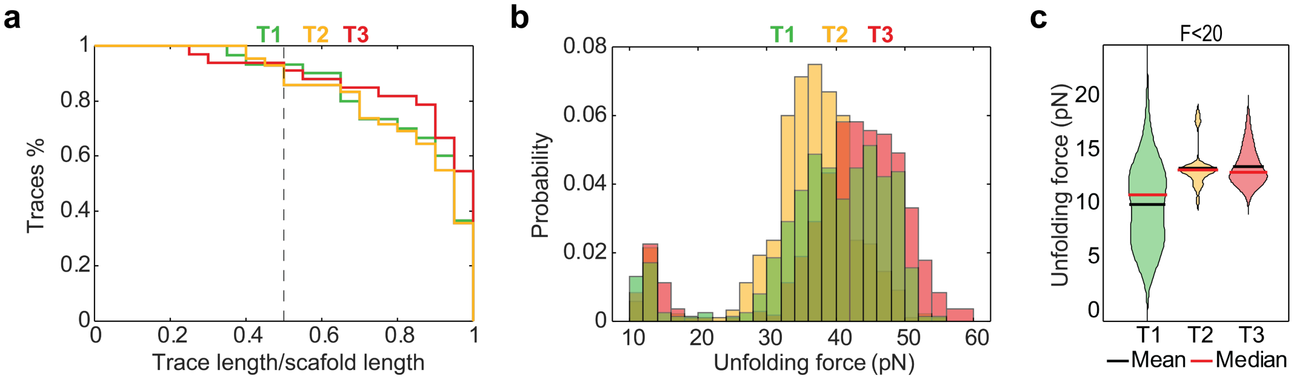


Supplementary Figure 36: (a) Distribution of the length of the experimental traces. Shown is the fraction of experimental traces for each triangle variant that reached a relative trace length. The dashed line indicates the 50% relative length. (b) Distribution of total unfolding forces for each triangle origami variant, represented in different colours. $n_{T1} = 379$, $n_{T2} = 441$, $n_{T3} = 531$ are the number of unfolding events. (c) Violin plot of unfolding forces below 20 pN, derived from all unfolding curves for each variant ($n_{T1} = 30$, $n_{T2} = 33$, $n_{T3} = 42$ are the number of measured traces).

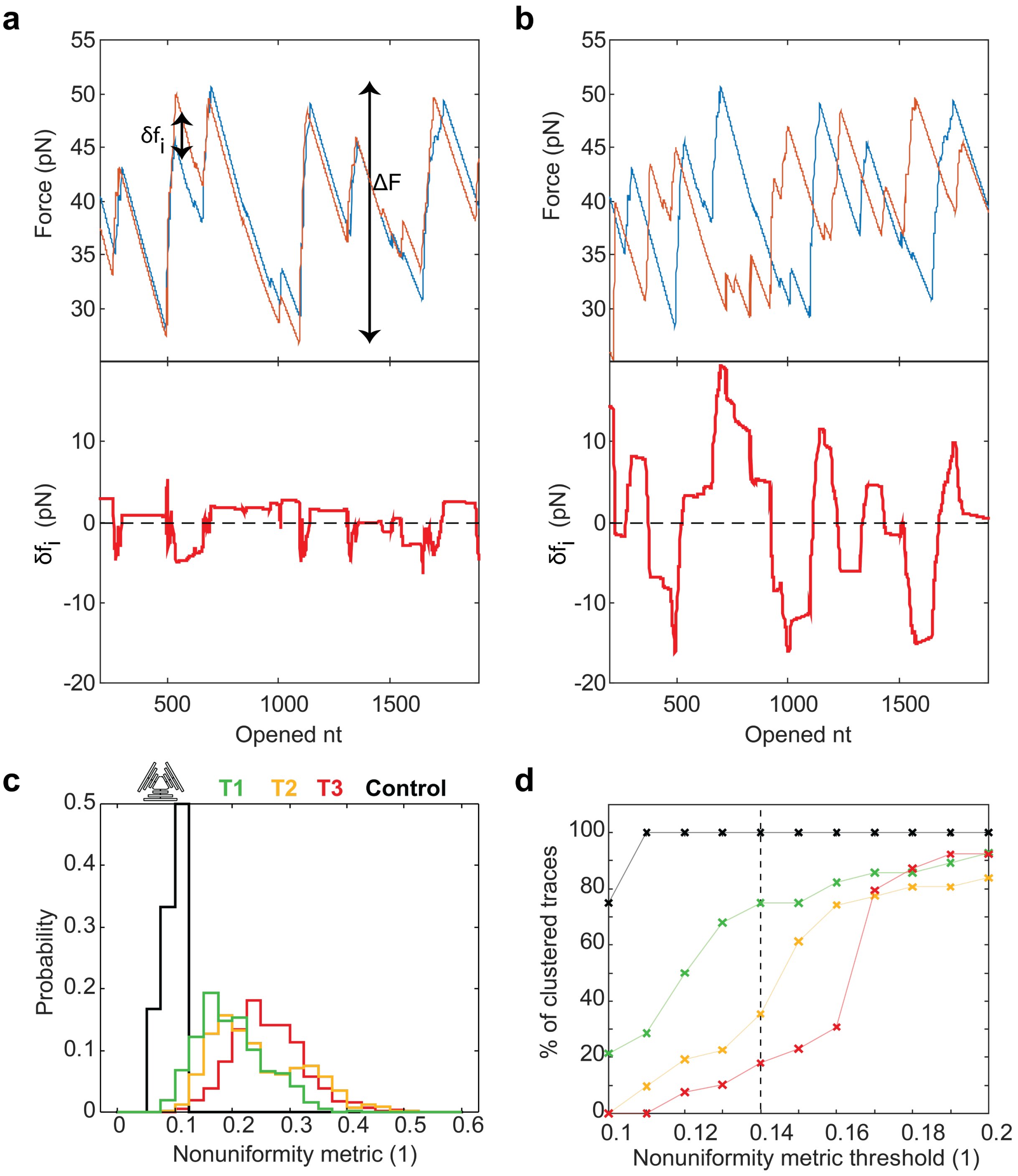


Supplementary Figure 37: (a,b) Top panel: Force vs. number of opened nucleotides (nt) for two unfolding curves. The variable $\delta f_i$ denotes the force difference at a specific point along the opened nt axis, while $\Delta F$ represents the mean peak-to-peak force difference calculated for a specific variant. Bottom panel: The force difference between the two curves from the top panel, shown for (a) similar unfolding curves and (b) non-similar unfolding curves. (c) Histogram of the non-uniformity metric for different triangle origami structure groups, each shown in different colours. The control group for DNA unzipping curves is shown in black. ($n_{T1} = 756$, $n_{T2} = 930$, $n_{T3} = 1482$). (d) The percentage of clustered uniform traces is shown at various NUM thresholds for each variant of the triangle origami structures and the control, represented in different colours. The dashed black line indicates the threshold used in Figure 6 of the main paper.

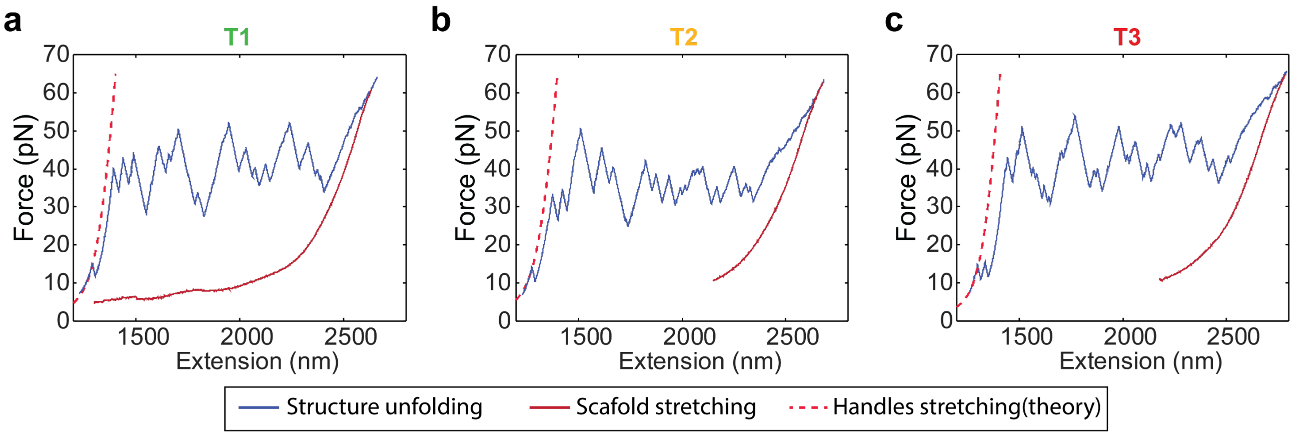


Supplementary Figure 38: Representative force vs. extension curves for triangle origami structure unfolding (blue). Shown in red are the corresponding unfolding curves obtained after subsequent relaxation and stretching again, indicating complete structure disassembly. The dashed line shows the theoretical force vs. extension curve for stretching the handles, as calculated from an Extensible Worm Like Chain model. (a) T1, (b) T2, and (c) T3.

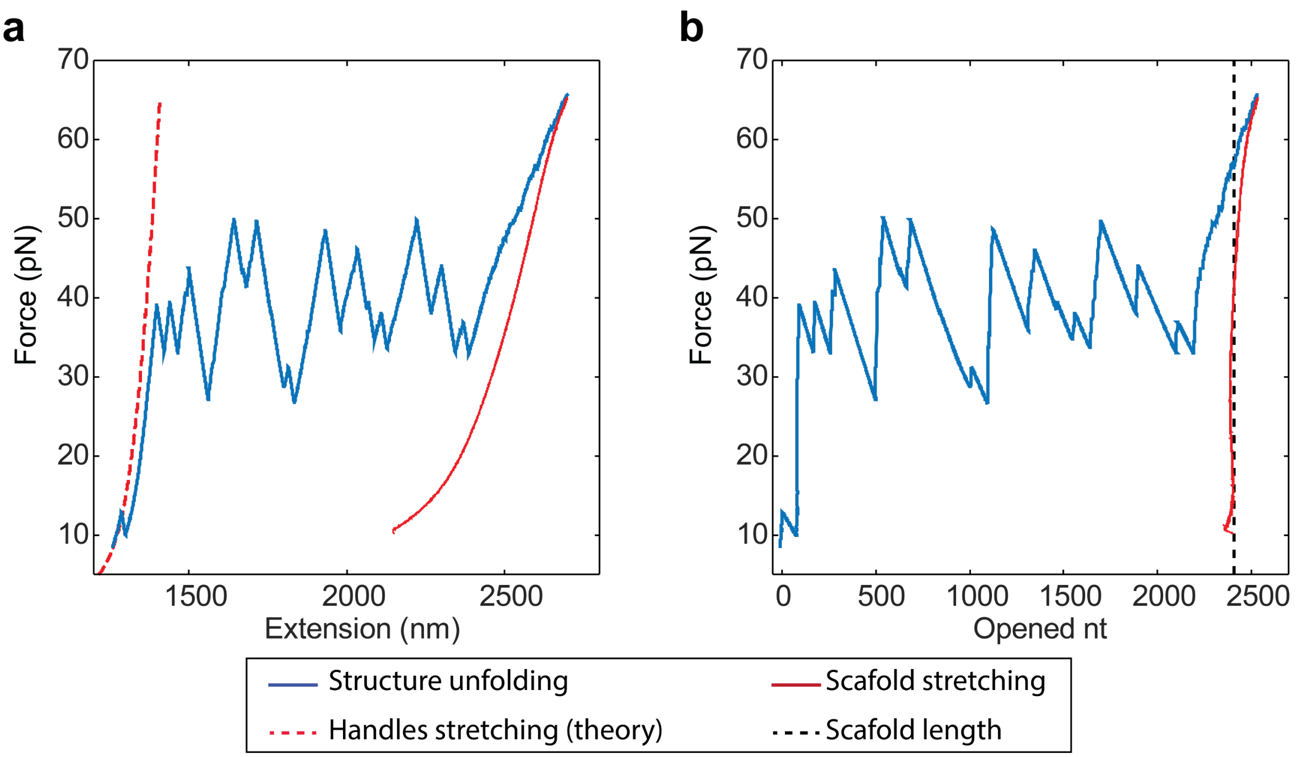


Supplementary Figure 39: (a) A representative force vs. extension curve, and (b) its corresponding force vs. opened nucleotides (nt) curve for a T1 triangle structure. In both (a) and (b), the unfolding of the origami structure is shown in blue, and the scaffold unfolding is represented by a solid red line. In (a), the theoretical stretching curve for the dsDNA handles is shown as a dashed red line, while in (b), the length of the scaffold is indicated by a dashed black line.

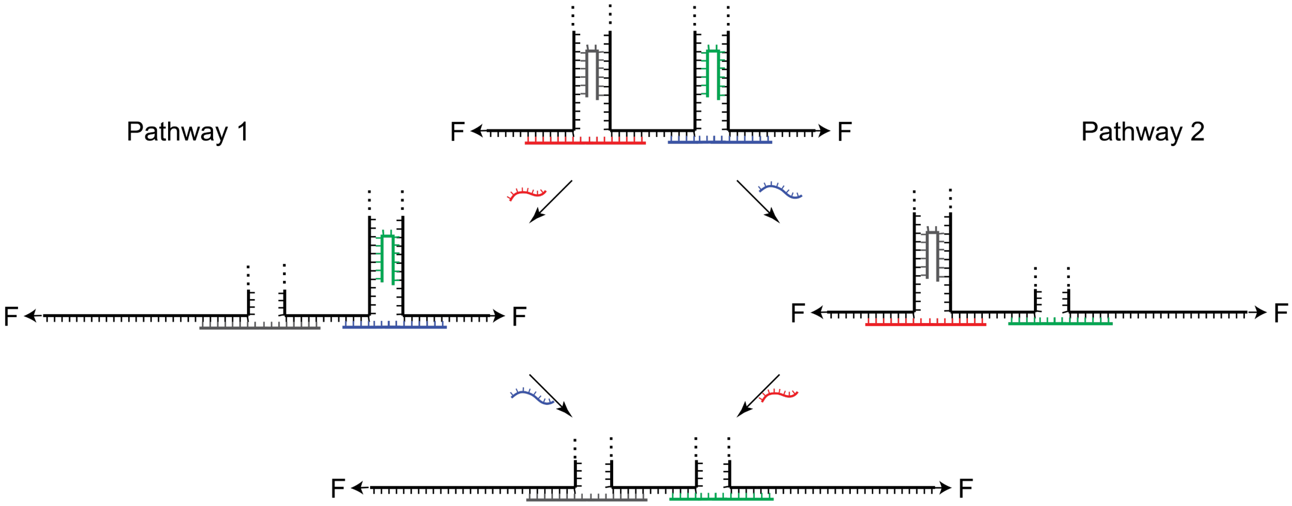


Supplementary Figure 40: Schematic representation of alternative pathways in the force-unfolding of an origami.

| | Experiments | Pairs | Unfolding events | Unfolding events F>20 | Unfolding events F<20 |
|---|---|---|---|---|---|
| T1 | 28 | 756 | 379 | 328 | 51 |
| T2 | 31 | 930 | 441 | 414 | 27 |
| T3 | 39 | 1482 | 531 | 481 | 50 |
| Control | 4 | 12 | | | |

Supplementary Table 7: Optical tweezers data collection statistics.

| Variant 1 | Variant 2 | p-value (F>20) | p-value (F<20) |
|---|---|---|---|
| T1 | T2 | 1.15E-15 | 8.55E-06 |
| T1 | T3 | 5.87E-12 | 2.36E-09 |
| T2 | T3 | 3.36E-59 | 0.72 |

Supplementary Table 8: p-value of the unfolding force between different triangle variants obtained using a Student's t-test.

| Variant 1 | Variant 2 | p-value |
|---|---|---|
| T1 | T2 | 1.23E-13 |
| T1 | T3 | 2.04E-45 |
| T1 | Control | 2.23E-11 |
| T2 | T3 | 1.19E-08 |
| T2 | Control | 2.36E-12 |
| T3 | Control | 1.28E-18 |

Supplementary Table 9: p-value of the unfolding non-uniformity metric between different variants, as well as the control group, obtained using a Student's t-test.

| | $\Delta F$ (pN) $\pm$ s.e.m. |
|---|---|
| T1 | 25.4$\pm$0.9 |
| T2 | 23.2$\pm$0.6 |
| T3 | 24.5$\pm$0.6 |
| Control | 4.7$\pm$0.2 |

Supplementary Table 10: The mean peak-to-peak force difference for each variant.

| Description | Sequence |
|---|---|
| Forward primer for handle 1 with Biotin tag | /5BioTinTEG/GATCTCCAGCCAGGAACTATTGA |
| Reverse primer handle 1 with NcoI site | CACCCATGGTTTCGACCTGCTCTTCAGCA |
| Forward primer for handle 2 with Bgl1 site | ATGGCCTAGACGGCGAGCCTGGGTTTATAAGGGGAGCGGTGA |
| Reverse primer for handle 2 with Bgl1 site | CAGCCTGCATGGCAAGGACCAGCGTTTTGTTGAAA |
| Double Dig oligo 1 with digoxigenin tag | /5Phos/ ACAGGGTGGTCCCGGCACCT /3Dig_N/ |
| Double Dig oligo 2 with digoxigenin tag | /5Phos/AC AGG GTG GTC CCG GCA CCT /3Dig_N/ |

Supplementary Table 11: Primers and oligos used to construct the two handles for the optical tweezers experiments.

#### Supplementary Note 16 AFM Examples of “Semi-Folded” and “Mis-Folded” Origamis

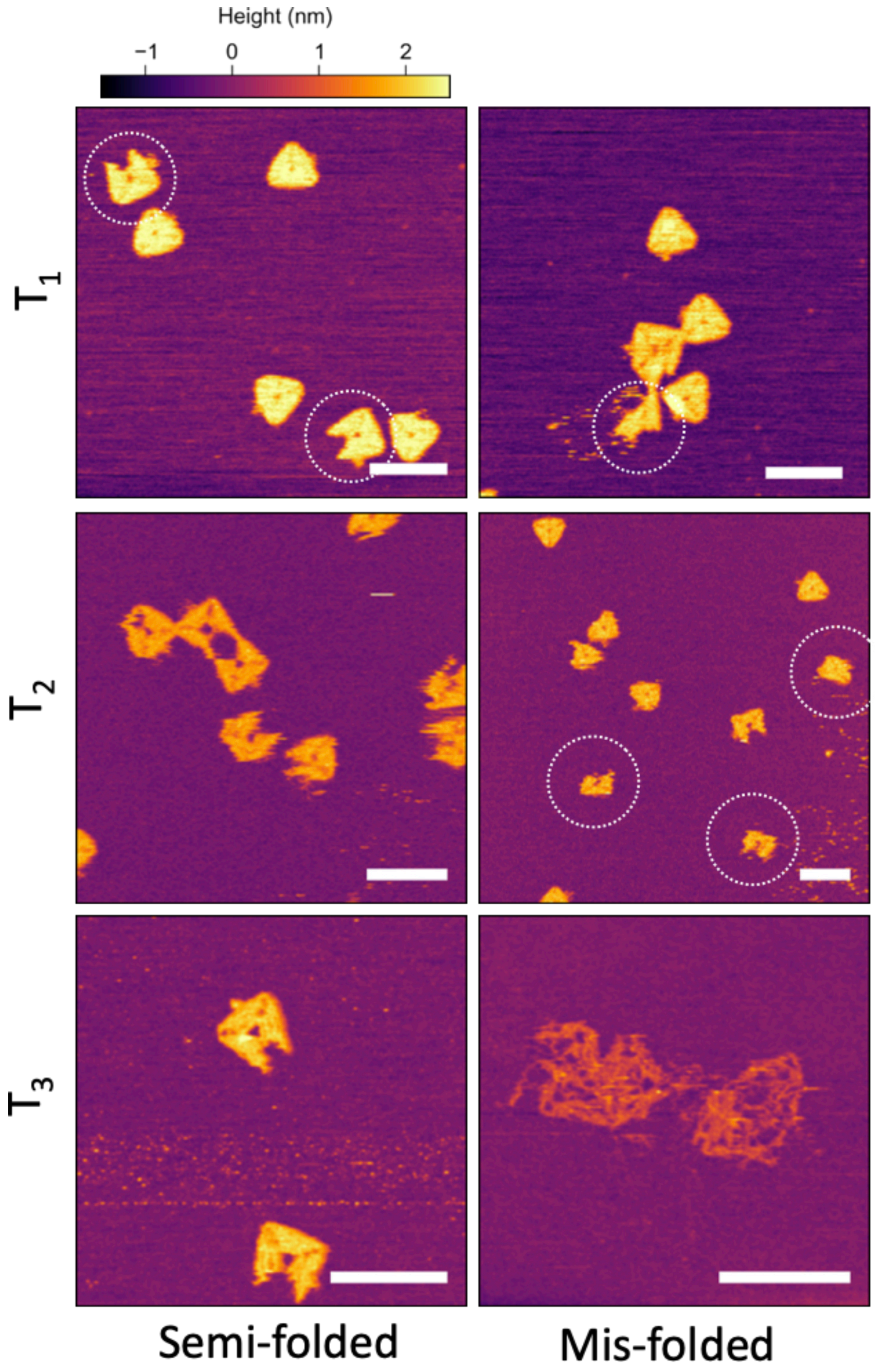


Supplementary Figure 41: Triangle origami. Representative high-resolution AFM images of semi-folded (left column) and mis-folded (right column) categories. If there are a mix of triangles folded in different degrees present, circles are used to denote triangles in the folding state corresponding to the column. All colour scales run from -1.5 to 2.5 nm (4.0 nm range; 0.5 nm offset) and all length scale bars are 100 nm.

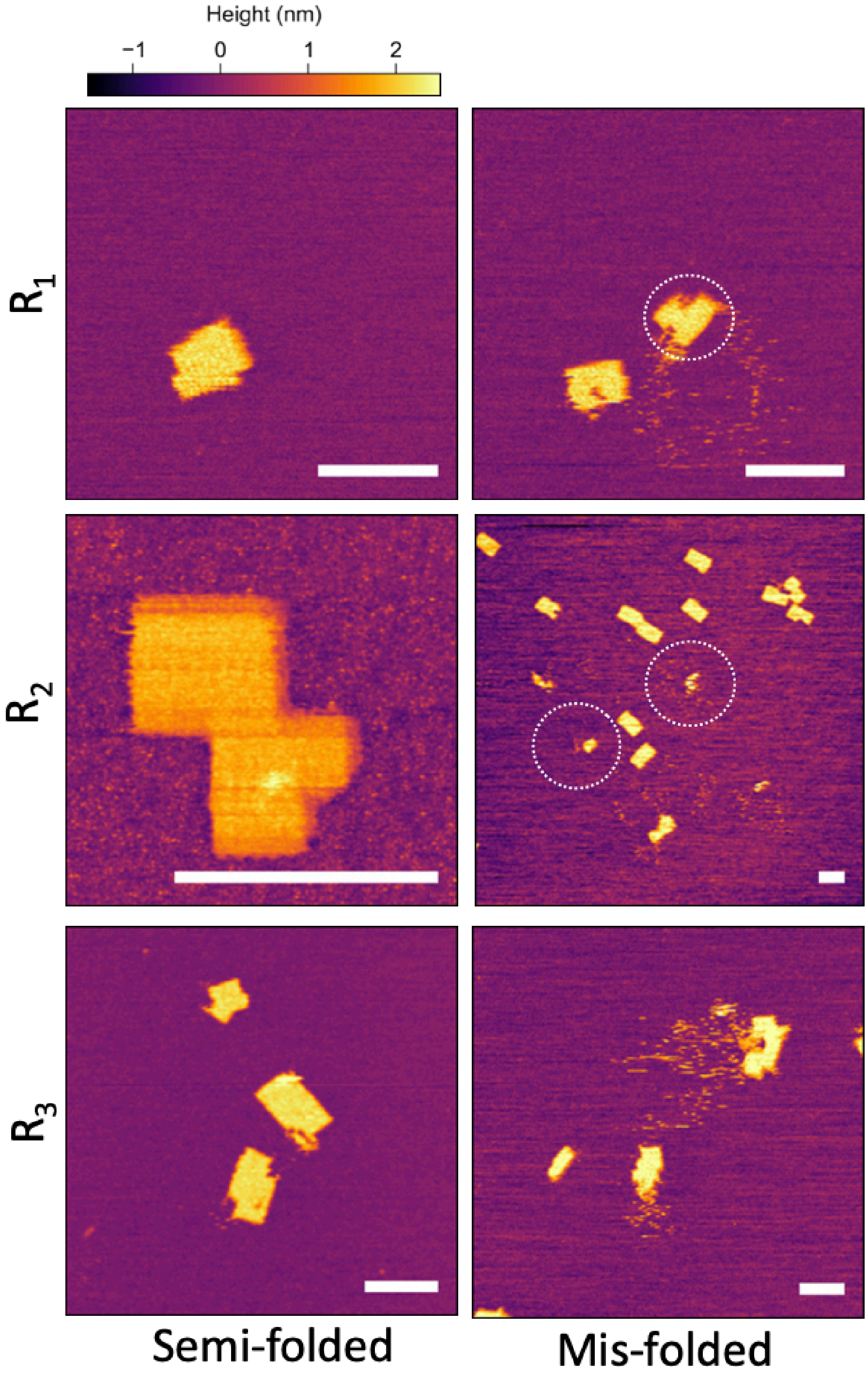


Supplementary Figure 42: Rectangle origami. Representative high-resolution AFM images of semi-folded (left column) and mis-folded (right column) categories. If there are a mix of rectangles folded in different degrees present, circles are used to denote rectangles in the folding state corresponding to the column. All colour scales run from -1.5 to 2.5 nm (4.0 nm range; 0.5 nm offset) and all length scale bars are 100 nm.
